## Supplemental Information 1 for "Targeted proteomics as a tool to detect SARS-CoV-2 proteins in clinical specimens"

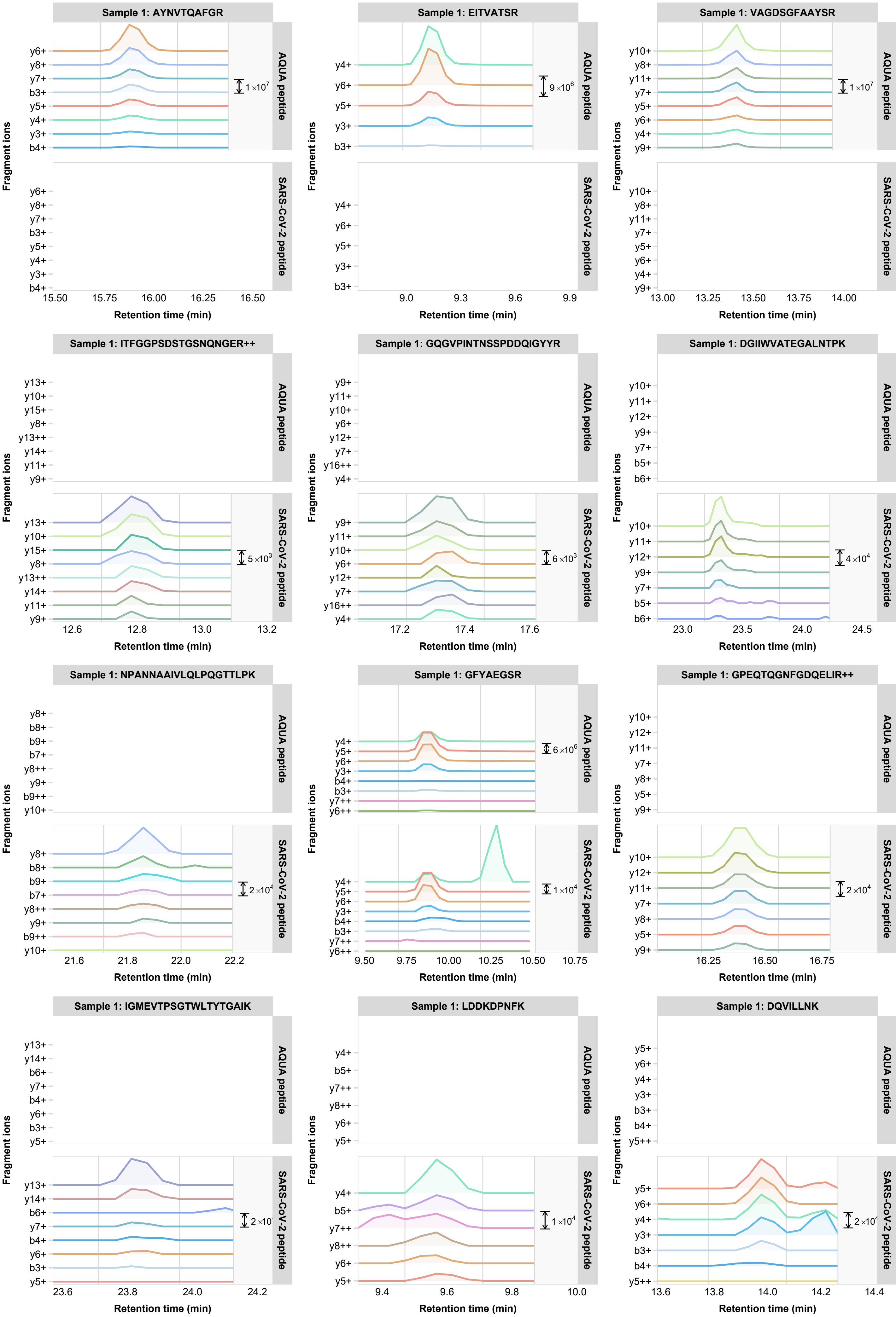
Sample 1: AYNVTQAFGR
AQUA peptide
SARS-CoV-2 peptide
y6+ y8+ y7+ b3+ y5+ y4+ y3+ b4+
1×10^7
15.50 15.75 16.00 16.25 16.50
Sample 1: EITVATSR
y4+ y6+ y5+ y3+ b3+
9×10^6
9.0 9.3 9.6 9.9
Sample 1: VAGDSGFAAYSR
y10+ y8+ y11+ y7+ y5+ y6+ y4+ y9+
1×10^7
13.00 13.25 13.50 13.75 14.00
Sample 1: ITFGGPSDSTGSNQNGER++
y13+ y10+ y15+ y8+ y13++ y14+ y11+ y9+
5×10^3
12.6 12.8 13.0 13.2
Sample 1: GQGVPINTNSSPDDQIGYYR
y9+ y11+ y10+ y6+ y12+ y7+ y16++ y4+
6×10^3
17.2 17.4 17.6
Sample 1: DGIIWVATEGALNTPK
y10+ y11+ y12+ y9+ y7+ b5+ b6+
4×10^4
23.0 23.5 24.0 24.5
Sample 1: NPANNAAIVLQLPQGTTLPK
y8+ b8+ b9+ b7+ y8++ y9+ b9++ y10+
2×10^4
21.6 21.8 22.0 22.2
Sample 1: GFYAEGSR
y4+ y5+ y6+ y3+ b4+ b3+ y7++ y6++
6×10^6
1×10^4
9.50 9.75 10.00 10.25 10.50 10.75
Sample 1: GPEQTQGNFGDQELIR++
y10+ y12+ y11+ y7+ y8+ y5+ y9+
2×10^4
16.25 16.50 16.75
Sample 1: IGMEVTPSGTWLTYTGAIK
y13+ y14+ b6+ y7+ b4+ y6+ b3+ y5+
23.6 23.8 24.0 24.2
Sample 1: LDDKDPNFK
y4+ b5+ y7++ y8++ y6+ y5+
1×10^4
9.4 9.6 9.8 10.0
Sample 1: DQVILLNK
y5+ y6+ y4+ y3+ b3+ b4+ y5++
13.6 13.8 14.0 14.2 14.4
Fragment ions
Retention time (min)

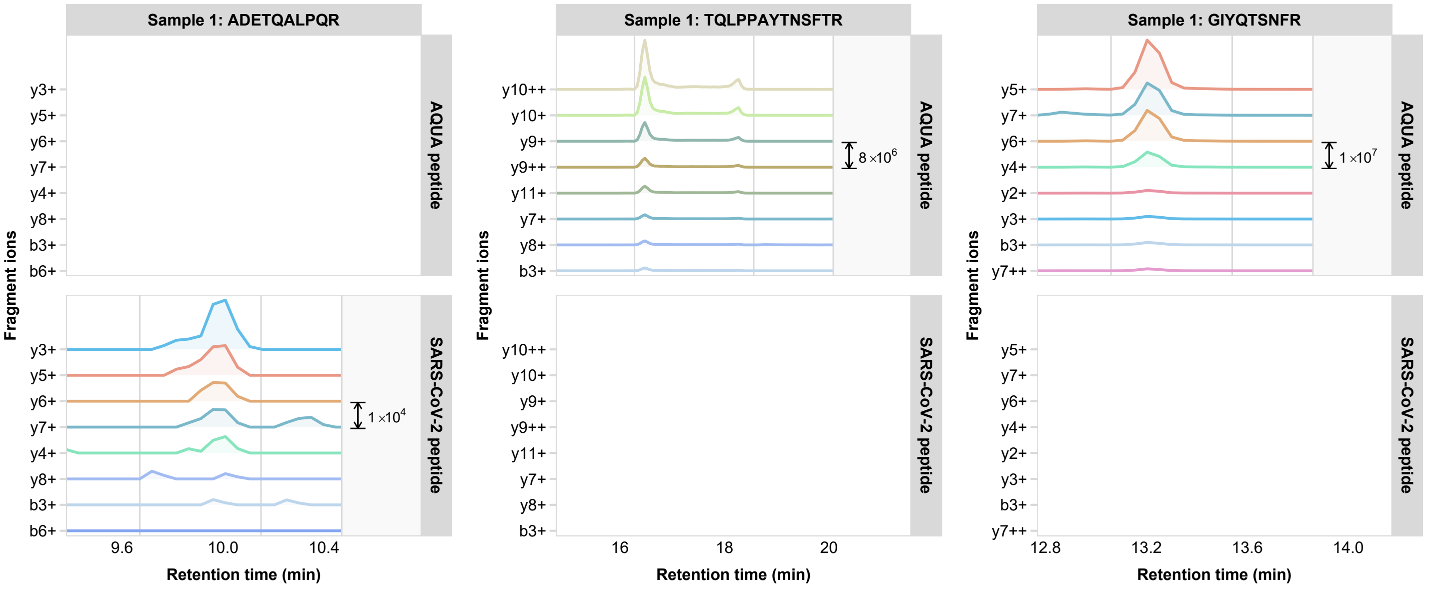

Sample 1: ADETQALPQR
Sample 1: TQLPPAYTNSFTR
Sample 1: GIYQTSNFR
AQUA peptide
SARS-CoV-2 peptide
Fragment ions
y3+
y5+
y6+
y7+
y4+
y8+
b3+
b6+
y10++
y10+
y9+
y9++
y11+
y2+
y7++
1 ×10^4
8 ×10^6
1 ×10^7
9.6
10.0
10.4
16
18
20
12.8
13.2
13.6
14.0
Retention time (min)

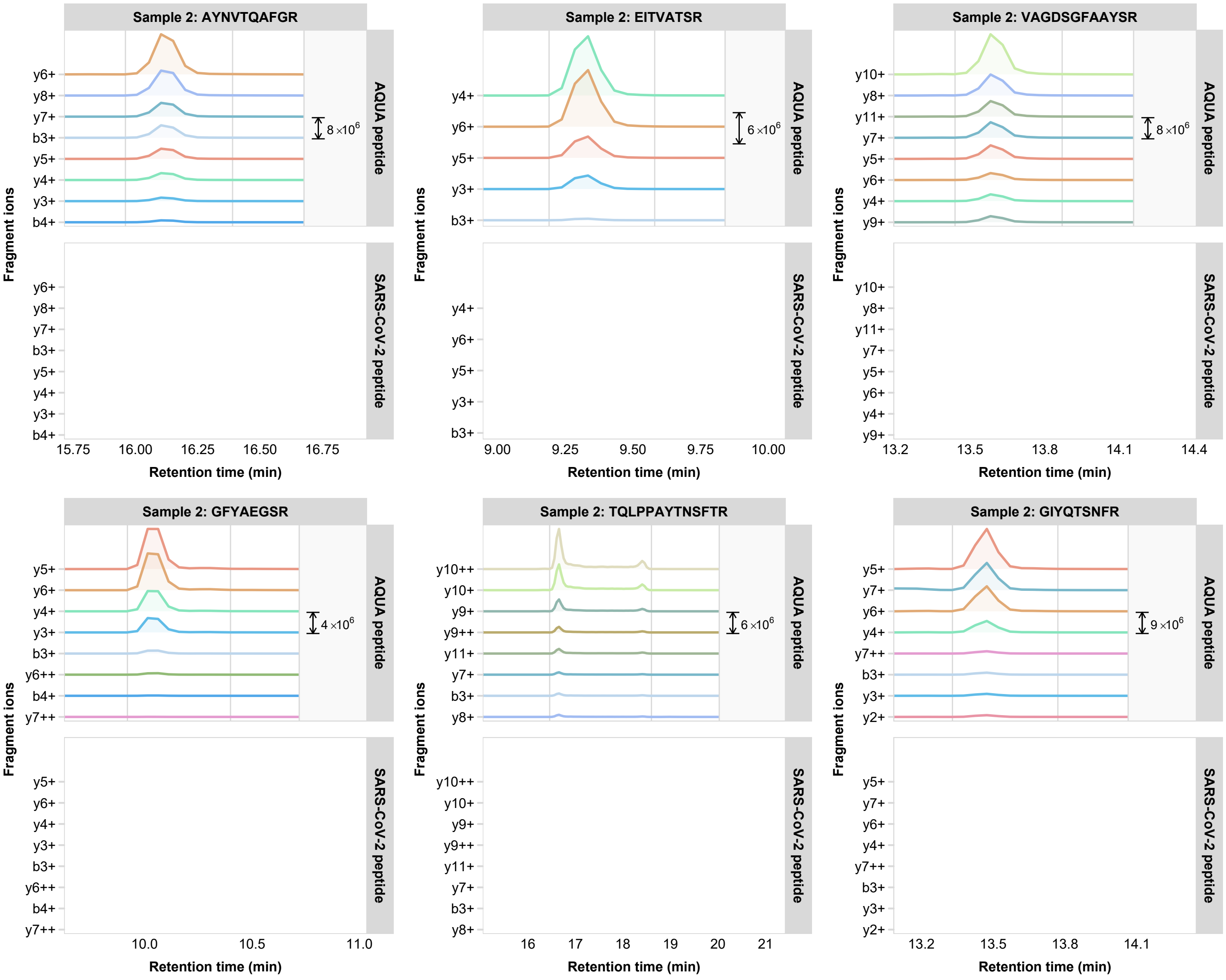
Sample 2: AYNVTQAFGR
Sample 2: EITVATSR
Sample 2: VAGDSGFAAYSR
Sample 2: GFYAEGSR
Sample 2: TQLPPAYTNSFTR
Sample 2: GIYQTSNFR
AQUA peptide
SARS-CoV-2 peptide
Fragment ions
Retention time (min)
8×10^6
6×10^6
8×10^6
4×10^6
6×10^6
9×10^6
y6+
y8+
y7+
b3+
y5+
y4+
y3+
b4+
y10+
y11+
y9+
y6++
y7++
y10++
y9++
y2+
15.75
16.00
16.25
16.50
16.75
9.00
9.25
9.50
9.75
10.00
13.2
13.5
13.8
14.1
14.4
10.0
10.5
11.0
16
17
18
19
20
21

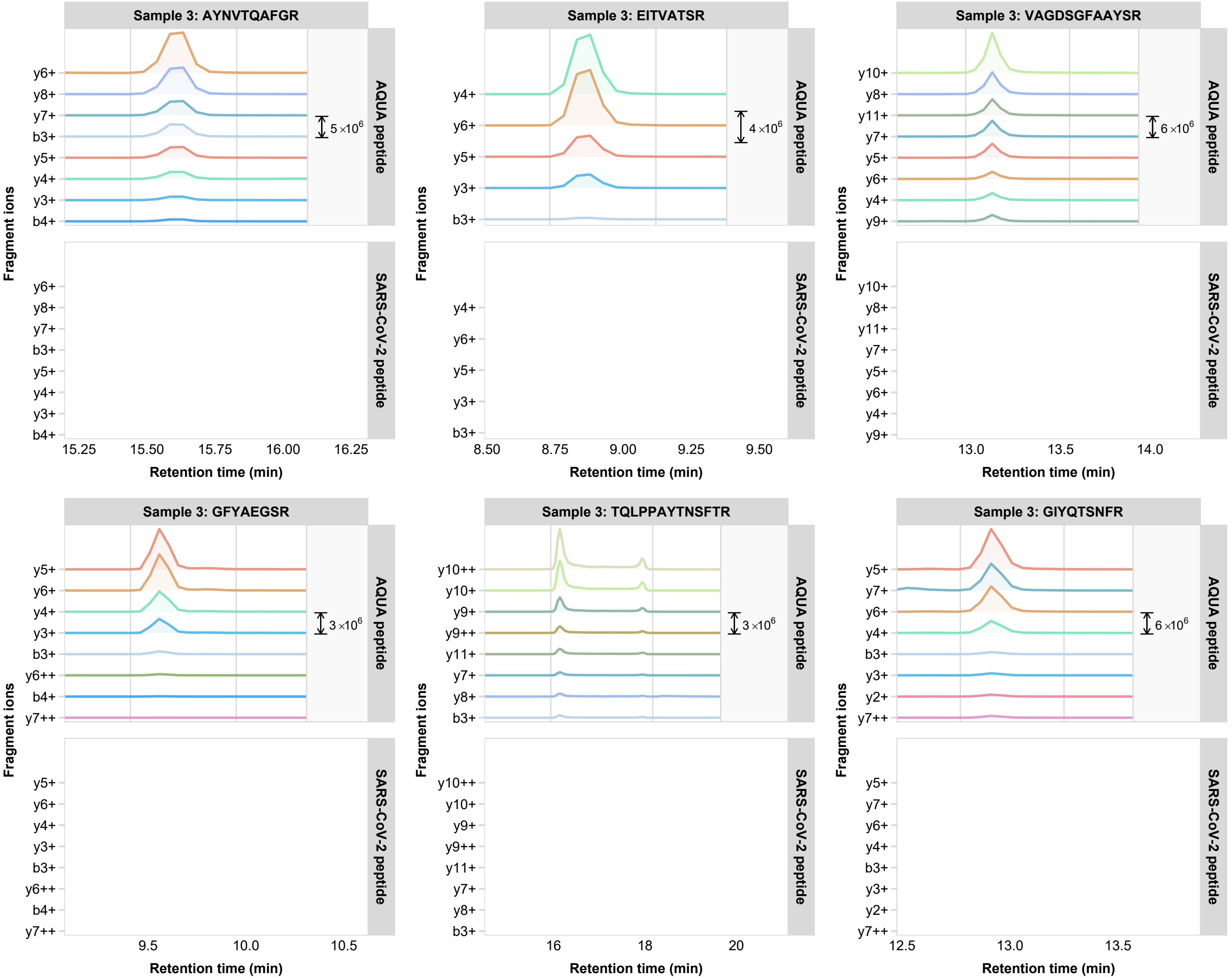

Sample 3: AYNVTQAFGR
AQUA peptide
SARS-CoV-2 peptide
5 ×10^6
y6+
y8+
y7+
b3+
y5+
y4+
y3+
b4+
15.25
15.50
15.75
16.00
16.25
Retention time (min)
Fragment ions
Sample 3: EITVATSR
AQUA peptide
SARS-CoV-2 peptide
4 ×10^6
y4+
y6+
y5+
y3+
b3+
8.50
8.75
9.00
9.25
9.50
Retention time (min)
Fragment ions
Sample 3: VAGDSGFAAYSR
AQUA peptide
SARS-CoV-2 peptide
6 ×10^6
y10+
y8+
y11+
y7+
y5+
y6+
y4+
y9+
13.0
13.5
14.0
Retention time (min)
Fragment ions
Sample 3: GFYAEGSR
AQUA peptide
SARS-CoV-2 peptide
3 ×10^6
y5+
y6+
y4+
y3+
b3+
y6++
b4+
y7++
9.5
10.0
10.5
Retention time (min)
Fragment ions
Sample 3: TQLPPAYTNSFTR
AQUA peptide
SARS-CoV-2 peptide
3 ×10^6
y10++
y10+
y9+
y9++
y11+
y7+
y8+
b3+
16
18
20
Retention time (min)
Fragment ions
Sample 3: GIYQTSNFR
AQUA peptide
SARS-CoV-2 peptide
6 ×10^6
y5+
y7+
y6+
y4+
b3+
y3+
y2+
y7++
12.5
13.0
13.5
Retention time (min)
Fragment ions

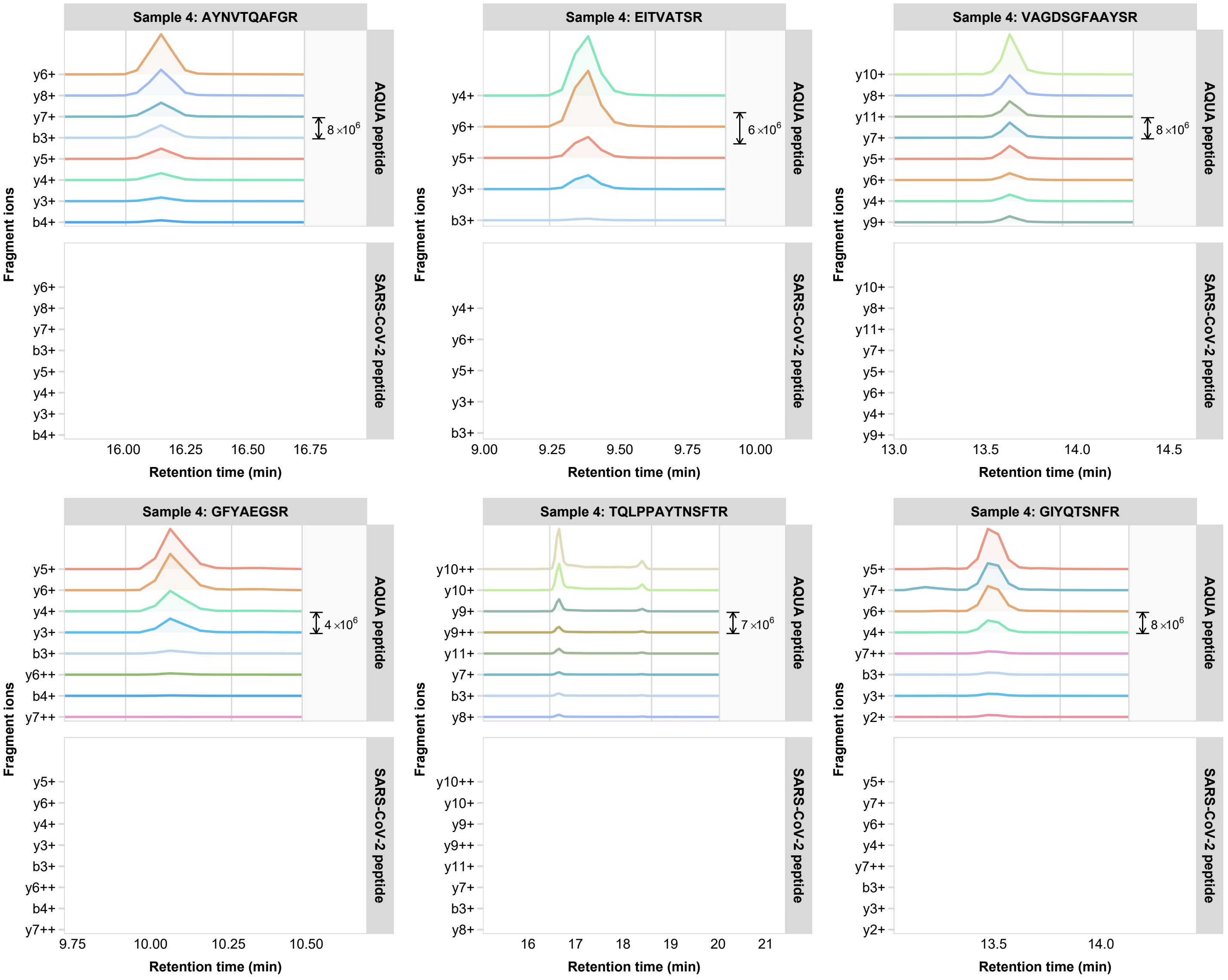

Sample 4: AYNVTQAFGR
Sample 4: EITVATSR
Sample 4: VAGDSGFAAYSR
Sample 4: GFYAEGSR
Sample 4: TQLPPAYTNSFTR
Sample 4: GIYQTSNFR
AQUA peptide
SARS-CoV-2 peptide
Fragment ions
Retention time (min)
8 ×10^6
6 ×10^6
4 ×10^6
7 ×10^6
y6+
y8+
y7+
b3+
y5+
y4+
y3+
b4+
y10+
y11+
y9+
y6++
y7++
y10++
y9++
y2+
y8+
16.00
16.25
16.50
16.75
9.00
9.25
9.50
9.75
10.00
13.0
13.5
14.0
14.5
9.75
10.00
10.25
10.50
16
17
18
19
20
21
13.5
14.0

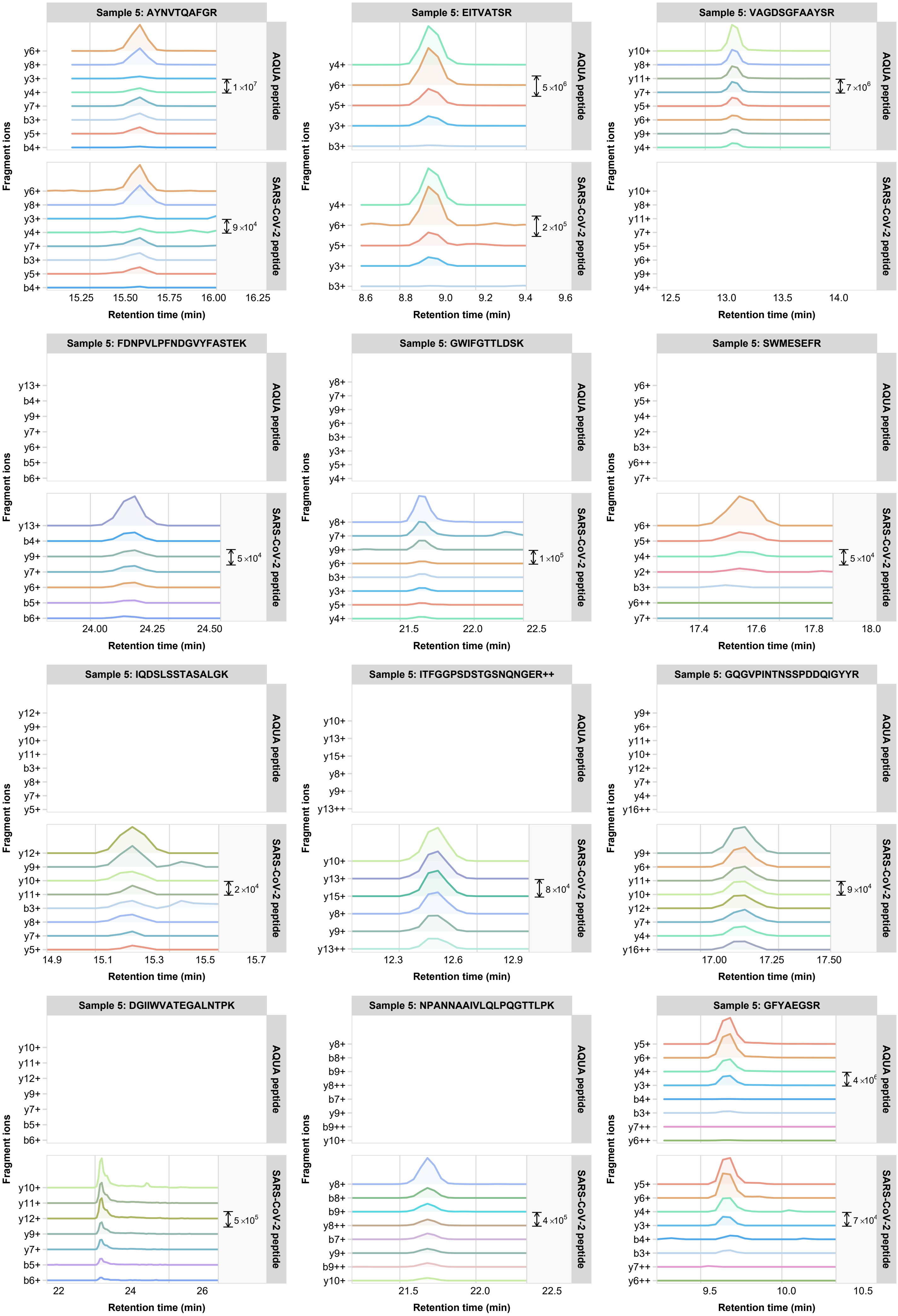

Sample 5: AYNVTQAFGR
Sample 5: EITVATSR
Sample 5: VAGDSGFAAYSR
Sample 5: FDNPVLPFNDGVYFASTEK
Sample 5: GWIFGTTLDSK
Sample 5: SWMESEFR
Sample 5: IQDSLSSTASALGK
Sample 5: ITFGGPSDSTGSNQNGER++
Sample 5: GQGVPINTNSSPDDQIGYYR
Sample 5: DGIIWVATEGALNTPK
Sample 5: NPANNAAIVLQLPQGTTLPK
Sample 5: GFYAEGSR
AQUA peptide
SARS-CoV-2 peptide
Fragment ions
Retention time (min)

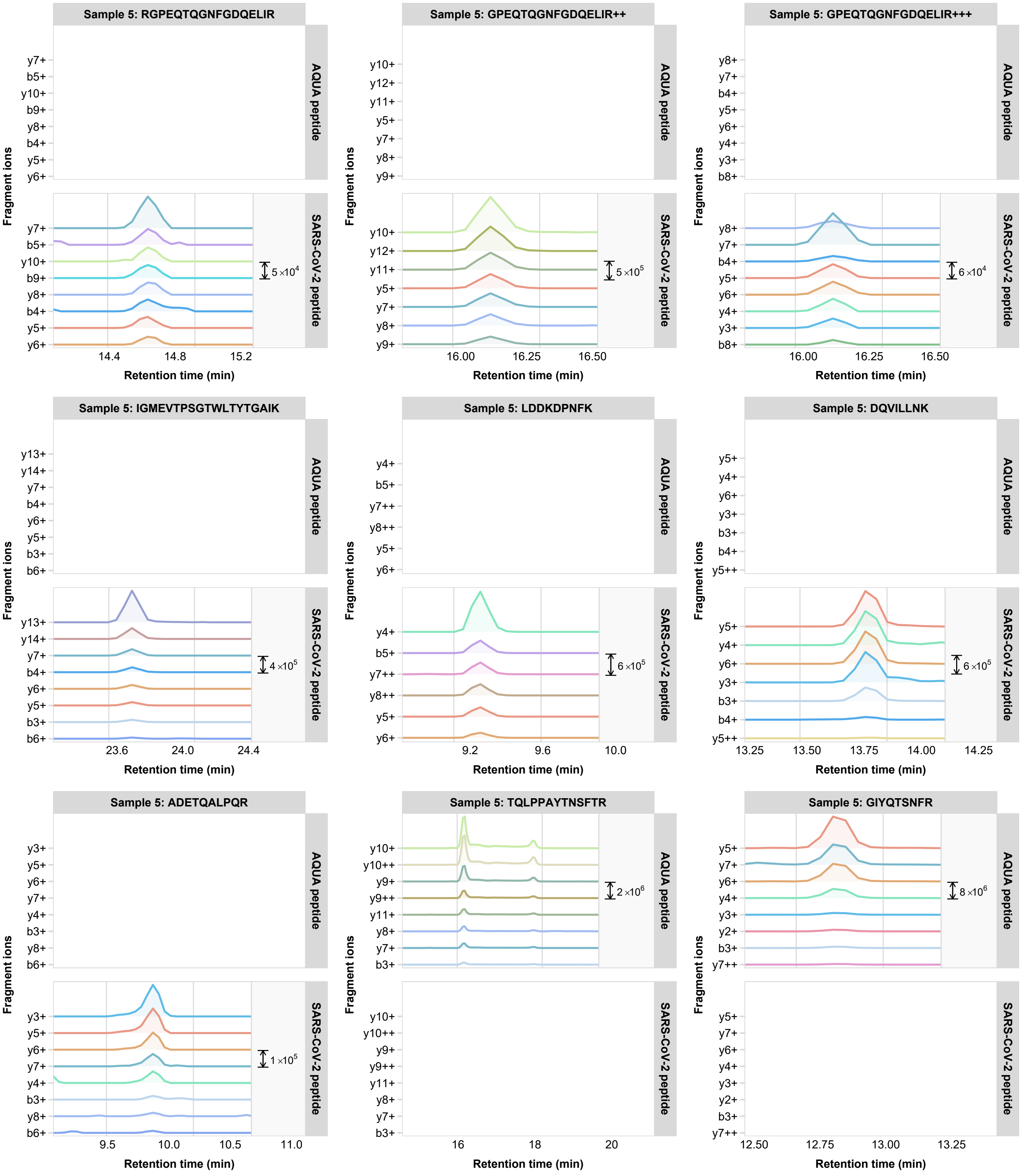

Sample 5: RGPEQTQGNFGDQELIR
Sample 5: GPEQTQGNFGDQELIR++
Sample 5: GPEQTQGNFGDQELIR+++
Sample 5: IGMEVTPSGTWLTYTGAIK
Sample 5: LDDKDPNFK
Sample 5: DQVILLNK
Sample 5: ADETQALPQR
Sample 5: TQLPPAYTNSFTR
Sample 5: GIYQTSNFR
AQUA peptide
SARS-CoV-2 peptide
Fragment ions
Retention time (min)
5×10^4
5×10^5
6×10^4
4×10^5
6×10^5
6×10^5
1×10^5
2×10^6
8×10^6

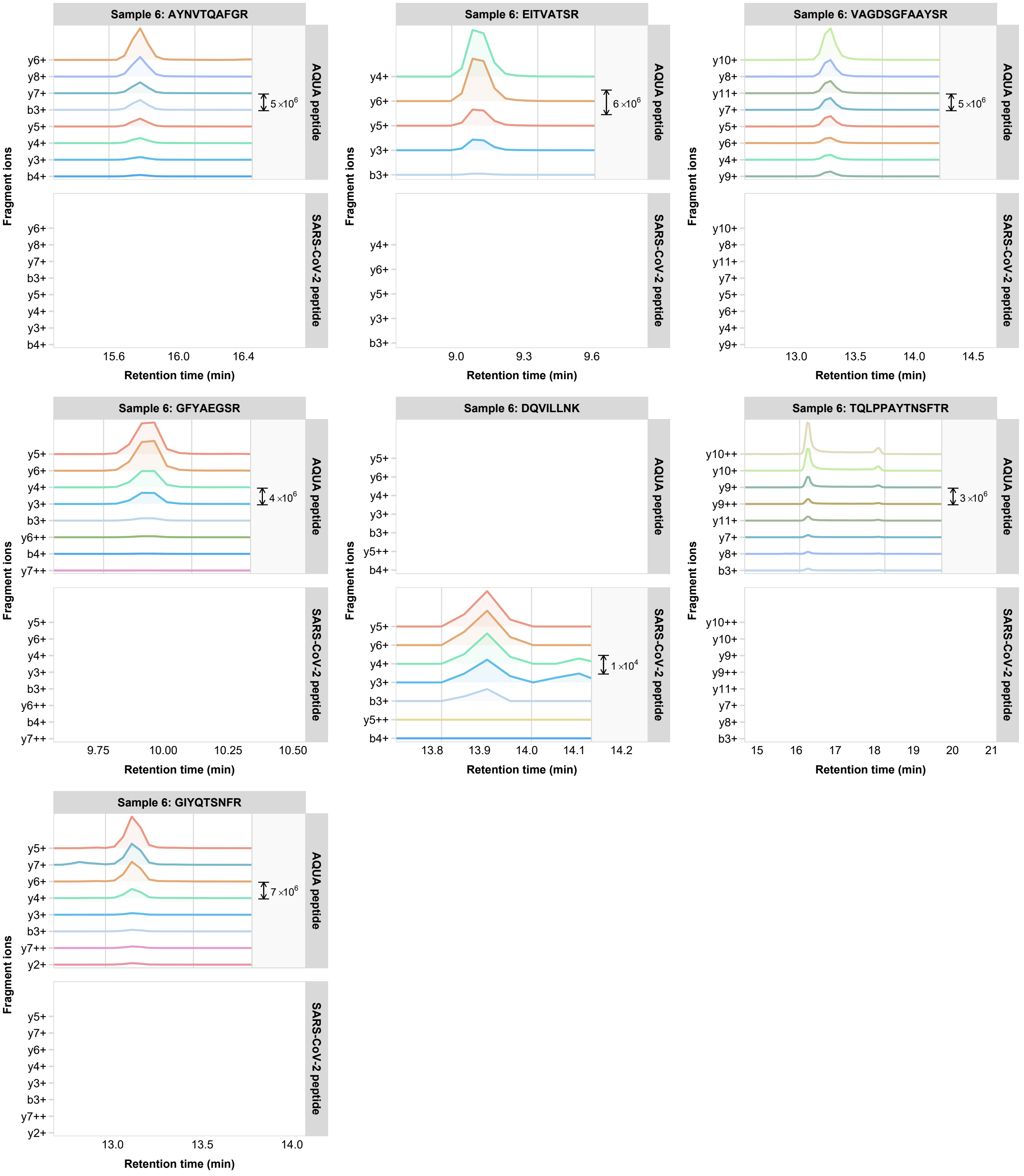

Sample 6: AYNVTQAFGR
Sample 6: EITVATSR
Sample 6: VAGDSGFAAYSR
Sample 6: GFYAEGSR
Sample 6: DQVILLNK
Sample 6: TQLPPAYTNSFTR
Sample 6: GIYQTSNFR
AQUA peptide
SARS-CoV-2 peptide
Fragment ions
Retention time (min)
5×10^6
6×10^6
4×10^6
1×10^4
3×10^6
7×10^6

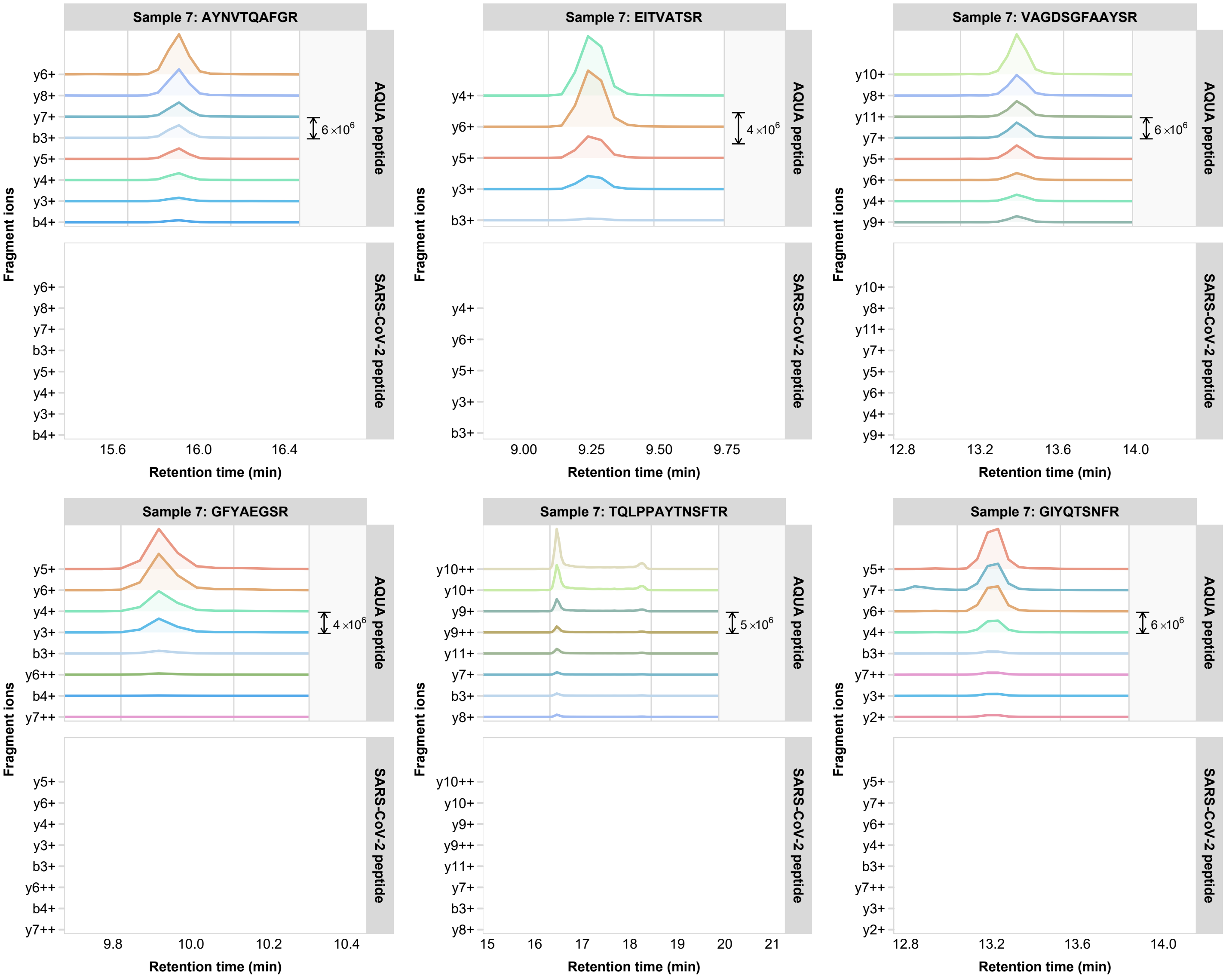
Sample 7: AYNVTQAFGR
Sample 7: EITVATSR
Sample 7: VAGDSGFAAYSR
Sample 7: GFYAEGSR
Sample 7: TQLPPAYTNSFTR
Sample 7: GIYQTSNFR
AQUA peptide
SARS-CoV-2 peptide
Fragment ions
Retention time (min)
6 ×10^6
4 ×10^6
5 ×10^6
y6+
y8+
y7+
b3+
y5+
y4+
y3+
b4+
y10+
y11+
y9+
y6++
y7++
y10++
y9++
y2+
y7++
15.6
16.0
16.4
9.00
9.25
9.50
9.75
12.8
13.2
13.6
14.0
9.8
10.0
10.2
10.4
15
16
17
18
19
20
21

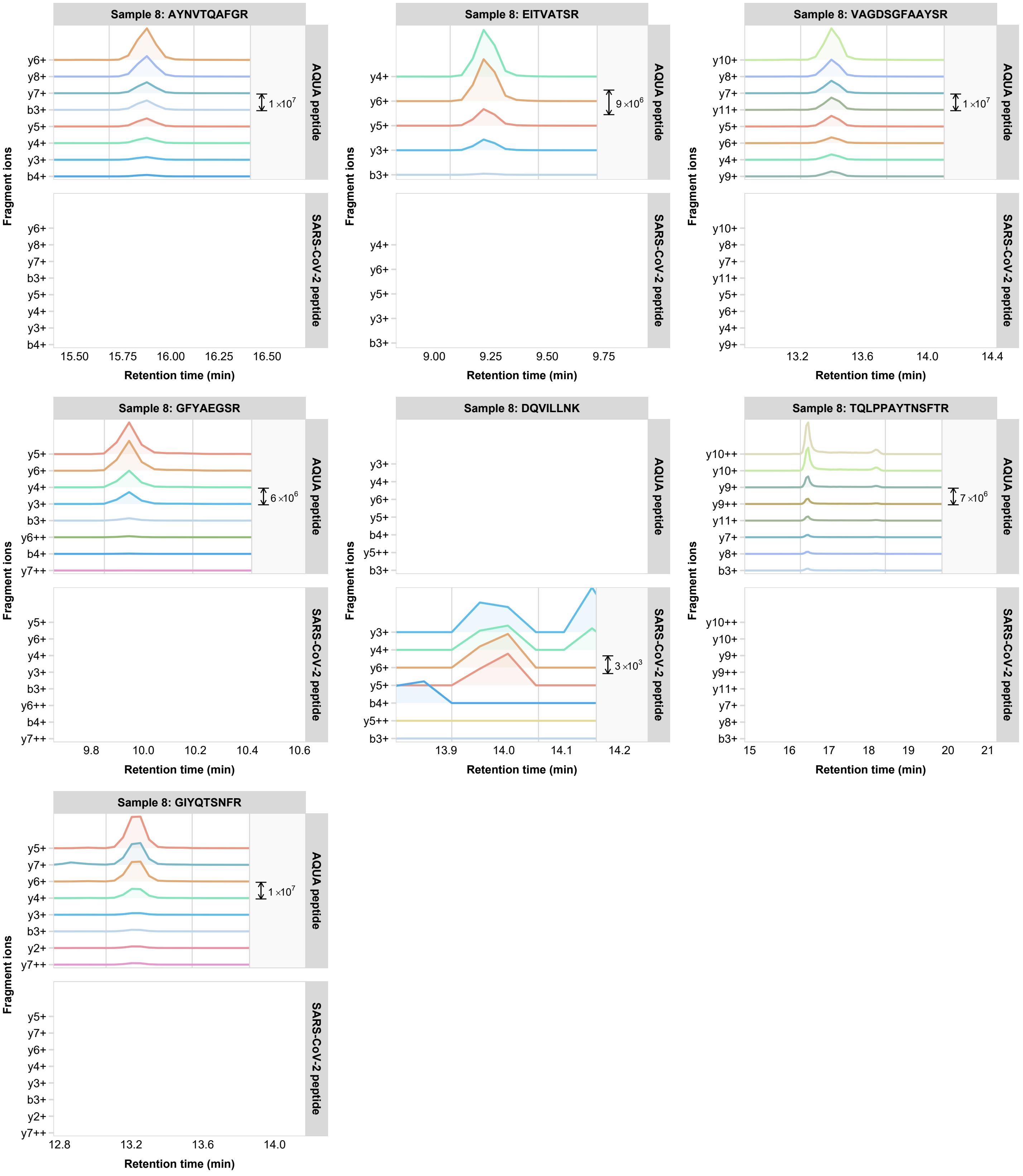

Sample 8: AYNVTQAFGR
AQUA peptide
SARS-CoV-2 peptide
y6+
y8+
y7+
b3+
y5+
y4+
y3+
b4+
1 ×10^7
15.50
15.75
16.00
16.25
16.50
Fragment ions
Retention time (min)
Sample 8: EITVATSR
y4+
y6+
y5+
y3+
b3+
9 ×10^6
9.00
9.25
9.50
9.75
Sample 8: VAGDSGFAAYSR
y10+
y8+
y7+
y11+
y5+
y6+
y4+
y9+
1 ×10^7
13.2
13.6
14.0
14.4
Sample 8: GFYAEGSR
y5+
y6+
y4+
y3+
b3+
y6++
b4+
y7++
6 ×10^6
9.8
10.0
10.2
10.4
10.6
Sample 8: DQVILLNK
y3+
y4+
y6+
y5+
b4+
y5++
b3+
3 ×10^3
13.9
14.0
14.1
14.2
Sample 8: TQLPPAYTNSFTR
y10++
y10+
y9+
y9++
y11+
y7+
y8+
b3+
7 ×10^6
15
16
17
18
19
20
21
Sample 8: GIYQTSNFR
y5+
y7+
y6+
y4+
y3+
b3+
y2+
y7++
1 ×10^7
12.8
13.2
13.6
14.0

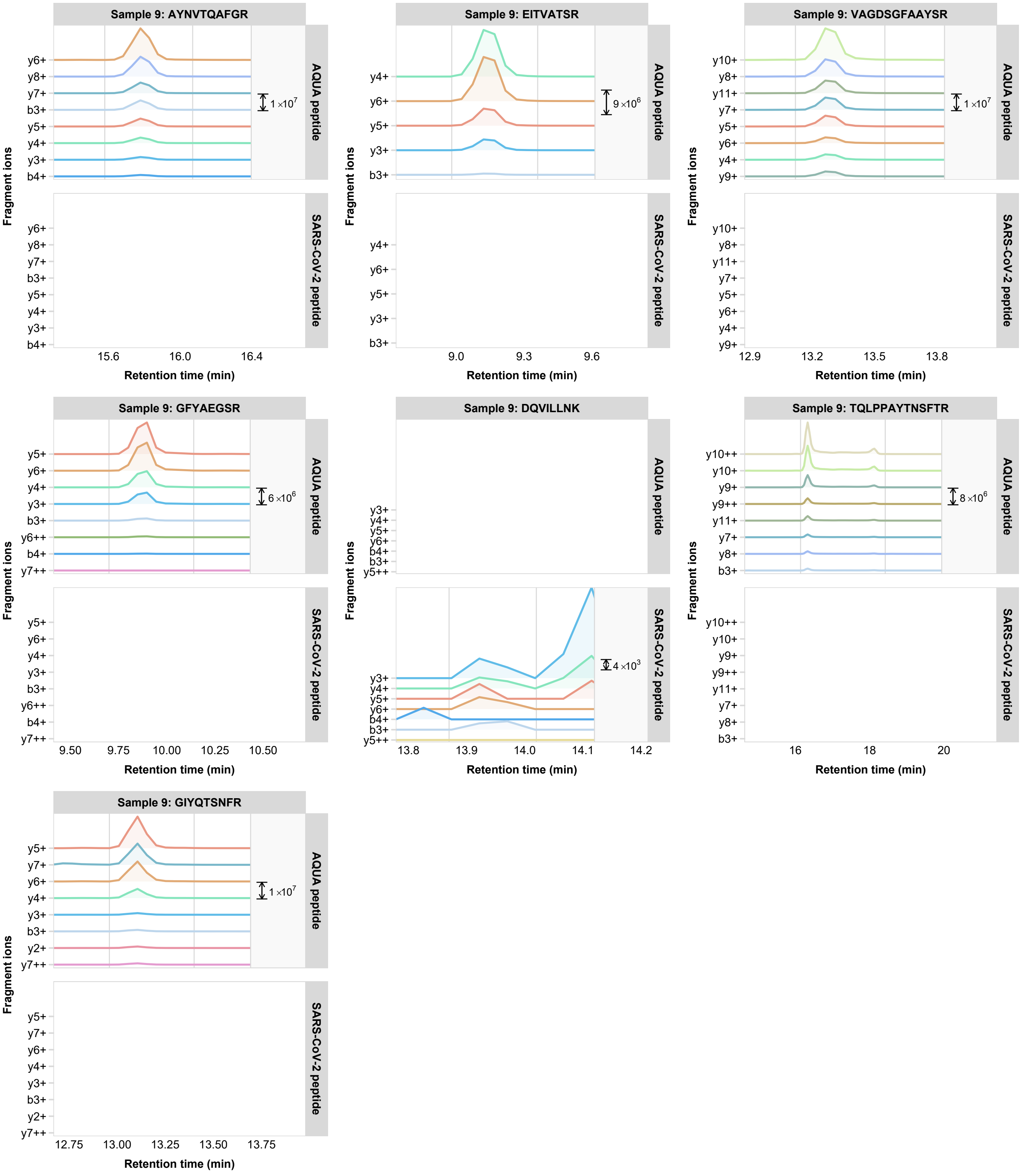
Sample 9: AYNVTQAFGR
AQUA peptide
SARS-CoV-2 peptide
1×10^7
y6+ y8+ y7+ b3+ y5+ y4+ y3+ b4+
15.6 16.0 16.4
Sample 9: EITVATSR
9×10^6
y4+ y6+ y5+ y3+ b3+
9.0 9.3 9.6
Sample 9: VAGDSGFAAYSR
1×10^7
y10+ y8+ y11+ y7+ y5+ y6+ y4+ y9+
12.9 13.2 13.5 13.8
Sample 9: GFYAEGSR
6×10^6
y5+ y6+ y4+ y3+ b3+ y6++ b4+ y7++
9.50 9.75 10.00 10.25 10.50
Sample 9: DQVILLNK
4×10^3
y3+ y4+ y5+ y6+ b4+ b3+ y5++
13.8 13.9 14.0 14.1 14.2
Sample 9: TQLPPAYTNSFTR
8×10^6
y10++ y10+ y9+ y9++ y11+ y7+ y8+ b3+
16 18 20
Sample 9: GIYQTSNFR
1×10^7
y5+ y7+ y6+ y4+ y3+ b3+ y2+ y7++
12.75 13.00 13.25 13.50 13.75
Fragment ions
Retention time (min)

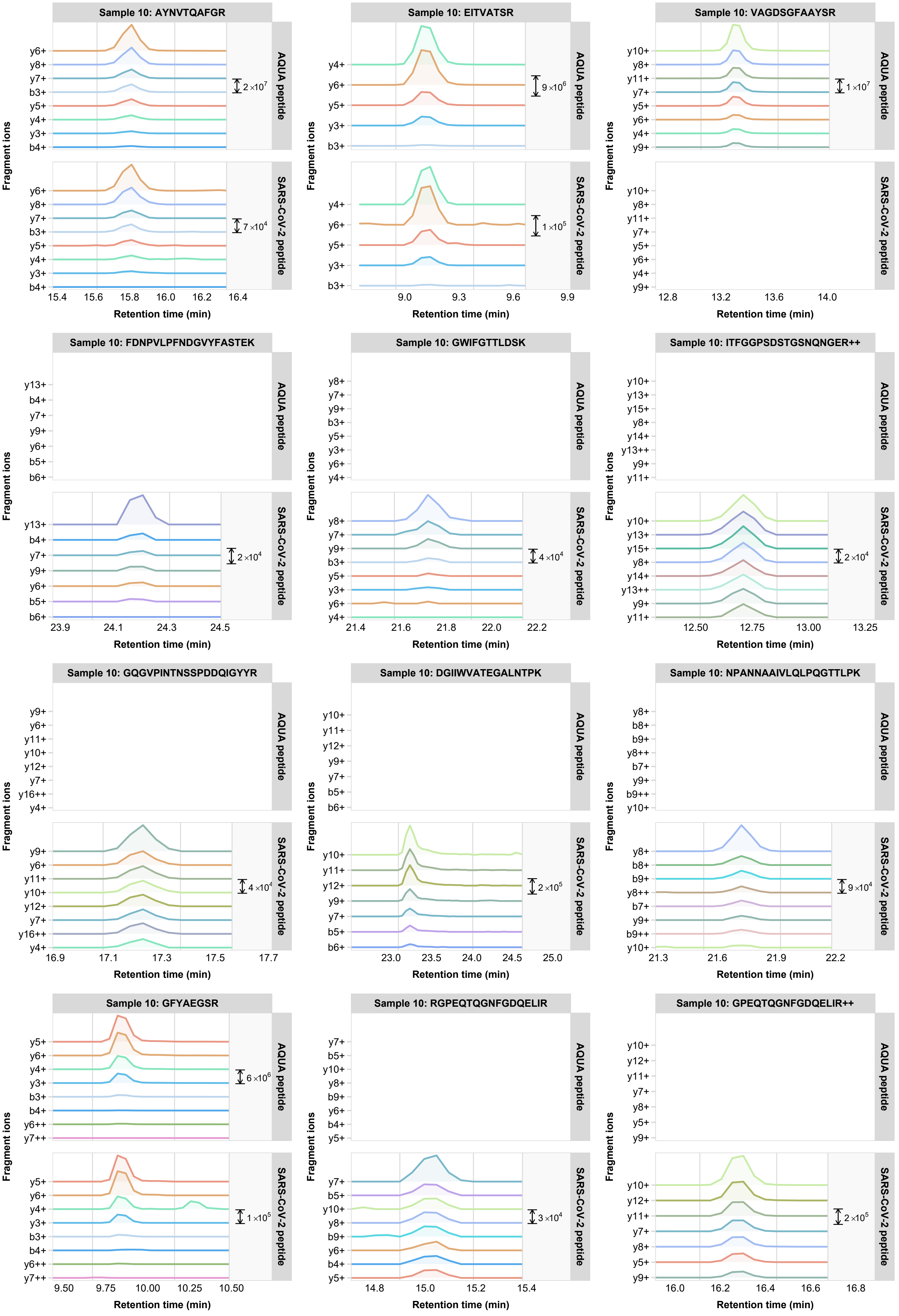
Sample 10: AYNVTQAFGR
Sample 10: EITVATSR
Sample 10: VAGDSGFAAYSR
Sample 10: FDNPVLPFNDGVYFASTEK
Sample 10: GWIFGTTLDSK
Sample 10: ITFGGPSDSTGSNQNGER++
Sample 10: GQGVPINTNSSPDDQIGYYR
Sample 10: DGIIWVATEGALNTPK
Sample 10: NPANNAAIVLQLPQGTTLPK
Sample 10: GFYAEGSR
Sample 10: RGPEQTQGNFGDQELIR
Sample 10: GPEQTQGNFGDQELIR++
AQUA peptide
SARS-CoV-2 peptide
Fragment ions
Retention time (min)
2×10^7
7×10^4
9×10^6
1×10^5
1×10^7
2×10^4
4×10^4
2×10^4
4×10^4
2×10^5
9×10^4
6×10^6
1×10^5
3×10^4
2×10^5
y6+ y8+ y7+ b3+ y5+ y4+ y3+ b4+
y4+ y6+ y5+ y3+ b3+
y10+ y8+ y11+ y7+ y5+ y6+ y4+ y9+
y13+ b4+ y7+ y9+ y6+ b5+ b6+
y8+ y7+ y9+ b3+ y5+ y3+ y6+ y4+
y10+ y13+ y15+ y8+ y14+ y13++ y9+ y11+
y9+ y6+ y11+ y10+ y12+ y7+ y16++ y4+
y10+ y11+ y12+ y9+ y7+ b5+ b6+
y8+ b8+ b9+ y8++ b7+ y9+ b9++ y10+
y5+ y6+ y4+ y3+ b3+ b4+ y6++ y7++
y7+ b5+ y10+ y8+ b9+ y6+ b4+ y5+
y10+ y12+ y11+ y7+ y8+ y5+ y9+
15.4 15.6 15.8 16.0 16.2 16.4
9.0 9.3 9.6 9.9
12.8 13.2 13.6 14.0
23.9 24.1 24.3 24.5
21.4 21.6 21.8 22.0 22.2
12.50 12.75 13.00 13.25
16.9 17.1 17.3 17.5 17.7
23.0 23.5 24.0 24.5 25.0
21.3 21.6 21.9 22.2
9.50 9.75 10.00 10.25 10.50
14.8 15.0 15.2 15.4
16.0 16.2 16.4 16.6 16.8

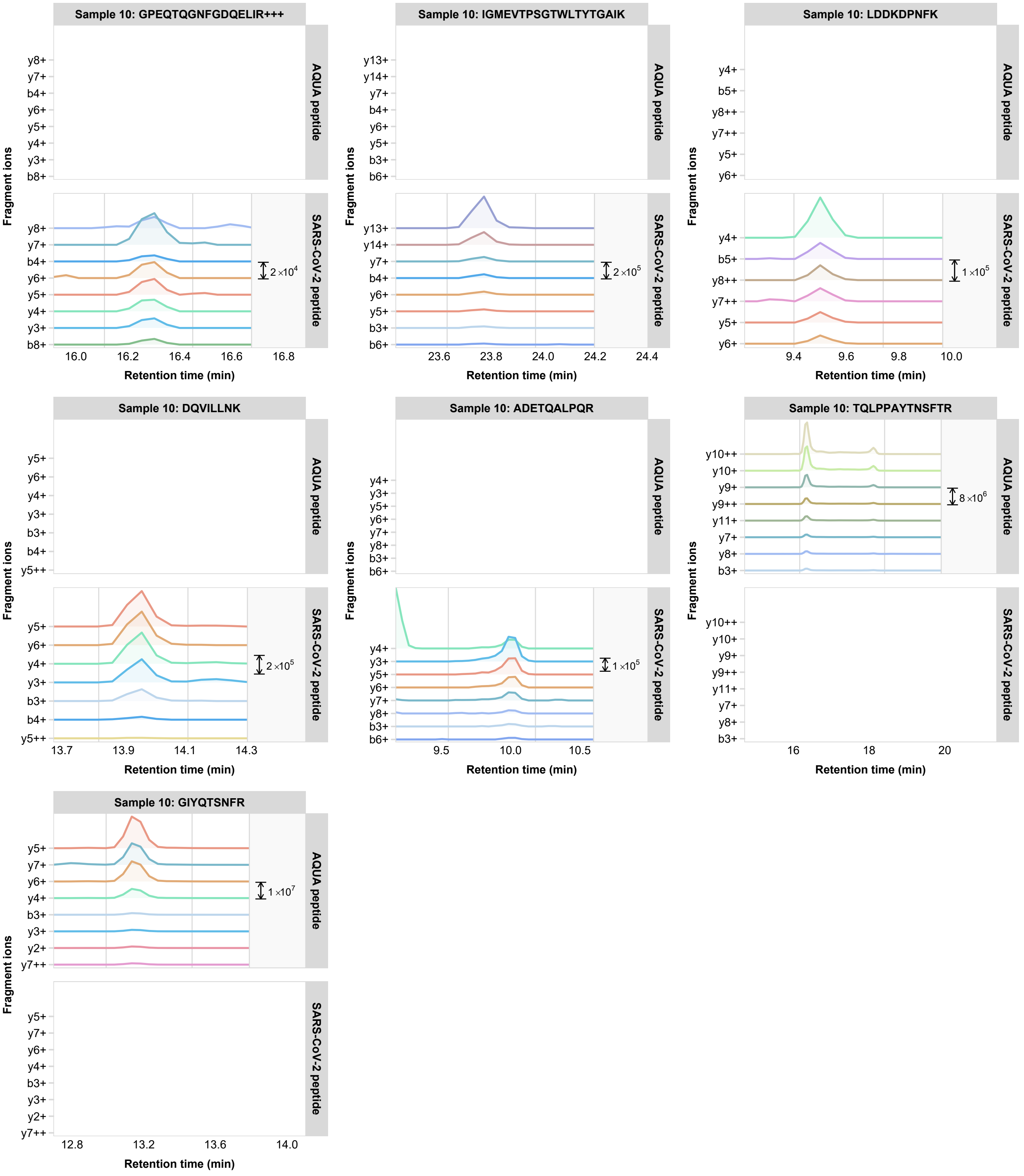

Sample 10: GPEQTQGNFGDQELIR+++
AQUA peptide
SARS-CoV-2 peptide
y8+
y7+
b4+
y6+
y5+
y4+
y3+
b8+
2×10^4
16.0
16.2
16.4
16.6
16.8
Fragment ions
Retention time (min)
Sample 10: IGMEVTPSGTWLTYTGAIK
AQUA peptide
SARS-CoV-2 peptide
y13+
y14+
y7+
b4+
y6+
y5+
b3+
b6+
2×10^5
23.6
23.8
24.0
24.2
24.4
Fragment ions
Retention time (min)
Sample 10: LDDKDPNFK
AQUA peptide
SARS-CoV-2 peptide
y4+
b5+
y8++
y7++
y5+
y6+
1×10^5
9.4
9.6
9.8
10.0
Fragment ions
Retention time (min)
Sample 10: DQVILLNK
AQUA peptide
SARS-CoV-2 peptide
y5+
y6+
y4+
y3+
b3+
b4+
y5++
2×10^5
13.7
13.9
14.1
14.3
Fragment ions
Retention time (min)
Sample 10: ADETQALPQR
AQUA peptide
SARS-CoV-2 peptide
y4+
y3+
y5+
y6+
y7+
y8+
b3+
b6+
1×10^5
9.5
10.0
10.5
Fragment ions
Retention time (min)
Sample 10: TQLPPAYTNSFTR
AQUA peptide
SARS-CoV-2 peptide
y10++
y10+
y9+
y9++
y11+
y7+
y8+
b3+
8×10^6
16
18
20
Fragment ions
Retention time (min)
Sample 10: GIYQTSNFR
AQUA peptide
SARS-CoV-2 peptide
y5+
y7+
y6+
y4+
b3+
y3+
y2+
y7++
1×10^7
12.8
13.2
13.6
14.0
Fragment ions
Retention time (min)

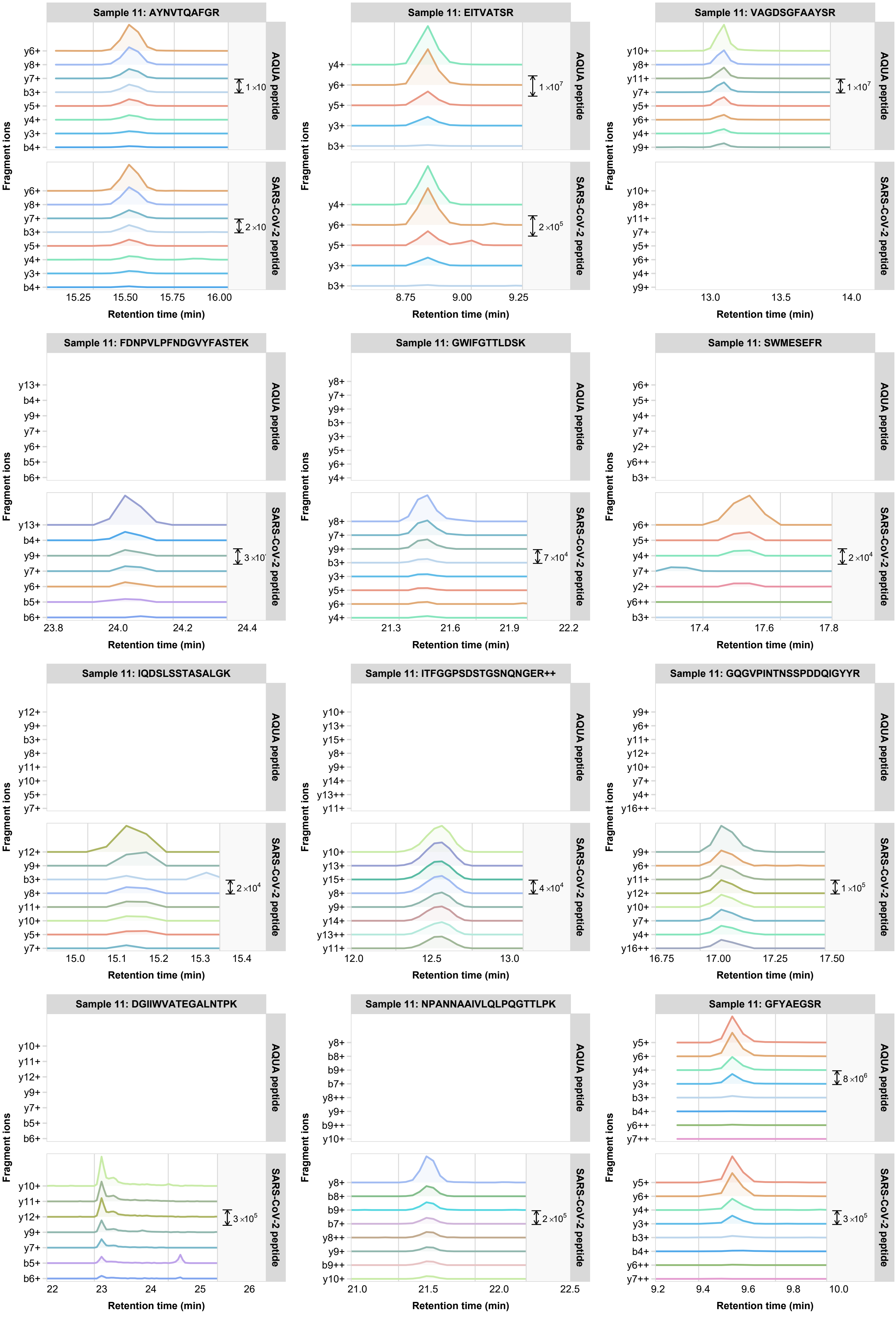

Sample 11: AYNVTQAFGR
AQUA peptide
SARS-CoV-2 peptide
y6+ y8+ y7+ b3+ y5+ y4+ y3+ b4+
1×10
2×10
15.25 15.50 15.75 16.00
Sample 11: EITVATSR
y4+ y6+ y5+ y3+ b3+
1×10^7
2×10^5
8.75 9.00 9.25
Sample 11: VAGDSGFAAYSR
y10+ y8+ y11+ y7+ y5+ y6+ y4+ y9+
1×10^7
13.0 13.5 14.0
Sample 11: FDNPVLPFNDGVYFASTEK
y13+ b4+ y9+ y7+ y6+ b5+ b6+
3×10
23.8 24.0 24.2 24.4
Sample 11: GWIFGTTLDSK
y8+ y7+ y9+ b3+ y3+ y5+ y6+ y4+
7×10^4
21.3 21.6 21.9 22.2
Sample 11: SWMESEFR
y6+ y5+ y4+ y7+ y2+ y6++ b3+
2×10^4
17.4 17.6 17.8
Sample 11: IQDSLSSTASALGK
y12+ y9+ b3+ y8+ y11+ y10+ y5+ y7+
2×10^4
15.0 15.1 15.2 15.3 15.4
Sample 11: ITFGGPSDSTGSNQNGER++
y10+ y13+ y15+ y8+ y9+ y14+ y13++ y11+
4×10^4
12.0 12.5 13.0
Sample 11: GQGVPINTNSSPDDQIGYYR
y9+ y6+ y11+ y12+ y10+ y7+ y4+ y16++
1×10^5
16.75 17.00 17.25 17.50
Sample 11: DGIIWVATEGALNTPK
y10+ y11+ y12+ y9+ y7+ b5+ b6+
3×10^5
22 23 24 25 26
Sample 11: NPANNAAIVLQLPQGTTLPK
y8+ b8+ b9+ b7+ y8++ y9+ b9++ y10+
2×10^5
21.0 21.5 22.0 22.5
Sample 11: GFYAEGSR
y5+ y6+ y4+ y3+ b3+ b4+ y6++ y7++
8×10^6
3×10^5
9.2 9.4 9.6 9.8 10.0
Fragment ions
Retention time (min)

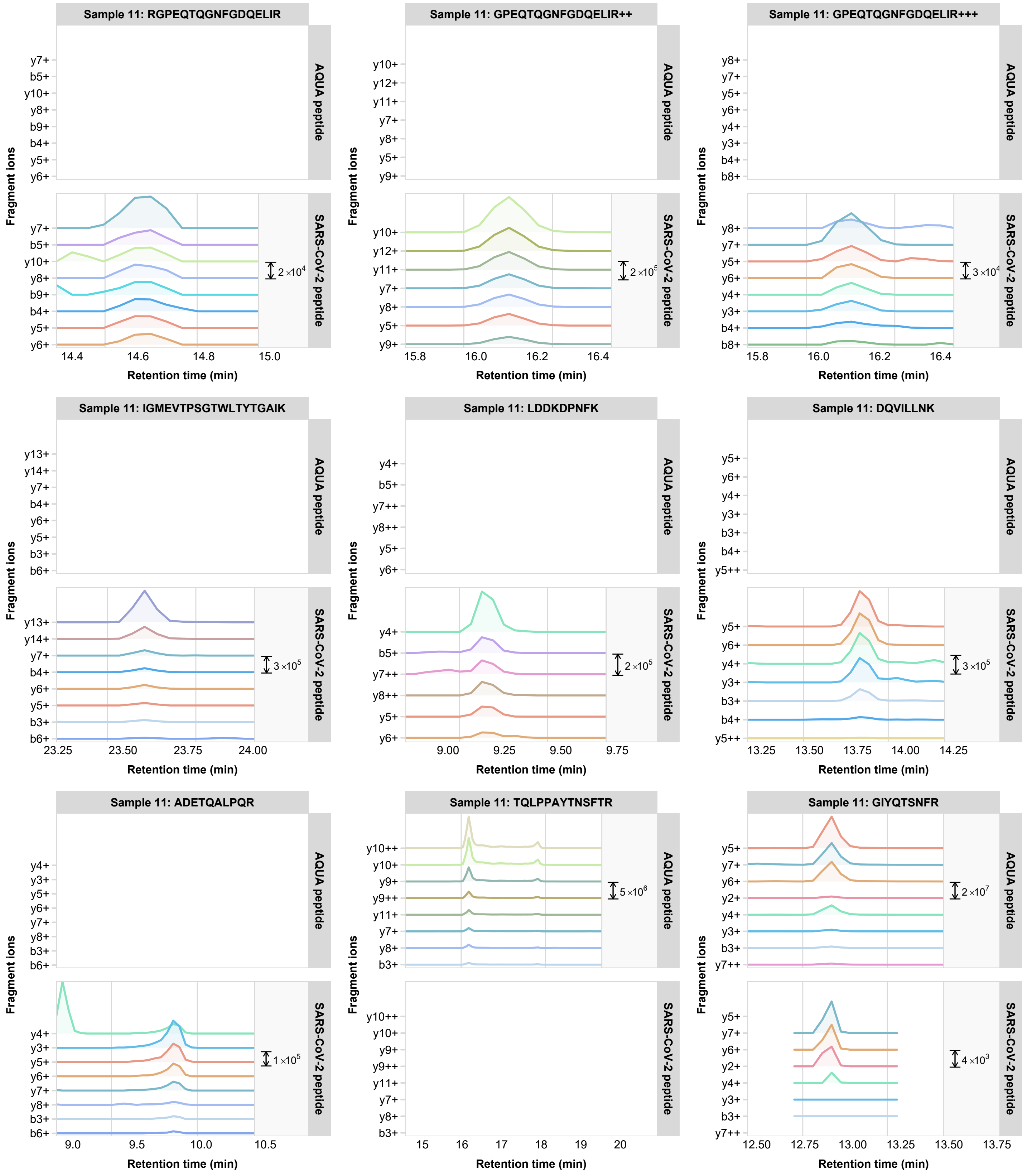

Sample 11: RGPEQTQGNFGDQELIR
Sample 11: GPEQTQGNFGDQELIR++
Sample 11: GPEQTQGNFGDQELIR+++
Sample 11: IGMEVTPSGTWLTYTGAIK
Sample 11: LDDKDPNFK
Sample 11: DQVILLNK
Sample 11: ADETQALPQR
Sample 11: TQLPPAYTNSFTR
Sample 11: GIYQTSNFR
AQUA peptide
SARS-CoV-2 peptide
Fragment ions
Retention time (min)
y7+ b5+ y10+ y8+ b9+ b4+ y5+ y6+
y10+ y12+ y11+ y7+ y8+ y5+ y9+
y8+ y7+ y5+ y6+ y4+ y3+ b4+ b8+
y13+ y14+ y7+ b4+ y6+ y5+ b3+ b6+
y4+ b5+ y7++ y8++ y5+ y6+
y5+ y6+ y4+ y3+ b3+ b4+ y5++
y4+ y3+ y5+ y6+ y7+ y8+ b3+ b6+
y10++ y10+ y9+ y9++ y11+ y7+ y8+ b3+
y5+ y7+ y6+ y2+ y4+ y3+ b3+ y7++
2×10^4
2×10^5
3×10^4
3×10^5
2×10^5
3×10^5
1×10^5
5×10^6
2×10^7
4×10^3
14.4 14.6 14.8 15.0
15.8 16.0 16.2 16.4
23.25 23.50 23.75 24.00
9.00 9.25 9.50 9.75
13.25 13.50 13.75 14.00 14.25
9.0 9.5 10.0 10.5
15 16 17 18 19 20
12.50 12.75 13.00 13.25 13.50 13.75
