## Supplemental Information 2 for "Targeted proteomics as a tool to detect SARS-CoV-2 proteins in clinical specimens"

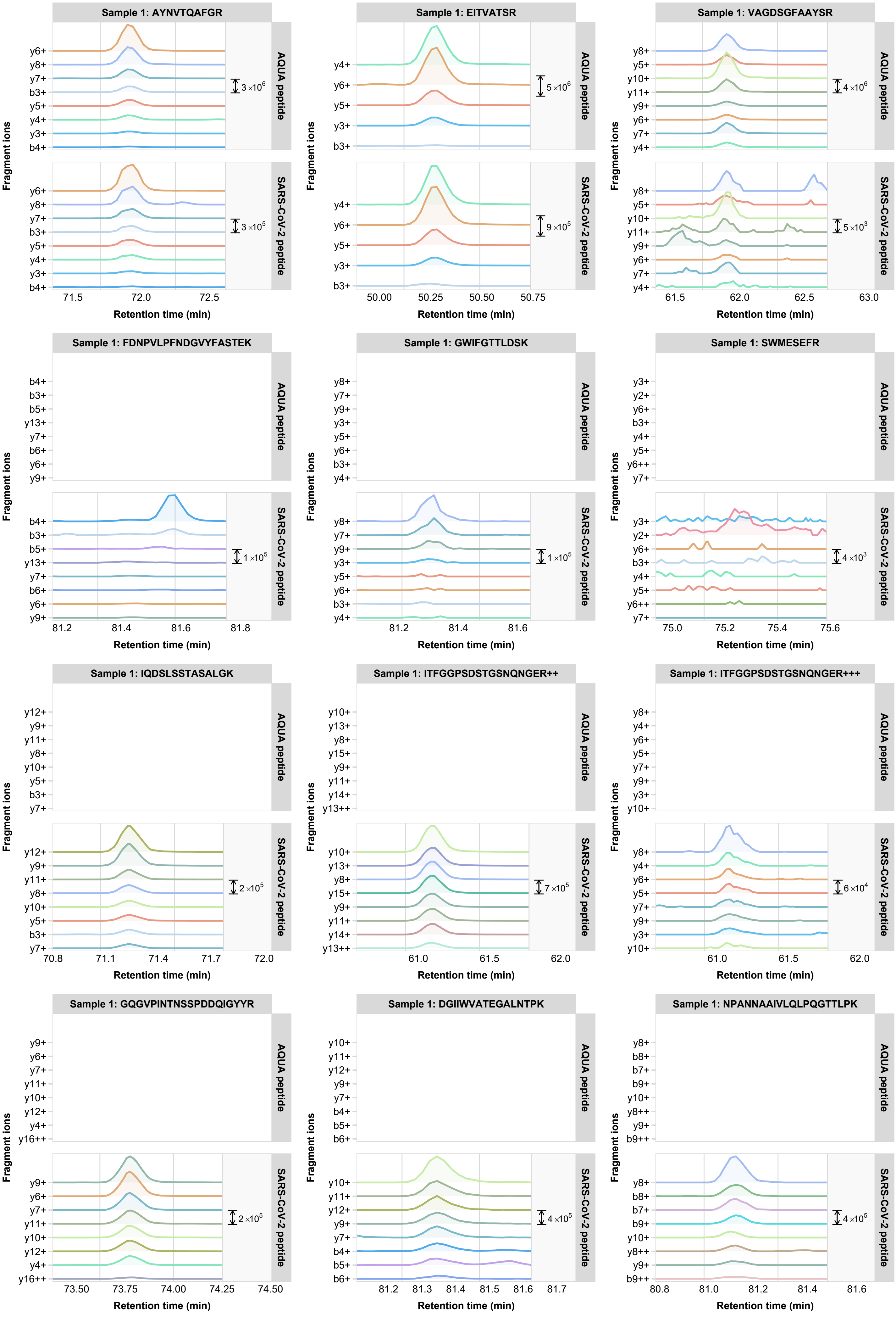
Sample 1: AYNVTQAFGR
AQUA peptide
SARS-CoV-2 peptide
y6+ y8+ y7+ b3+ y5+ y4+ y3+ b4+
$3 \times 10^6$
$3 \times 10^5$
71.5 72.0 72.5
Sample 1: EITVATSR
y4+ y6+ y5+ y3+ b3+
$5 \times 10^6$
$9 \times 10^5$
50.00 50.25 50.50 50.75
Sample 1: VAGDSGFAAYSR
y8+ y5+ y10+ y11+ y9+ y6+ y7+ y4+
$4 \times 10^6$
$5 \times 10^3$
61.5 62.0 62.5 63.0
Sample 1: FDNPVLPFNDGVYFASTEK
b4+ b3+ b5+ y13+ y7+ b6+ y6+ y9+
$1 \times 10^5$
81.2 81.4 81.6 81.8
Sample 1: GWIFGTTLDSK
y8+ y7+ y9+ y3+ y5+ y6+ b3+ y4+
$1 \times 10^5$
81.2 81.4 81.6
Sample 1: SWMESEFR
y3+ y2+ y6+ b3+ y4+ y5+ y6++ y7+
$4 \times 10^3$
75.0 75.2 75.4 75.6
Sample 1: IQDSLSSTASALGK
y12+ y9+ y11+ y8+ y10+ y5+ b3+ y7+
$2 \times 10^5$
70.8 71.1 71.4 71.7 72.0
Sample 1: ITFGGPSDSTGSNQNGER++
y10+ y13+ y8+ y15+ y9+ y11+ y14+ y13++
$7 \times 10^5$
61.0 61.5 62.0
Sample 1: ITFGGPSDSTGSNQNGER+++
y8+ y4+ y6+ y5+ y7+ y9+ y3+ y10+
$6 \times 10^4$
61.0 61.5 62.0
Sample 1: GQGVPINTNSSPDDQIGYYR
y9+ y6+ y7+ y11+ y10+ y12+ y4+ y16++
$2 \times 10^5$
73.50 73.75 74.00 74.25 74.50
Sample 1: DGIIWVATEGALNTPK
y10+ y11+ y12+ y9+ y7+ b4+ b5+ b6+
$4 \times 10^5$
81.2 81.3 81.4 81.5 81.6 81.7
Sample 1: NPANNAAIVLQLPQGTTLPK
y8+ b8+ b7+ b9+ y10+ y8++ y9+ b9++
$4 \times 10^5$
80.8 81.0 81.2 81.4 81.6
Fragment ions
Retention time (min)

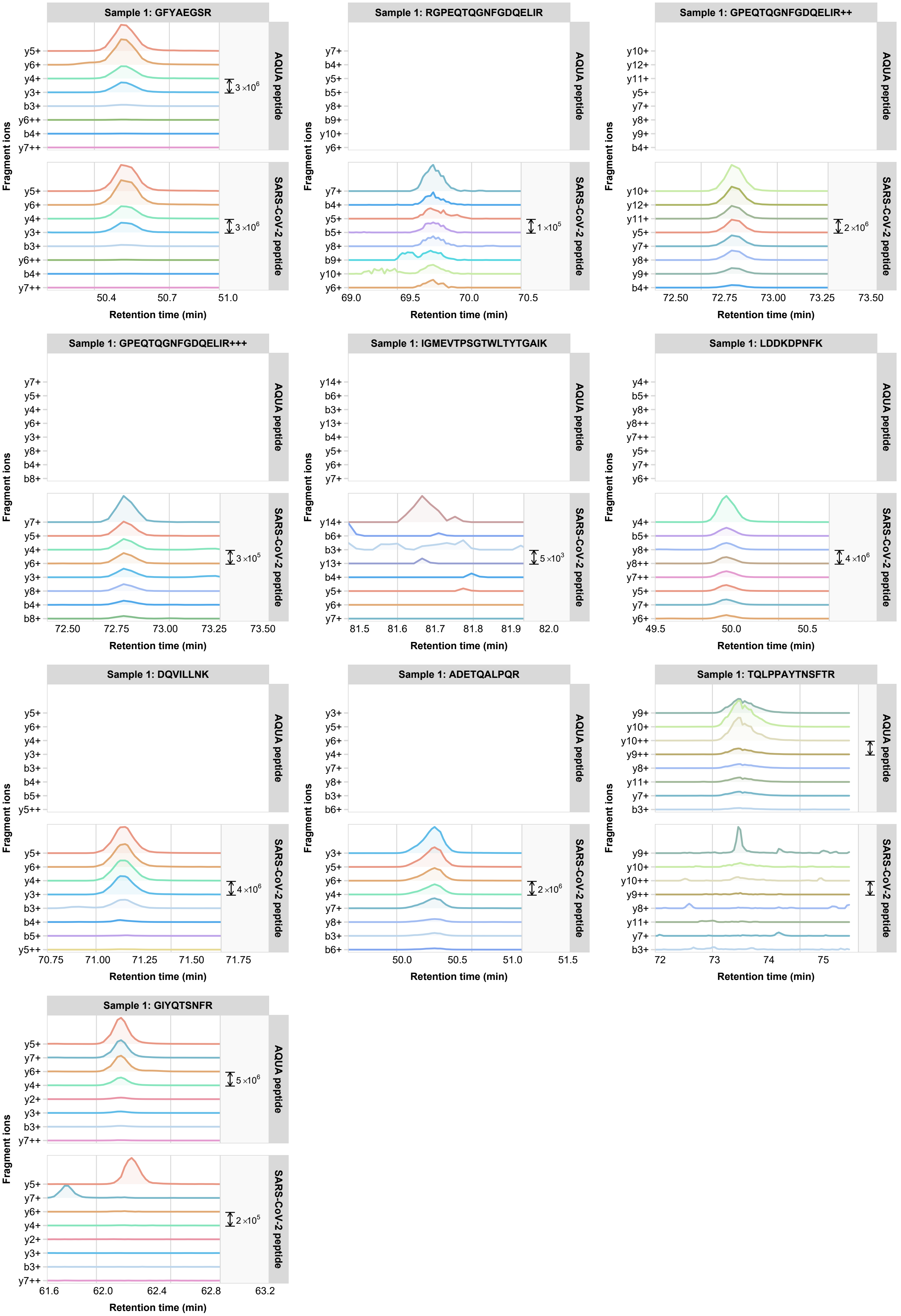
Sample 1: GFYAEGSR
AQUA peptide
SARS-CoV-2 peptide
y5+ y6+ y4+ y3+ b3+ y6++ b4+ y7++
3 ×10^6
50.4 50.7 51.0
Sample 1: RGPEQTQGNFGDQELIR
y7+ b4+ y5+ b5+ y8+ b9+ y10+ y6+
1 ×10^5
69.0 69.5 70.0 70.5
Sample 1: GPEQTQGNFGDQELIR++
y10+ y12+ y11+ y5+ y7+ y8+ y9+ b4+
2 ×10^6
72.50 72.75 73.00 73.25 73.50
Sample 1: GPEQTQGNFGDQELIR+++
y7+ y5+ y4+ y6+ y3+ y8+ b4+ b8+
3 ×10^5
72.50 72.75 73.00 73.25 73.50
Sample 1: IGMEVTPSGTWLTYTGAIK
y14+ b6+ b3+ y13+ b4+ y5+ y6+ y7+
5 ×10^3
81.5 81.6 81.7 81.8 81.9 82.0
Sample 1: LDDKDPNFK
y4+ b5+ y8+ y8++ y7++ y5+ y7+ y6+
4 ×10^6
49.5 50.0 50.5
Sample 1: DQVILLNK
y5+ y6+ y4+ y3+ b3+ b4+ b5+ y5++
4 ×10^6
70.75 71.00 71.25 71.50 71.75
Sample 1: ADETQALPQR
y3+ y5+ y6+ y4+ y7+ y8+ b3+ b6+
2 ×10^6
50.0 50.5 51.0 51.5
Sample 1: TQLPPAYTNSFTR
y9+ y10+ y10++ y9++ y8+ y11+ y7+ b3+
72 73 74 75
Sample 1: GIYQTSNFR
y5+ y7+ y6+ y4+ y2+ y3+ b3+ y7++
5 ×10^6
2 ×10^5
61.6 62.0 62.4 62.8 63.2
Fragment ions
Retention time (min)

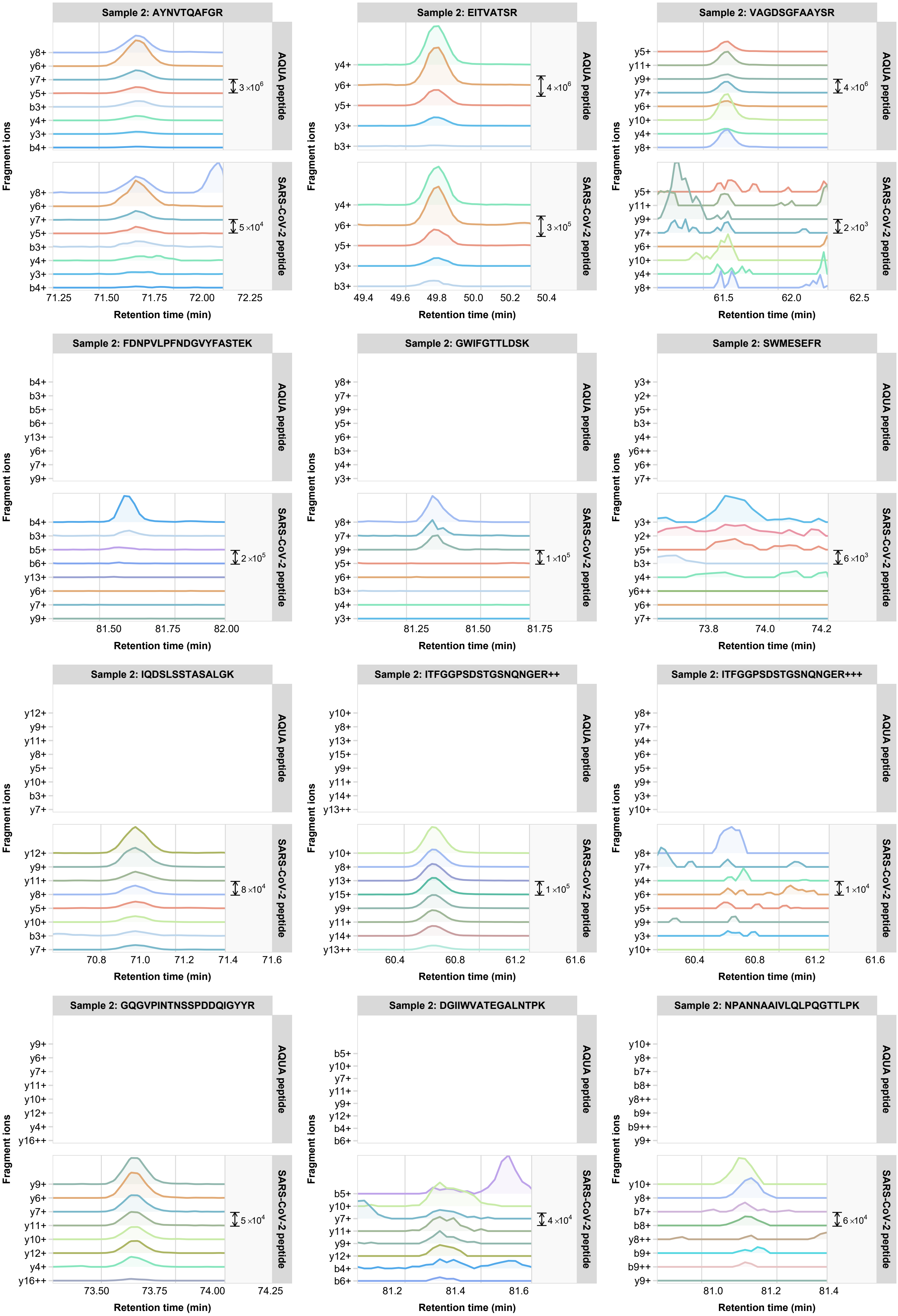

Sample 2: AYNVTQAFGR
AQUA peptide
SARS-CoV-2 peptide
y8+ y6+ y7+ y5+ b3+ y4+ y3+ b4+
3×10^6
5×10^4
71.25 71.50 71.75 72.00 72.25
Sample 2: EITVATSR
y4+ y6+ y5+ y3+ b3+
4×10^6
3×10^5
49.4 49.6 49.8 50.0 50.2 50.4
Sample 2: VAGDSGFAAYSR
y5+ y11+ y9+ y7+ y6+ y10+ y4+ y8+
4×10^6
2×10^3
61.5 62.0 62.5
Sample 2: FDNPVLPFNDGVYFASTEK
b4+ b3+ b5+ b6+ y13+ y6+ y7+ y9+
2×10^5
81.50 81.75 82.00
Sample 2: GWIFGTTLDSK
y8+ y7+ y9+ y5+ y6+ b3+ y4+ y3+
1×10^5
81.25 81.50 81.75
Sample 2: SWMESEFR
y3+ y2+ y5+ b3+ y4+ y6++ y6+ y7+
6×10^3
73.8 74.0 74.2
Sample 2: IQDSLSSTASALGK
y12+ y9+ y11+ y8+ y5+ y10+ b3+ y7+
8×10^4
70.8 71.0 71.2 71.4 71.6
Sample 2: ITFGGPSDSTGSNQNGER++
y10+ y8+ y13+ y15+ y9+ y11+ y14+ y13++
1×10^5
60.4 60.8 61.2 61.6
Sample 2: ITFGGPSDSTGSNQNGER+++
y8+ y7+ y4+ y6+ y5+ y9+ y3+ y10+
1×10^4
60.4 60.8 61.2 61.6
Sample 2: GQGVPINTNSSPDDQIGYYR
y9+ y6+ y7+ y11+ y10+ y12+ y4+ y16++
5×10^4
73.50 73.75 74.00 74.25
Sample 2: DGIIWVATEGALNTPK
b5+ y10+ y7+ y11+ y9+ y12+ b4+ b6+
4×10^4
81.2 81.4 81.6
Sample 2: NPANNAAIVLQLPQGTTLPK
y10+ y8+ b7+ b8+ y8++ b9+ b9++ y9+
6×10^4
81.0 81.2 81.4
Fragment ions
Retention time (min)

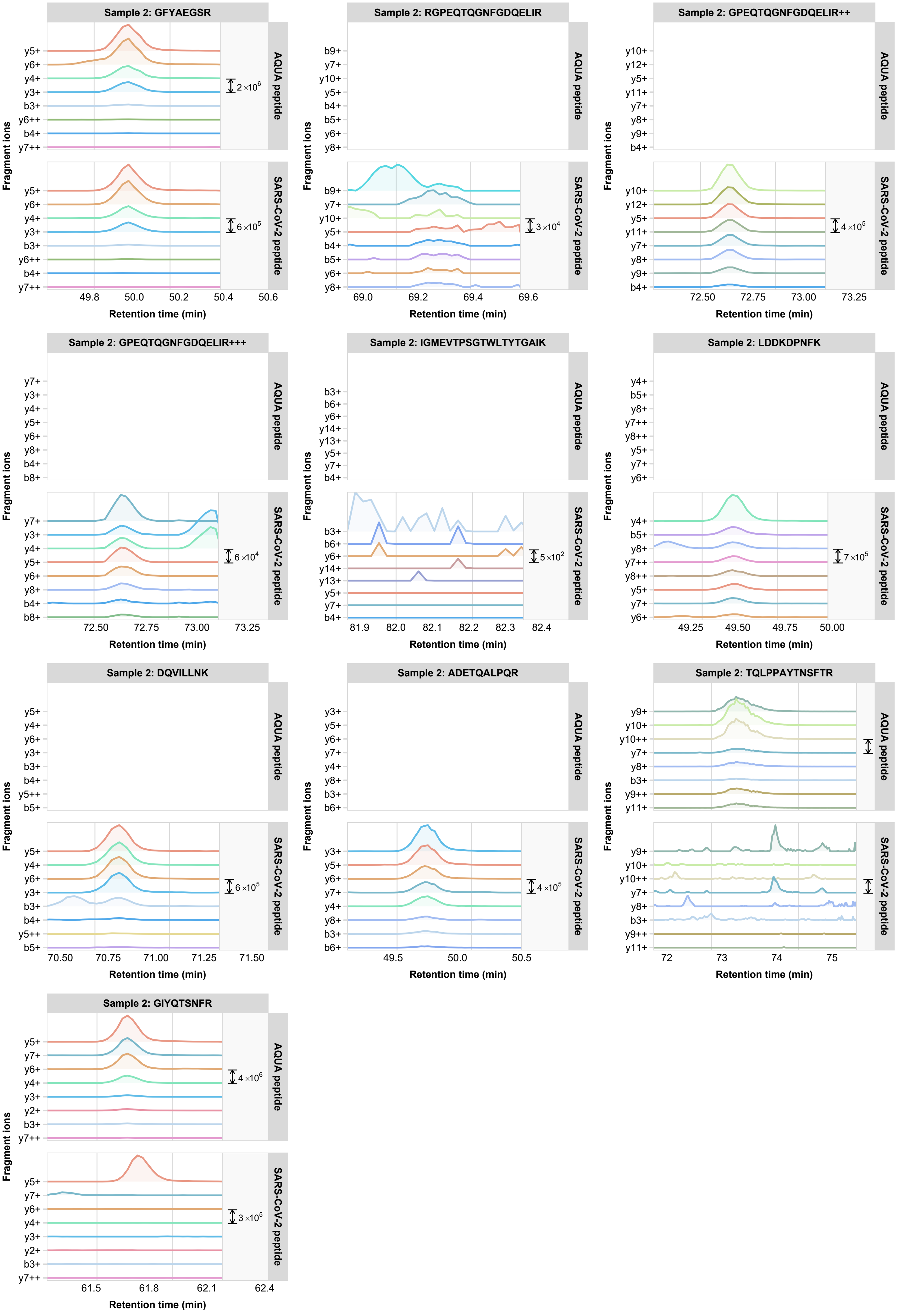
Sample 2: GFYAEGSR
y5+ y6+ y4+ y3+ b3+ y6++ b4+ y7++
2×10^6
6×10^5
49.8 50.0 50.2 50.4 50.6
Sample 2: RGPEQTQGNFGDQELIR
b9+ y7+ y10+ y5+ b4+ b5+ y6+ y8+
3×10^4
69.0 69.2 69.4 69.6
Sample 2: GPEQTQGNFGDQELIR++
y10+ y12+ y5+ y11+ y7+ y8+ y9+ b4+
4×10^5
72.50 72.75 73.00 73.25
Sample 2: GPEQTQGNFGDQELIR+++
y7+ y3+ y4+ y5+ y6+ y8+ b4+ b8+
6×10^4
72.50 72.75 73.00 73.25
Sample 2: IGMEVTPSGTWLTYTGAIK
b3+ b6+ y6+ y14+ y13+ y5+ y7+ b4+
5×10^2
81.9 82.0 82.1 82.2 82.3 82.4
Sample 2: LDDKDPNFK
y4+ b5+ y8+ y7++ y8++ y5+ y7+ y6+
7×10^5
49.25 49.50 49.75 50.00
Sample 2: DQVILLNK
y5+ y4+ y6+ y3+ b3+ b4+ y5++ b5+
6×10^5
70.50 70.75 71.00 71.25 71.50
Sample 2: ADETQALPQR
y3+ y5+ y6+ y7+ y4+ y8+ b3+ b6+
4×10^5
49.5 50.0 50.5
Sample 2: TQLPPAYTNSFTR
y9+ y10+ y10++ y7+ y8+ b3+ y9++ y11+
72 73 74 75
Sample 2: GIYQTSNFR
y5+ y7+ y6+ y4+ y3+ y2+ b3+ y7++
4×10^6
3×10^5
61.5 61.8 62.1 62.4
AQUA peptide
SARS-CoV-2 peptide
Fragment ions
Retention time (min)

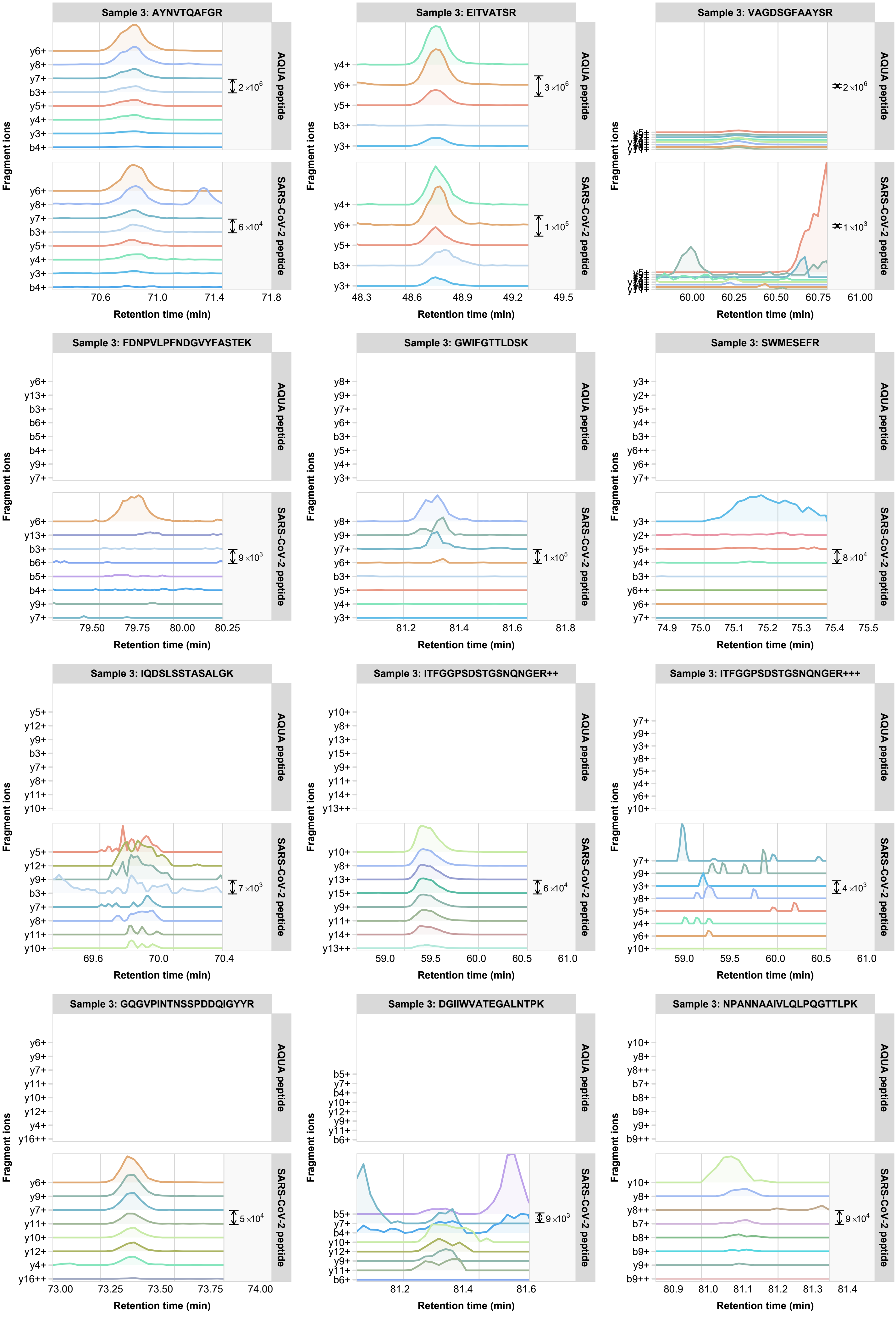

Sample 3: AYNVTQAFGR
AQUA peptide
SARS-CoV-2 peptide
y6+ y8+ y7+ b3+ y5+ y4+ y3+ b4+
2×10^6
6×10^4
70.6 71.0 71.4 71.8
Sample 3: EITVATSR
y4+ y6+ y5+ b3+ y3+
3×10^6
1×10^5
48.3 48.6 48.9 49.2 49.5
Sample 3: VAGDSGFAAYSR
2×10^6
1×10^3
60.00 60.25 60.50 60.75 61.00
Sample 3: FDNPVLPFNDGVYFASTEK
y6+ y13+ b3+ b6+ b5+ b4+ y9+ y7+
9×10^3
79.50 79.75 80.00 80.25
Sample 3: GWIFGTTLDSK
y8+ y9+ y7+ y6+ b3+ y5+ y4+ y3+
1×10^5
81.2 81.4 81.6 81.8
Sample 3: SWMESEFR
y3+ y2+ y5+ y4+ b3+ y6++ y6+ y7+
8×10^4
74.9 75.0 75.1 75.2 75.3 75.4 75.5
Sample 3: IQDSLSSTASALGK
y5+ y12+ y9+ b3+ y7+ y8+ y11+ y10+
7×10^3
69.6 70.0 70.4
Sample 3: ITFGGPSDSTGSNQNGER++
y10+ y8+ y13+ y15+ y9+ y11+ y14+ y13++
6×10^4
59.0 59.5 60.0 60.5 61.0
Sample 3: ITFGGPSDSTGSNQNGER+++
y7+ y9+ y3+ y8+ y5+ y4+ y6+ y10+
4×10^3
59.0 59.5 60.0 60.5 61.0
Sample 3: GQGVPINTNSSPDDQIGYYR
y6+ y9+ y7+ y11+ y10+ y12+ y4+ y16++
5×10^4
73.00 73.25 73.50 73.75 74.00
Sample 3: DGIIWVATEGALNTPK
b5+ y7+ b4+ y10+ y12+ y9+ y11+ b6+
9×10^3
81.2 81.4 81.6
Sample 3: NPANNAAIVLQLPQGTTLPK
y10+ y8+ y8++ b7+ b8+ b9+ y9+ b9++
9×10^4
80.9 81.0 81.1 81.2 81.3 81.4
Fragment ions
Retention time (min)

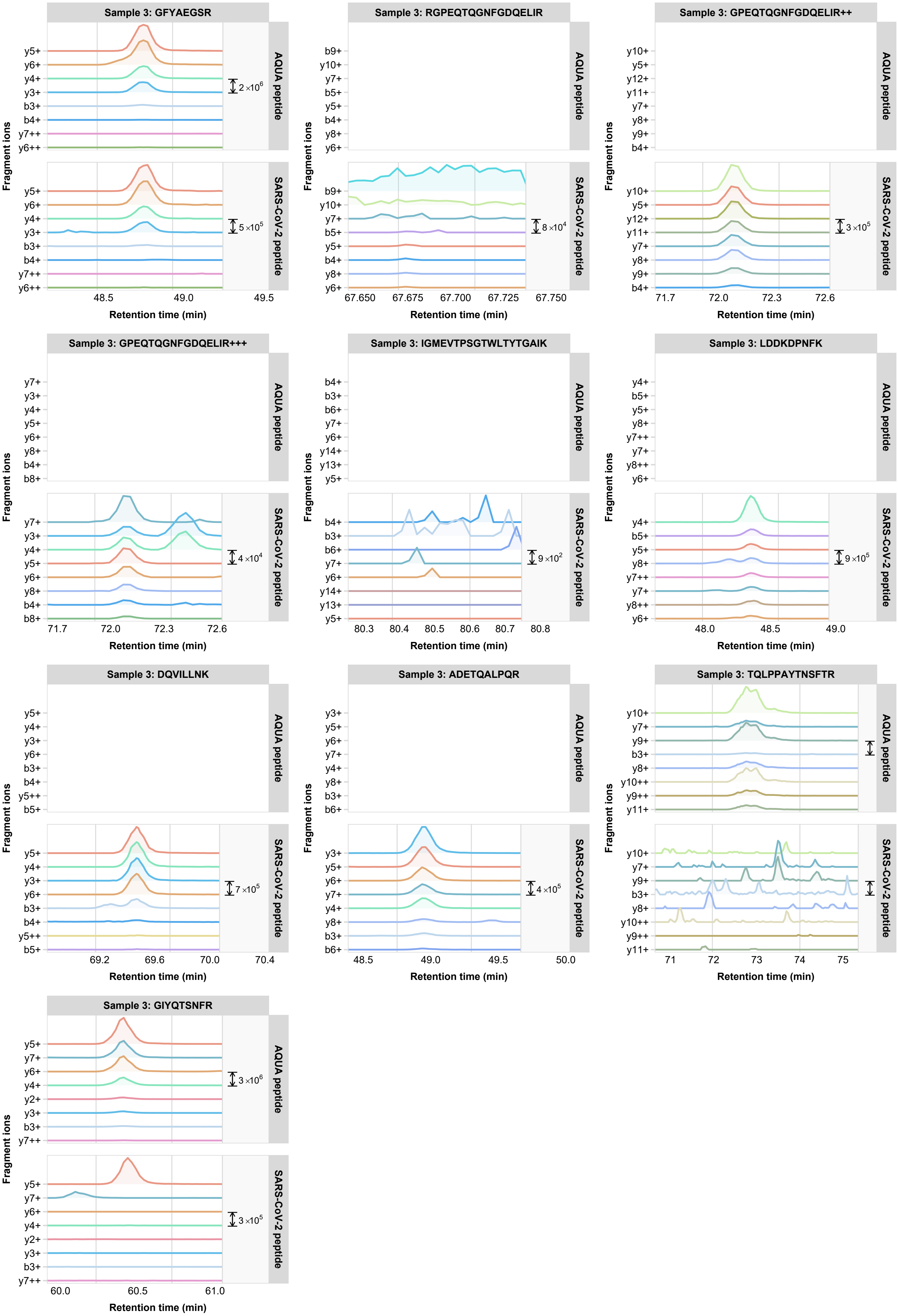

Sample 3: GFYAEGSR
AQUA peptide
y5+
y6+
y4+
y3+
b3+
b4+
y7++
y6++
2×10^6
SARS-CoV-2 peptide
5×10^5
48.5
49.0
49.5
Sample 3: RGPEQTQGNFGDQELIR
AQUA peptide
b9+
y10+
y7+
b5+
y5+
b4+
y8+
y6+
SARS-CoV-2 peptide
8×10^4
67.650
67.675
67.700
67.725
67.750
Sample 3: GPEQTQGNFGDQELIR++
AQUA peptide
y10+
y5+
y12+
y11+
y7+
y8+
y9+
b4+
SARS-CoV-2 peptide
3×10^5
71.7
72.0
72.3
72.6
Sample 3: GPEQTQGNFGDQELIR+++
AQUA peptide
y7+
y3+
y4+
y5+
y6+
y8+
b4+
b8+
SARS-CoV-2 peptide
4×10^4
71.7
72.0
72.3
72.6
Sample 3: IGMEVTPSGTWLTYTGAIK
AQUA peptide
b4+
b3+
b6+
y7+
y6+
y14+
y13+
y5+
SARS-CoV-2 peptide
9×10^2
80.3
80.4
80.5
80.6
80.7
80.8
Sample 3: LDDKDPNFK
AQUA peptide
y4+
b5+
y5+
y8+
y7++
y7+
y8++
y6+
SARS-CoV-2 peptide
9×10^5
48.0
48.5
49.0
Sample 3: DQVILLNK
AQUA peptide
y5+
y4+
y3+
y6+
b3+
b4+
y5++
b5+
SARS-CoV-2 peptide
7×10^5
69.2
69.6
70.0
70.4
Sample 3: ADETQALPQR
AQUA peptide
y3+
y5+
y6+
y7+
y4+
y8+
b3+
b6+
SARS-CoV-2 peptide
4×10^5
48.5
49.0
49.5
50.0
Sample 3: TQLPPAYTNSFTR
AQUA peptide
y10+
y7+
y9+
b3+
y8+
y10++
y9++
y11+
SARS-CoV-2 peptide
71
72
73
74
75
Sample 3: GIYQTSNFR
AQUA peptide
y5+
y7+
y6+
y4+
y2+
y3+
b3+
y7++
3×10^6
SARS-CoV-2 peptide
3×10^5
60.0
60.5
61.0
Fragment ions
Retention time (min)

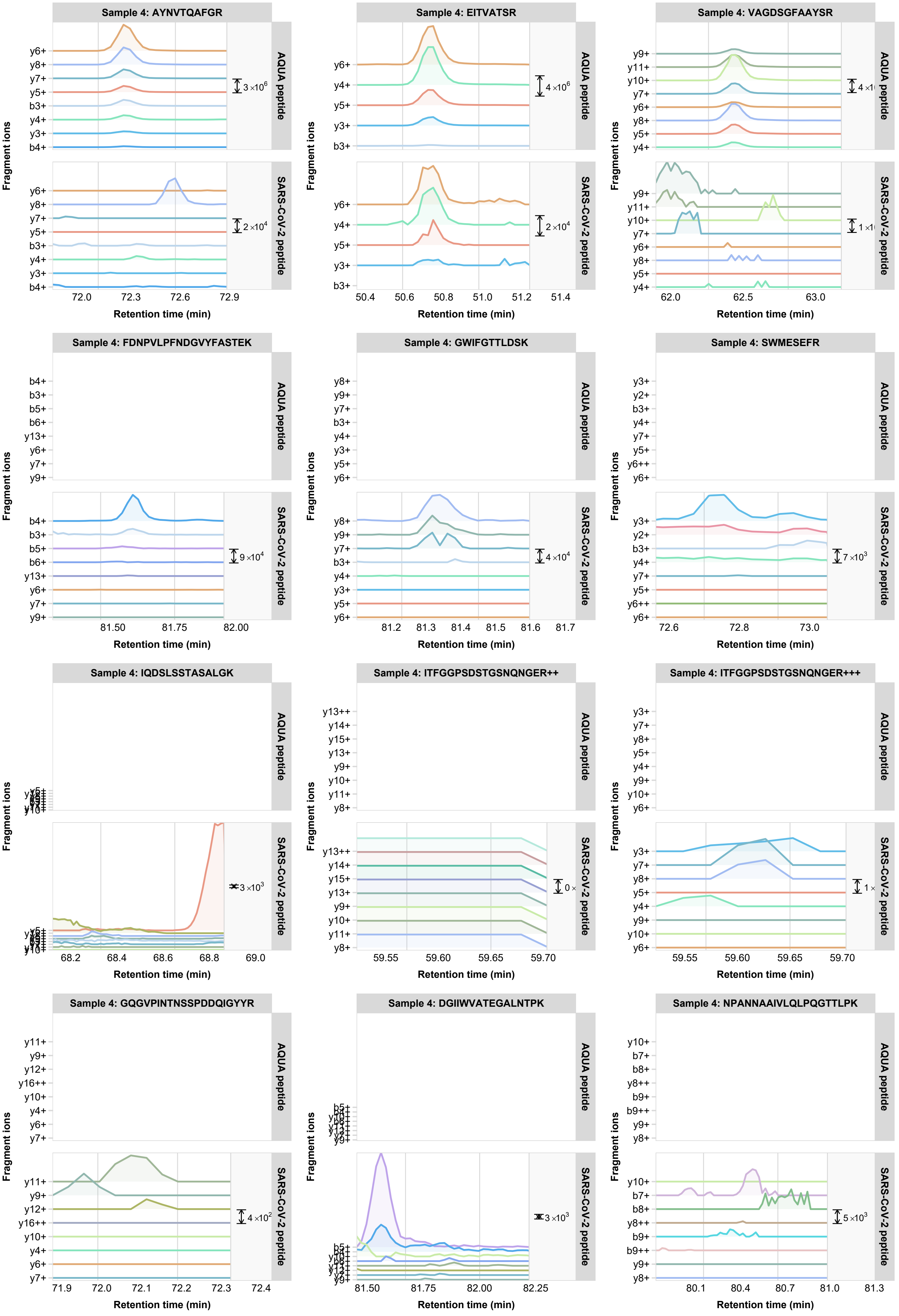

Sample 4: AYNVTQAFGR
AQUA peptide
SARS-CoV-2 peptide
y6+ y8+ y7+ y5+ b3+ y4+ y3+ b4+
3×10^6
2×10^4
72.0 72.3 72.6 72.9
Sample 4: EITVATSR
y6+ y4+ y5+ y3+ b3+
4×10^6
2×10^4
50.4 50.6 50.8 51.0 51.2 51.4
Sample 4: VAGDSGFAAYSR
y9+ y11+ y10+ y7+ y6+ y8+ y5+ y4+
62.0 62.5 63.0
Sample 4: FDNPVLPFNDGVYFASTEK
b4+ b3+ b5+ b6+ y13+ y6+ y7+ y9+
9×10^4
81.50 81.75 82.00
Sample 4: GWIFGTTLDSK
y8+ y9+ y7+ b3+ y4+ y3+ y5+ y6+
4×10^4
81.2 81.3 81.4 81.5 81.6 81.7
Sample 4: SWMESEFR
y3+ y2+ b3+ y4+ y7+ y5+ y6++ y6+
7×10^3
72.6 72.8 73.0
Sample 4: IQDSLSSTASALGK
3×10^3
68.2 68.4 68.6 68.8 69.0
Sample 4: ITFGGPSDSTGSNQNGER++
y13++ y14+ y15+ y13+ y9+ y10+ y11+ y8+
59.55 59.60 59.65 59.70
Sample 4: ITFGGPSDSTGSNQNGER+++
y3+ y7+ y8+ y5+ y4+ y9+ y10+ y6+
59.55 59.60 59.65 59.70
Sample 4: GQGVPINTNSSPDDQIGYYR
y11+ y9+ y12+ y16++ y10+ y4+ y6+ y7+
4×10^2
71.9 72.0 72.1 72.2 72.3 72.4
Sample 4: DGIIWVATEGALNTPK
3×10^3
81.50 81.75 82.00 82.25
Sample 4: NPANNAAIVLQLPQGTTLPK
y10+ b7+ b8+ y8++ b9+ b9++ y9+ y8+
5×10^3
80.1 80.4 80.7 81.0 81.3
Fragment ions
Retention time (min)

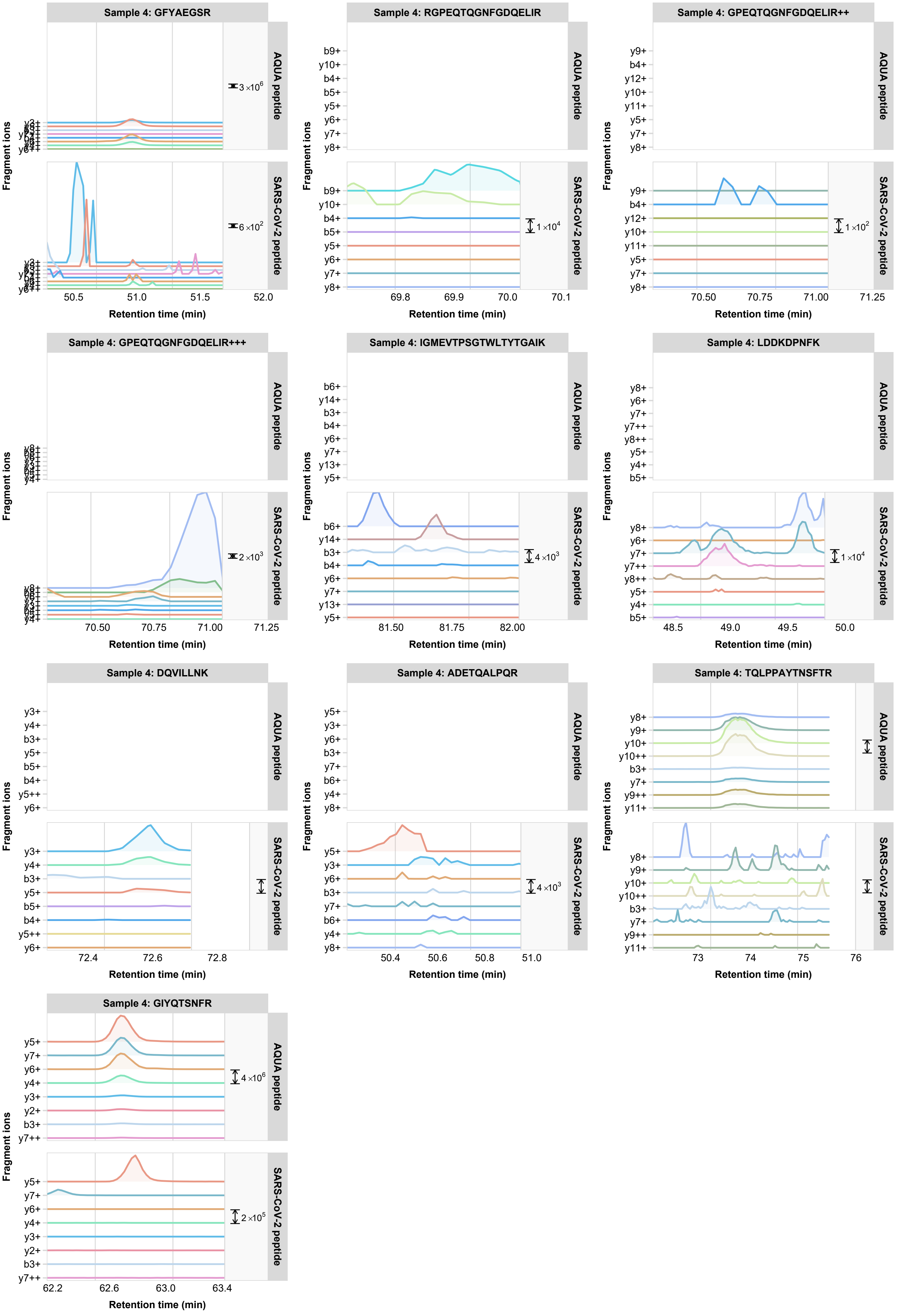

Sample 4: GFYAEGSR
AQUA peptide
SARS-CoV-2 peptide
3 ×10^6
6 ×10^2
Fragment ions
Retention time (min)
50.5
51.0
51.5
52.0
Sample 4: RGPEQTQGNFGDQELIR
b9+
y10+
b4+
b5+
y5+
y6+
y7+
y8+
1 ×10^4
69.8
69.9
70.0
70.1
Sample 4: GPEQTQGNFGDQELIR++
y9+
b4+
y12+
y10+
y11+
y5+
y7+
y8+
1 ×10^2
70.50
70.75
71.00
71.25
Sample 4: GPEQTQGNFGDQELIR+++
2 ×10^3
70.50
70.75
71.00
71.25
Sample 4: IGMEVTPSGTWLTYTGAIK
b6+
y14+
b3+
b4+
y6+
y7+
y13+
y5+
4 ×10^3
81.50
81.75
82.00
Sample 4: LDDKDPNFK
y8+
y6+
y7+
y7++
y8++
y5+
y4+
b5+
1 ×10^4
48.5
49.0
49.5
50.0
Sample 4: DQVILLNK
y3+
y4+
b3+
y5+
b5+
b4+
y5++
y6+
72.4
72.6
72.8
Sample 4: ADETQALPQR
y5+
y3+
y6+
b3+
y7+
b6+
y4+
y8+
4 ×10^3
50.4
50.6
50.8
51.0
Sample 4: TQLPPAYTNSFTR
y8+
y9+
y10+
y10++
b3+
y7+
y9++
y11+
73
74
75
76
Sample 4: GIYQTSNFR
y5+
y7+
y6+
y4+
y3+
y2+
b3+
y7++
4 ×10^6
2 ×10^5
62.2
62.6
63.0
63.4

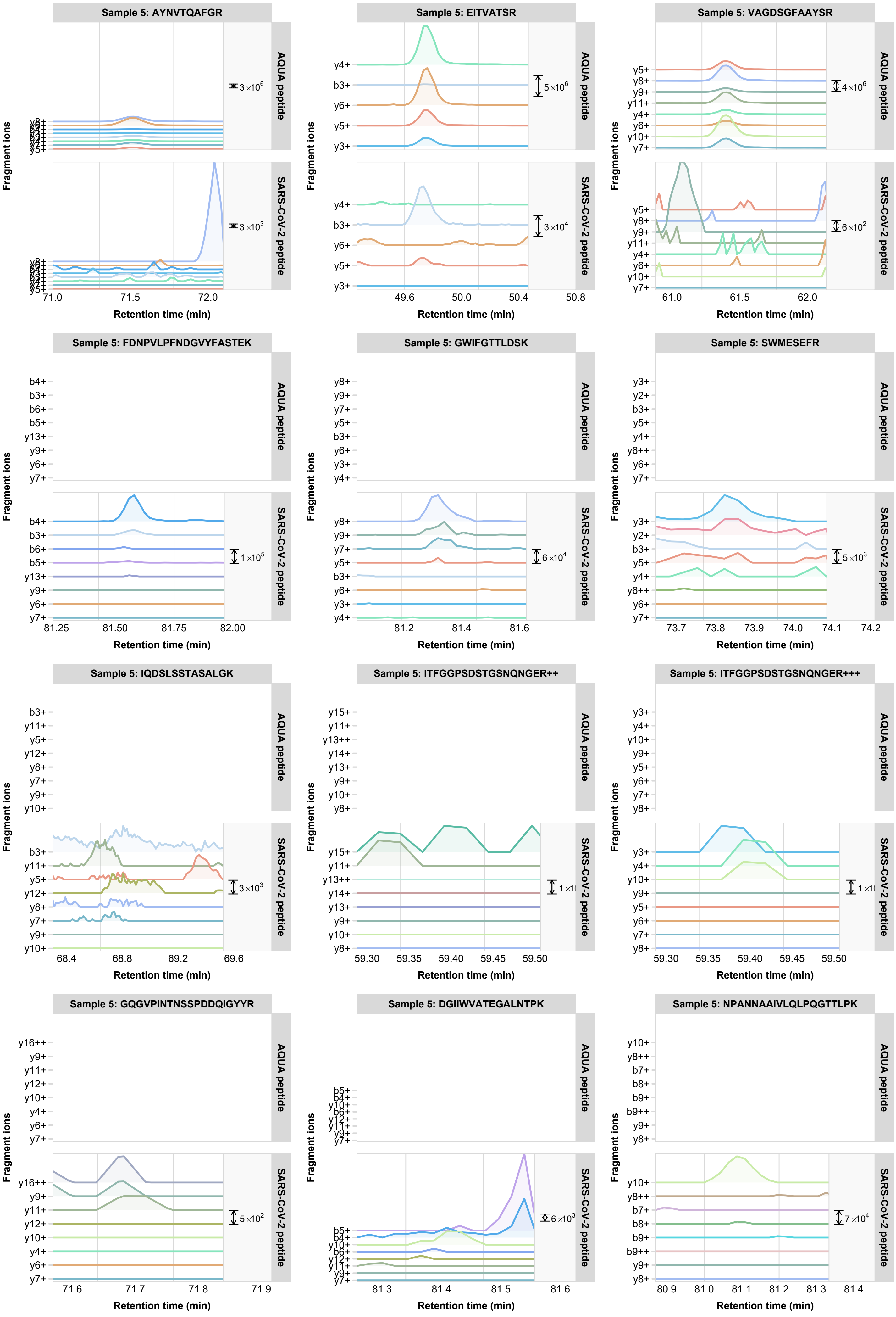

Sample 5: AYNVTQAFGR
Sample 5: EITVATSR
Sample 5: VAGDSGFAAYSR
AQUA peptide
SARS-CoV-2 peptide
Fragment ions
Retention time (min)
3×10^6
3×10^3
71.0 71.5 72.0
y4+ b3+ y6+ y5+ y3+
5×10^6
3×10^4
49.6 50.0 50.4 50.8
y5+ y8+ y9+ y11+ y4+ y6+ y10+ y7+
4×10^6
6×10^2
61.0 61.5 62.0
Sample 5: FDNPVLPFNDGVYFASTEK
b4+ b3+ b6+ b5+ y13+ y9+ y6+ y7+
1×10^5
81.25 81.50 81.75 82.00
Sample 5: GWIFGTTLDSK
y8+ y9+ y7+ y5+ b3+ y6+ y3+ y4+
6×10^4
81.2 81.4 81.6
Sample 5: SWMESEFR
y3+ y2+ b3+ y5+ y4+ y6++ y6+ y7+
5×10^3
73.7 73.8 73.9 74.0 74.1 74.2
Sample 5: IQDSLSSTASALGK
b3+ y11+ y5+ y12+ y8+ y7+ y9+ y10+
3×10^3
68.4 68.8 69.2 69.6
Sample 5: ITFGGPSDSTGSNQNGER++
y15+ y11+ y13++ y14+ y13+ y9+ y10+ y8+
59.30 59.35 59.40 59.45 59.50
Sample 5: ITFGGPSDSTGSNQNGER+++
y3+ y4+ y10+ y9+ y5+ y6+ y7+ y8+
59.30 59.35 59.40 59.45 59.50
Sample 5: GQGVPINTNSSPDDQIGYYR
y16++ y9+ y11+ y12+ y10+ y4+ y6+ y7+
5×10^2
71.6 71.7 71.8 71.9
Sample 5: DGIIWVATEGALNTPK
b5+ b4+ y10+ b6+ y12+ y11+ y9+ y7+
6×10^3
81.3 81.4 81.5 81.6
Sample 5: NPANNAAIVLQLPQGTTLPK
y10+ y8++ b7+ b8+ b9+ b9++ y9+ y8+
7×10^4
80.9 81.0 81.1 81.2 81.3 81.4

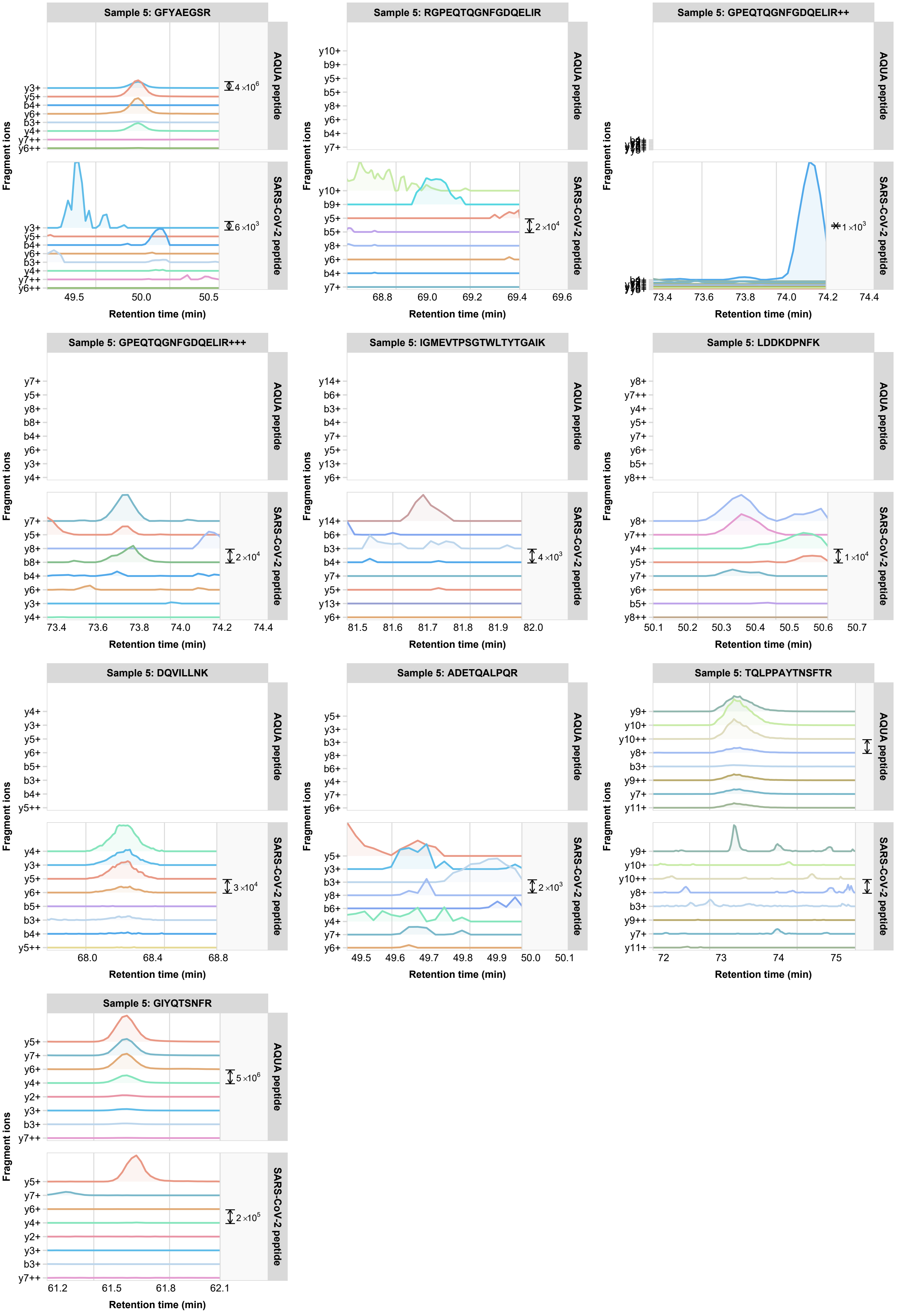

Sample 5: GFYAEGSR
AQUA peptide
SARS-CoV-2 peptide
y3+
y5+
b4+
y6+
b3+
y4+
y7++
y6++
4 ×10^6
6 ×10^3
49.5
50.0
50.5
Sample 5: RGPEQTQGNFGDQELIR
y10+
b9+
y5+
b5+
y8+
y6+
b4+
y7+
2 ×10^4
68.8
69.0
69.2
69.4
69.6
Sample 5: GPEQTQGNFGDQELIR++
1 ×10^3
73.4
73.6
73.8
74.0
74.2
74.4
Sample 5: GPEQTQGNFGDQELIR+++
y7+
y5+
y8+
b8+
b4+
y6+
y3+
y4+
2 ×10^4
73.4
73.6
73.8
74.0
74.2
74.4
Sample 5: IGMEVTPSGTWLTYTGAIK
y14+
b6+
b3+
b4+
y7+
y5+
y13+
y6+
4 ×10^3
81.5
81.6
81.7
81.8
81.9
82.0
Sample 5: LDDKDPNFK
y8+
y7++
y4+
y5+
y7+
y6+
b5+
y8++
1 ×10^4
50.1
50.2
50.3
50.4
50.5
50.6
50.7
Sample 5: DQVILLNK
y4+
y3+
y5+
y6+
b5+
b3+
b4+
y5++
3 ×10^4
68.0
68.4
68.8
Sample 5: ADETQALPQR
y5+
y3+
b3+
y8+
b6+
y4+
y7+
y6+
2 ×10^3
49.5
49.6
49.7
49.8
49.9
50.0
50.1
Sample 5: TQLPPAYTNSFTR
y9+
y10+
y10++
y8+
b3+
y9++
y7+
y11+
72
73
74
75
Sample 5: GIYQTSNFR
y5+
y7+
y6+
y4+
y2+
y3+
b3+
y7++
5 ×10^6
2 ×10^5
61.2
61.5
61.8
62.1
Fragment ions
Retention time (min)

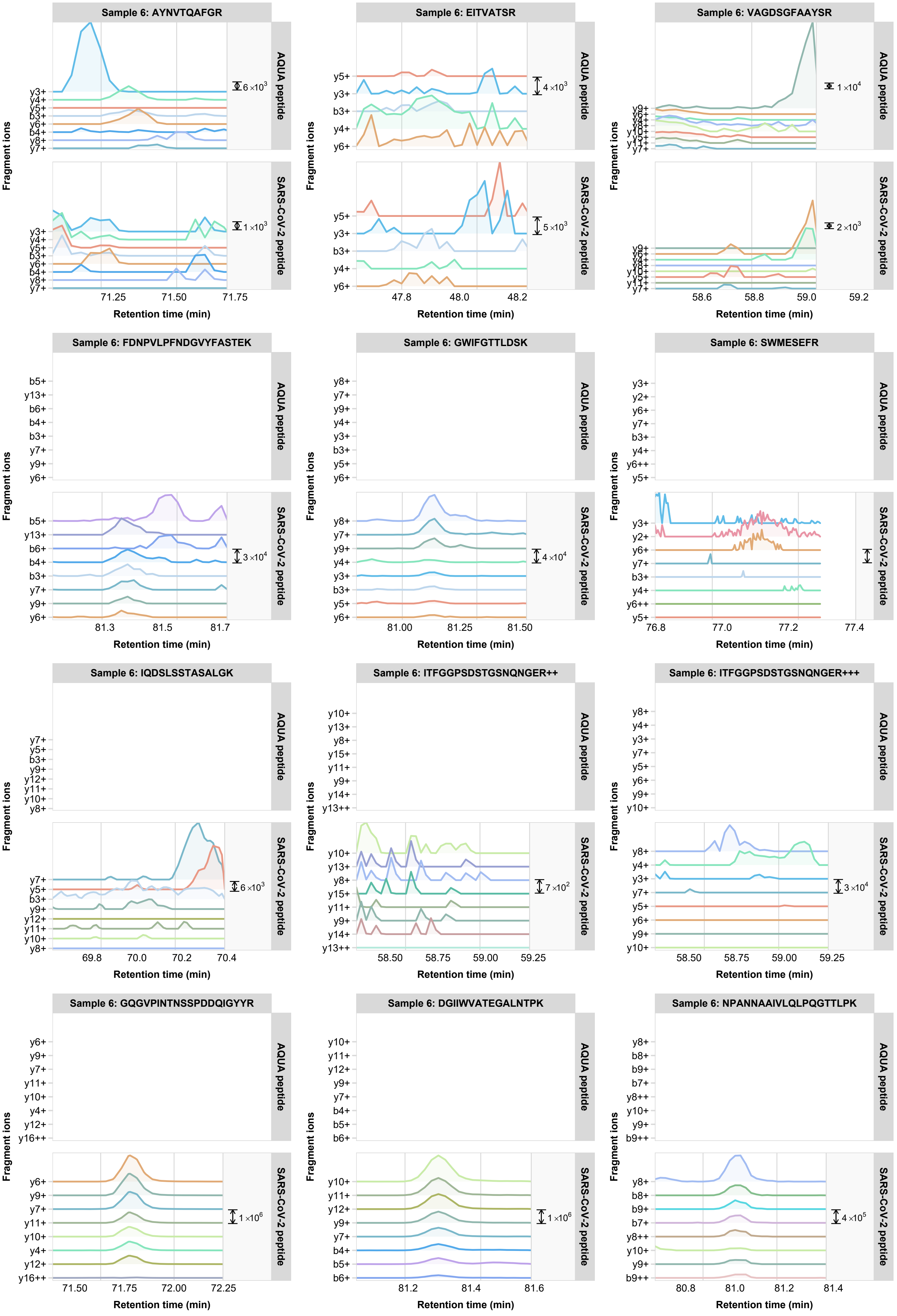
Sample 6: AYNVTQAFGR
AQUA peptide
SARS-CoV-2 peptide
y3+ y4+ y5+ b3+ y6+ b4+ y8+ y7+
6×10³
1×10³
71.25 71.50 71.75
Sample 6: EITVATSR
y5+ y3+ b3+ y4+ y6+
4×10³
5×10³
47.8 48.0 48.2
Sample 6: VAGDSGFAAYSR
y9+ y6+ y4+ y8+ y10+ y5+ y11+ y7+
1×10⁴
2×10³
58.6 58.8 59.0 59.2
Sample 6: FDNPVLPFNDGVYFASTEK
b5+ y13+ b6+ b4+ b3+ y7+ y9+ y6+
3×10⁴
81.3 81.5 81.7
Sample 6: GWIFGTTLDSK
y8+ y7+ y9+ y4+ y3+ b3+ y5+ y6+
4×10⁴
81.00 81.25 81.50
Sample 6: SWMESEFR
y3+ y2+ y6+ y7+ b3+ y4+ y6++ y5+
76.8 77.0 77.2 77.4
Sample 6: IQDSLSSTASALGK
y7+ y5+ b3+ y9+ y12+ y11+ y10+ y8+
6×10³
69.8 70.0 70.2 70.4
Sample 6: ITFGGPSDSTGSNQNGER++
y10+ y13+ y8+ y15+ y11+ y9+ y14+ y13++
7×10²
58.50 58.75 59.00 59.25
Sample 6: ITFGGPSDSTGSNQNGER+++
y8+ y4+ y3+ y7+ y5+ y6+ y9+ y10+
3×10⁴
58.50 58.75 59.00 59.25
Sample 6: GQGVPINTNSSPDDQIGYYR
y6+ y9+ y7+ y11+ y10+ y4+ y12+ y16++
1×10⁶
71.50 71.75 72.00 72.25
Sample 6: DGIIWVATEGALNTPK
y10+ y11+ y12+ y9+ y7+ b4+ b5+ b6+
1×10⁶
81.2 81.4 81.6
Sample 6: NPANNAAIVLQLPQGTTLPK
y8+ b8+ b9+ b7+ y8++ y10+ y9+ b9++
4×10⁵
80.8 81.0 81.2 81.4
Fragment ions
Retention time (min)

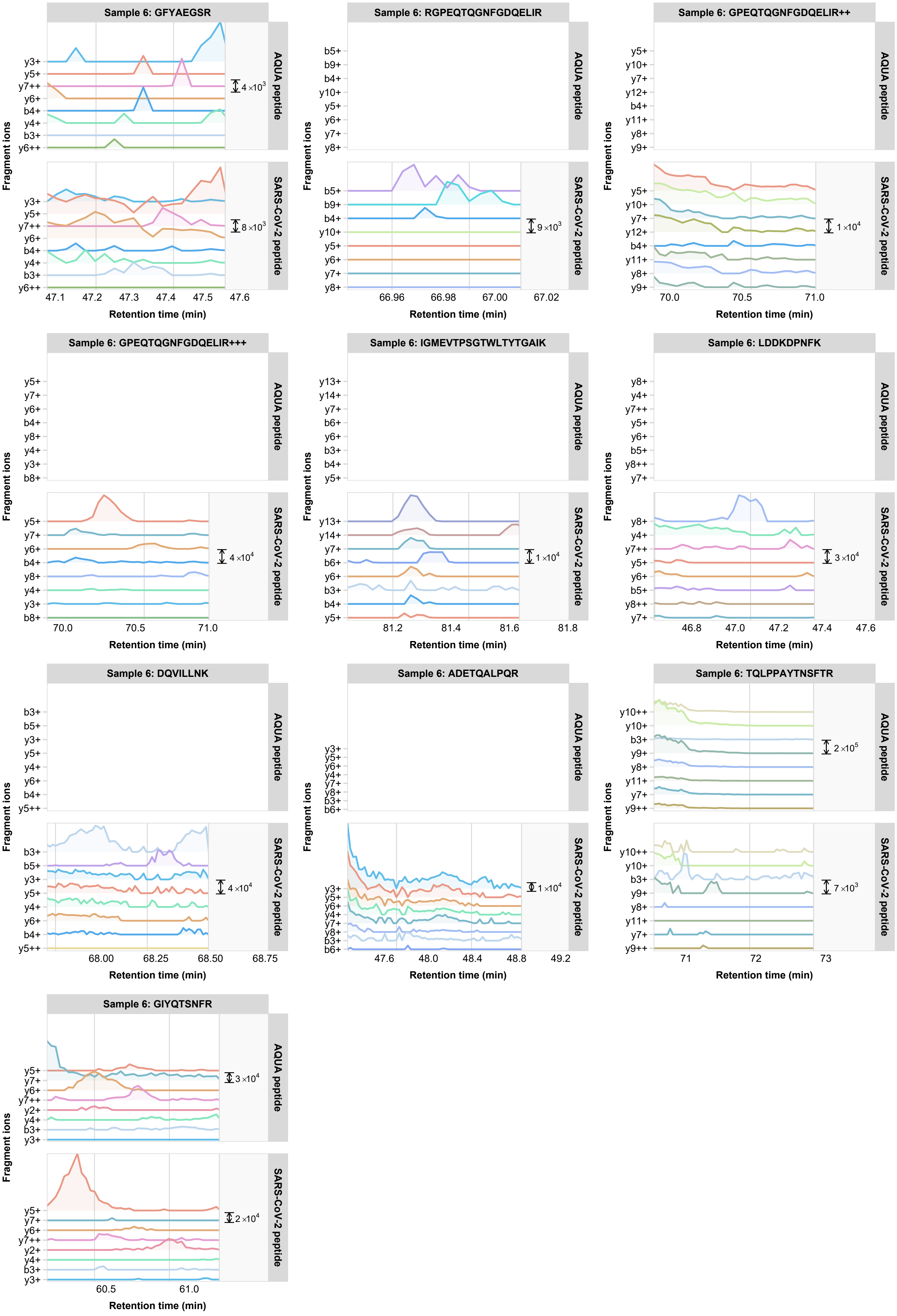

Sample 6: GFYAEGSR
AQUA peptide
SARS-CoV-2 peptide
y3+ y5+ y7++ y6+ b4+ y4+ b3+ y6++
4×10^3
8×10^3
47.1 47.2 47.3 47.4 47.5 47.6
Sample 6: RGPEQTQGNFGDQELIR
b5+ b9+ b4+ y10+ y5+ y6+ y7+ y8+
9×10^3
66.96 66.98 67.00 67.02
Sample 6: GPEQTQGNFGDQELIR++
y5+ y10+ y7+ y12+ b4+ y11+ y8+ y9+
1×10^4
70.0 70.5 71.0
Sample 6: GPEQTQGNFGDQELIR+++
y5+ y7+ y6+ b4+ y8+ y4+ y3+ b8+
4×10^4
70.0 70.5 71.0
Sample 6: IGMEVTPSGTWLTYTGAIK
y13+ y14+ y7+ b6+ y6+ b3+ b4+ y5+
1×10^4
81.2 81.4 81.6 81.8
Sample 6: LDDKDPNFK
y8+ y4+ y7++ y5+ y6+ b5+ y8++ y7+
3×10^4
46.8 47.0 47.2 47.4 47.6
Sample 6: DQVILLNK
b3+ b5+ y3+ y5+ y4+ y6+ b4+ y5++
4×10^4
68.00 68.25 68.50 68.75
Sample 6: ADETQALPQR
y3+ y5+ y6+ y4+ y7+ y8+ b3+ b6+
1×10^4
47.6 48.0 48.4 48.8 49.2
Sample 6: TQLPPAYTNSFTR
y10++ y10+ b3+ y9+ y8+ y11+ y7+ y9++
2×10^5
7×10^3
71 72 73
Sample 6: GIYQTSNFR
y5+ y7+ y6+ y7++ y2+ y4+ b3+ y3+
3×10^4
2×10^4
60.5 61.0
Fragment ions
Retention time (min)

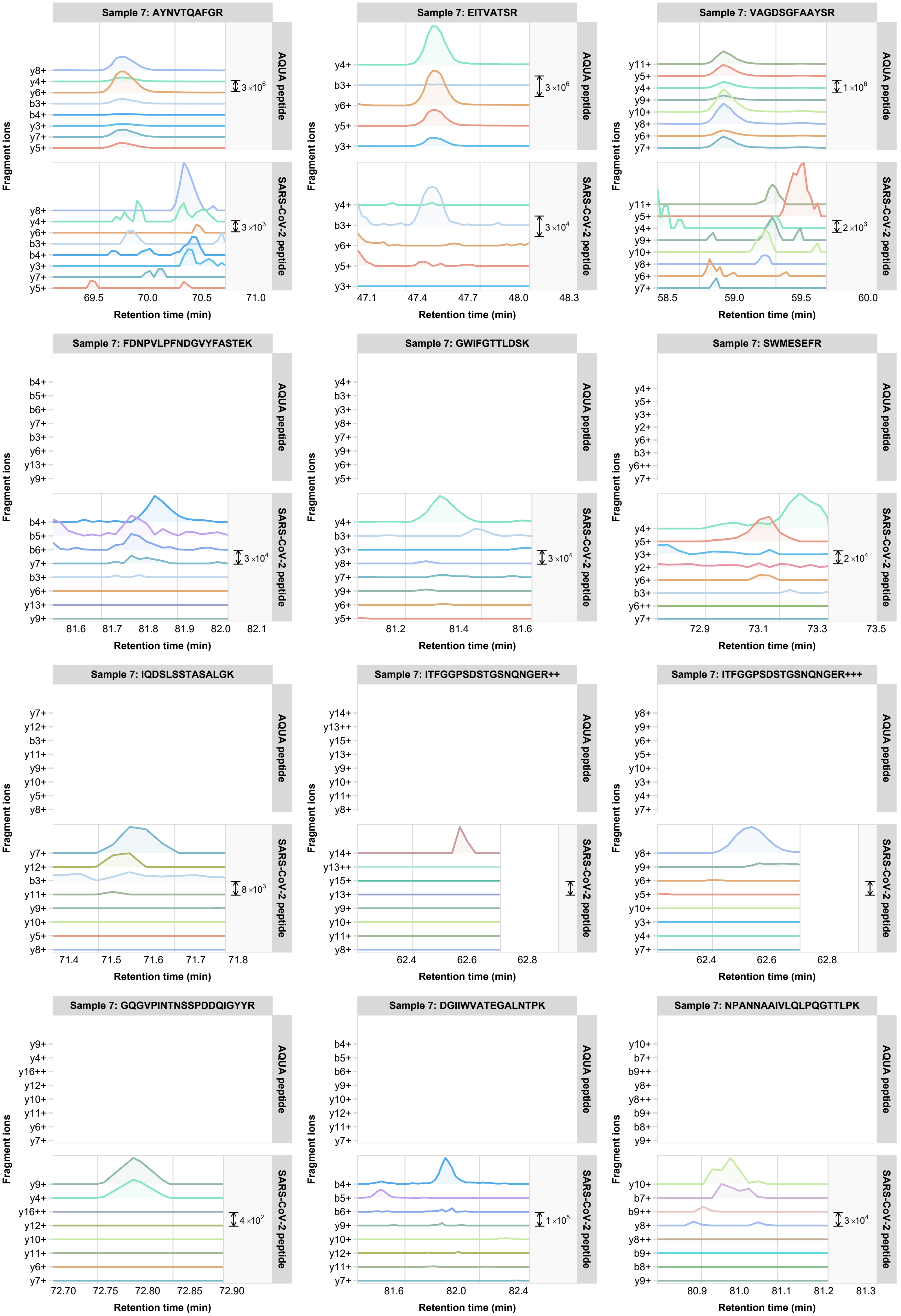
Sample 7: AYNVTQAFGR
y8+
y4+
y6+
b3+
b4+
y3+
y7+
y5+
3×10^6
3×10^3
AQUA peptide
SARS-CoV-2 peptide
69.5
70.0
70.5
71.0
Sample 7: EITVATSR
y4+
b3+
y6+
y5+
y3+
3×10^6
3×10^4
47.1
47.4
47.7
48.0
48.3
Sample 7: VAGDSGFAAYSR
y11+
y5+
y4+
y9+
y10+
y8+
y6+
y7+
1×10^6
2×10^3
58.5
59.0
59.5
60.0
Sample 7: FDNPVLPFNDGVYFASTEK
b4+
b5+
b6+
y7+
b3+
y6+
y13+
y9+
3×10^4
81.6
81.7
81.8
81.9
82.0
82.1
Sample 7: GWIFGTTLDSK
y4+
b3+
y3+
y8+
y7+
y9+
y6+
y5+
3×10^4
81.2
81.4
81.6
Sample 7: SWMESEFR
y4+
y5+
y3+
y2+
y6+
b3+
y6++
y7+
2×10^4
72.9
73.1
73.3
73.5
Sample 7: IQDSLSSTASALGK
y7+
y12+
b3+
y11+
y9+
y10+
y5+
y8+
8×10^3
71.4
71.5
71.6
71.7
71.8
Sample 7: ITFGGPSDSTGSNQNGER++
y14+
y13++
y15+
y13+
y9+
y10+
y11+
y8+
62.4
62.6
62.8
Sample 7: ITFGGPSDSTGSNQNGER+++
y8+
y9+
y6+
y5+
y10+
y3+
y4+
y7+
62.4
62.6
62.8
Sample 7: GQGVPINTNSSPDDQIGYYR
y9+
y4+
y16++
y12+
y10+
y11+
y6+
y7+
4×10^2
72.70
72.75
72.80
72.85
72.90
Sample 7: DGIIWVATEGALNTPK
b4+
b5+
b6+
y9+
y10+
y12+
y11+
y7+
1×10^5
81.6
82.0
82.4
Sample 7: NPANNAAIVLQLPQGTTLPK
y10+
b7+
b9++
y8+
y8++
b9+
b8+
y9+
3×10^4
80.9
81.0
81.1
81.2
81.3
Fragment ions
Retention time (min)

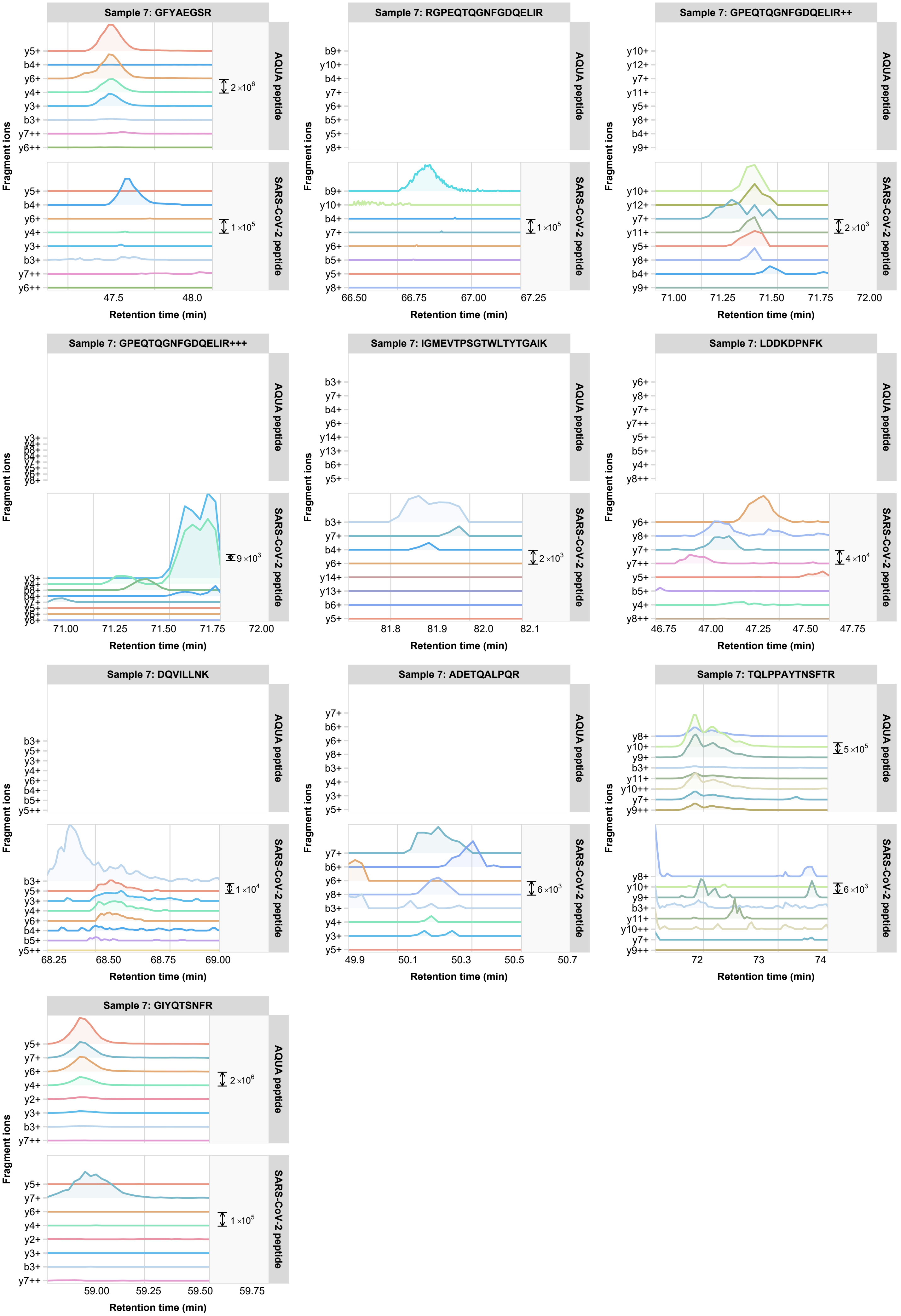

Sample 7: GFYAEGSR
AQUA peptide
SARS-CoV-2 peptide
2×10^6
1×10^5
y5+ b4+ y6+ y4+ y3+ b3+ y7++ y6++
47.5
48.0
Sample 7: RGPEQTQGNFGDQELIR
b9+ y10+ b4+ y7+ y6+ b5+ y5+ y8+
1×10^5
66.50 66.75 67.00 67.25
Sample 7: GPEQTQGNFGDQELIR++
y10+ y12+ y7+ y11+ y5+ y8+ b4+ y9+
2×10^3
71.00 71.25 71.50 71.75 72.00
Sample 7: GPEQTQGNFGDQELIR+++
9×10^3
71.00 71.25 71.50 71.75 72.00
Sample 7: IGMEVTPSGTWLTYTGAIK
b3+ y7+ b4+ y6+ y14+ y13+ b6+ y5+
2×10^3
81.8 81.9 82.0 82.1
Sample 7: LDDKDPNFK
y6+ y8+ y7+ y7++ y5+ b5+ y4+ y8++
4×10^4
46.75 47.00 47.25 47.50 47.75
Sample 7: DQVILLNK
b3+ y5+ y3+ y4+ y6+ b4+ b5+ y5++
1×10^4
68.25 68.50 68.75 69.00
Sample 7: ADETQALPQR
y7+ b6+ y6+ y8+ b3+ y4+ y3+ y5+
6×10^3
49.9 50.1 50.3 50.5 50.7
Sample 7: TQLPPAYTNSFTR
y8+ y10+ y9+ b3+ y11+ y10++ y7+ y9++
5×10^5
6×10^3
72 73 74
Sample 7: GIYQTSNFR
y5+ y7+ y6+ y4+ y2+ y3+ b3+ y7++
2×10^6
1×10^5
59.00 59.25 59.50 59.75
Fragment ions
Retention time (min)

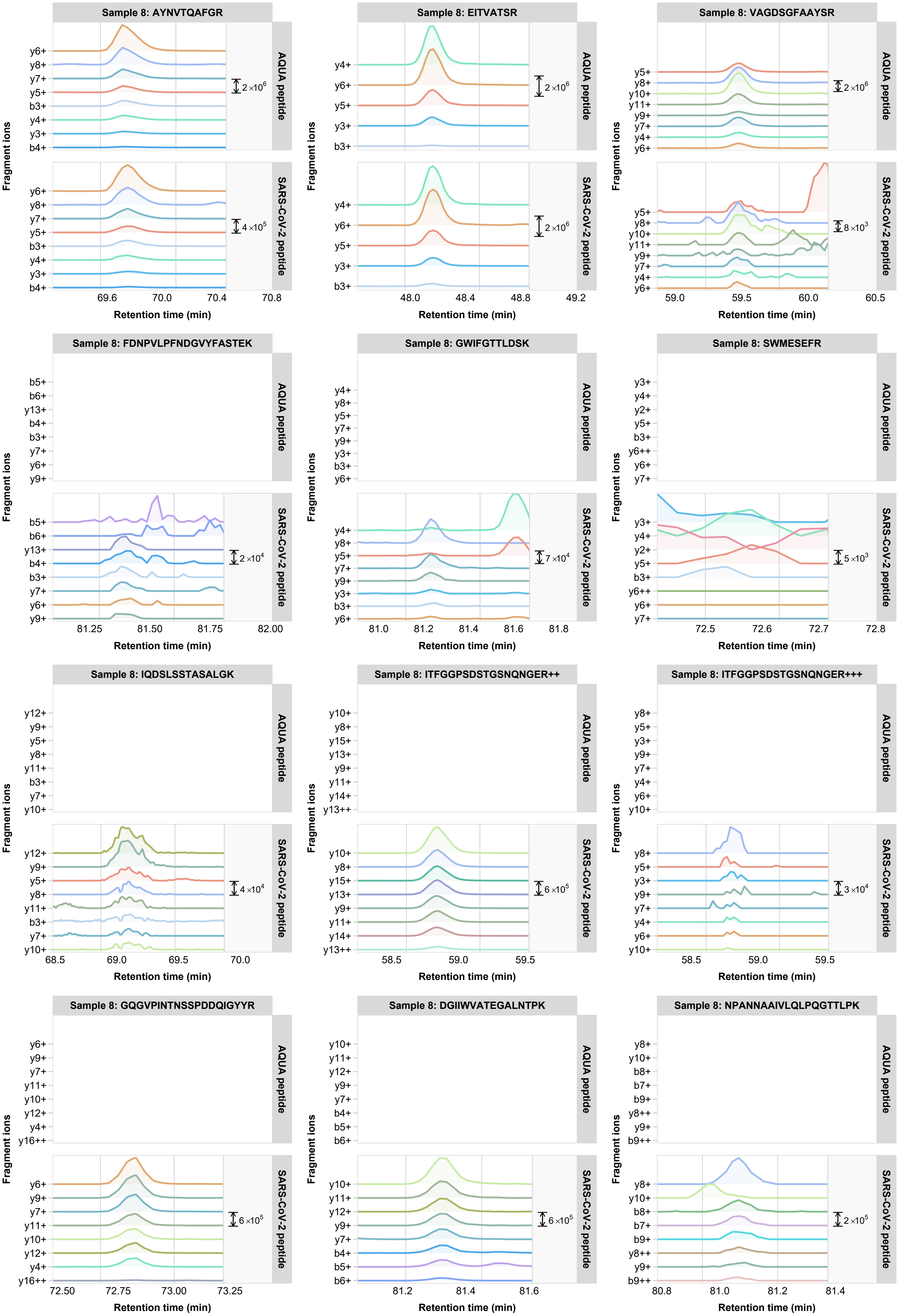
Sample 8: AYNVTQAFGR
AQUA peptide
SARS-CoV-2 peptide
y6+ y8+ y7+ y5+ b3+ y4+ y3+ b4+
2×10^6
4×10^5
69.6 70.0 70.4 70.8
Sample 8: EITVATSR
y4+ y6+ y5+ y3+ b3+
2×10^6
2×10^6
48.0 48.4 48.8 49.2
Sample 8: VAGDSGFAAYSR
y5+ y8+ y10+ y11+ y9+ y7+ y4+ y6+
2×10^6
8×10^3
59.0 59.5 60.0 60.5
Sample 8: FDNPVLPFNDGVYFASTEK
b5+ b6+ y13+ b4+ b3+ y7+ y6+ y9+
2×10^4
81.25 81.50 81.75 82.00
Sample 8: GWIFGTTLDSK
y4+ y8+ y5+ y7+ y9+ y3+ b3+ y6+
7×10^4
81.0 81.2 81.4 81.6 81.8
Sample 8: SWMESEFR
y3+ y4+ y2+ y5+ b3+ y6++ y6+ y7+
5×10^3
72.5 72.6 72.7 72.8
Sample 8: IQDSLSSTASALGK
y12+ y9+ y5+ y8+ y11+ b3+ y7+ y10+
4×10^4
68.5 69.0 69.5 70.0
Sample 8: ITFGGPSDSTGSNQNGER++
y10+ y8+ y15+ y13+ y9+ y11+ y14+ y13++
6×10^5
58.5 59.0 59.5
Sample 8: ITFGGPSDSTGSNQNGER+++
y8+ y5+ y3+ y9+ y7+ y4+ y6+ y10+
3×10^4
58.5 59.0 59.5
Sample 8: GQGVPINTNSSPDDQIGYYR
y6+ y9+ y7+ y11+ y10+ y12+ y4+ y16++
6×10^5
72.50 72.75 73.00 73.25
Sample 8: DGIIWVATEGALNTPK
y10+ y11+ y12+ y9+ y7+ b4+ b5+ b6+
6×10^5
81.2 81.4 81.6
Sample 8: NPANNAAIVLQLPQGTTLPK
y8+ y10+ b8+ b7+ b9+ y8++ y9+ b9++
2×10^5
80.8 81.0 81.2 81.4
Fragment ions
Retention time (min)

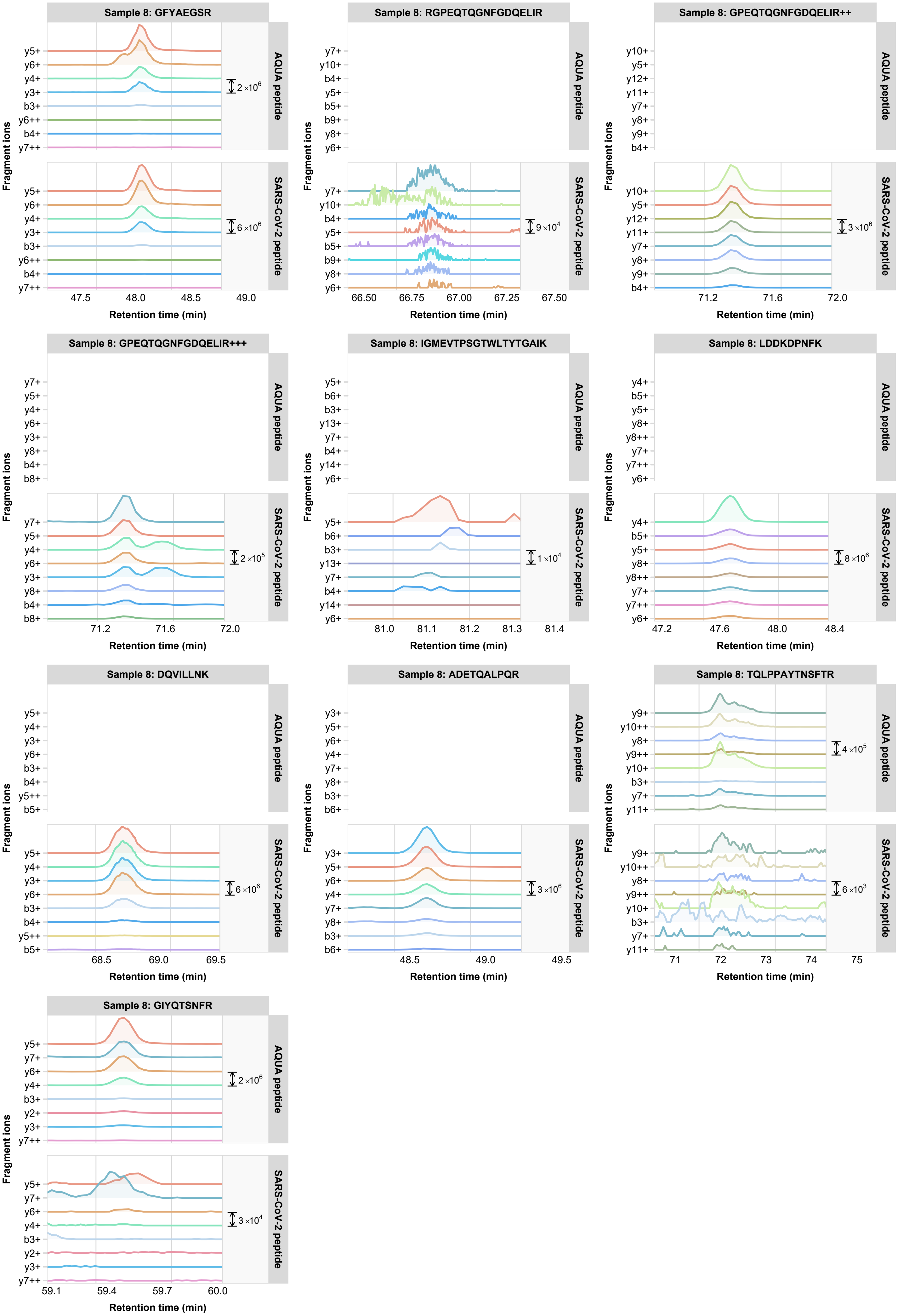

Sample 8: GFYAEGSR
AQUA peptide
y5+ y6+ y4+ y3+ b3+ y6++ b4+ y7++
2×10^6
SARS-CoV-2 peptide
y5+ y6+ y4+ y3+ b3+ y6++ b4+ y7++
6×10^6
47.5 48.0 48.5 49.0
Sample 8: RGPEQTQGNFGDQELIR
AQUA peptide
y7+ y10+ b4+ y5+ b5+ b9+ y8+ y6+
SARS-CoV-2 peptide
y7+ y10+ b4+ y5+ b5+ b9+ y8+ y6+
9×10^4
66.50 66.75 67.00 67.25 67.50
Sample 8: GPEQTQGNFGDQELIR++
AQUA peptide
y10+ y5+ y12+ y11+ y7+ y8+ y9+ b4+
SARS-CoV-2 peptide
y10+ y5+ y12+ y11+ y7+ y8+ y9+ b4+
3×10^6
71.2 71.6 72.0
Sample 8: GPEQTQGNFGDQELIR+++
AQUA peptide
y7+ y5+ y4+ y6+ y3+ y8+ b4+ b8+
SARS-CoV-2 peptide
y7+ y5+ y4+ y6+ y3+ y8+ b4+ b8+
2×10^5
71.2 71.6 72.0
Sample 8: IGMEVTPSGTWLTYTGAIK
AQUA peptide
y5+ b6+ b3+ y13+ y7+ b4+ y14+ y6+
SARS-CoV-2 peptide
y5+ b6+ b3+ y13+ y7+ b4+ y14+ y6+
1×10^4
81.0 81.1 81.2 81.3 81.4
Sample 8: LDDKDPNFK
AQUA peptide
y4+ b5+ y5+ y8+ y8++ y7+ y7++ y6+
SARS-CoV-2 peptide
y4+ b5+ y5+ y8+ y8++ y7+ y7++ y6+
8×10^6
47.2 47.6 48.0 48.4
Sample 8: DQVILLNK
AQUA peptide
y5+ y4+ y3+ y6+ b3+ b4+ y5++ b5+
SARS-CoV-2 peptide
y5+ y4+ y3+ y6+ b3+ b4+ y5++ b5+
6×10^6
68.5 69.0 69.5
Sample 8: ADETQALPQR
AQUA peptide
y3+ y5+ y6+ y4+ y7+ y8+ b3+ b6+
SARS-CoV-2 peptide
y3+ y5+ y6+ y4+ y7+ y8+ b3+ b6+
3×10^6
48.5 49.0 49.5
Sample 8: TQLPPAYTNSFTR
AQUA peptide
y9+ y10++ y8+ y9++ y10+ b3+ y7+ y11+
4×10^5
SARS-CoV-2 peptide
y9+ y10++ y8+ y9++ y10+ b3+ y7+ y11+
6×10^3
71 72 73 74 75
Sample 8: GIYQTSNFR
AQUA peptide
y5+ y7+ y6+ y4+ b3+ y2+ y3+ y7++
2×10^6
SARS-CoV-2 peptide
y5+ y7+ y6+ y4+ b3+ y2+ y3+ y7++
3×10^4
59.1 59.4 59.7 60.0
Fragment ions
Retention time (min)

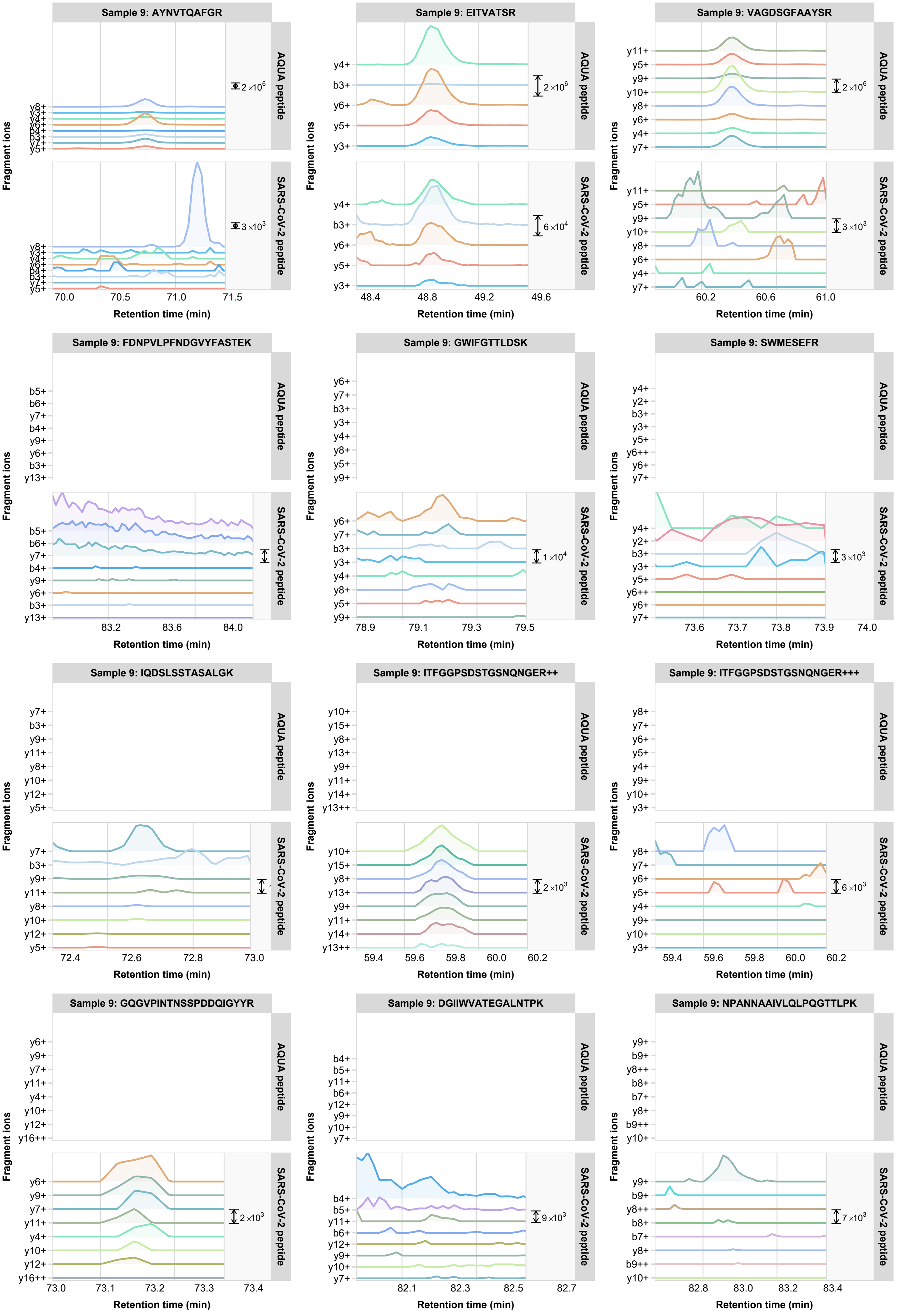
Sample 9: AYNVTQAFGR
y8+ y3+ y4+ y6+ b4+ b3+ y7+ y5+
2×10^6
3×10^3
70.0 70.5 71.0 71.5
Sample 9: EITVATSR
y4+ b3+ y6+ y5+ y3+
2×10^6
6×10^4
48.4 48.8 49.2 49.6
Sample 9: VAGDSGFAAYSR
y11+ y5+ y9+ y10+ y8+ y6+ y4+ y7+
2×10^6
3×10^3
60.2 60.6 61.0
Sample 9: FDNPVLPFNDGVYFASTEK
b5+ b6+ y7+ b4+ y9+ y6+ b3+ y13+
83.2 83.6 84.0
Sample 9: GWIFGTTLDSK
y6+ y7+ b3+ y3+ y4+ y8+ y5+ y9+
1×10^4
78.9 79.1 79.3 79.5
Sample 9: SWMESEFR
y4+ y2+ b3+ y3+ y5+ y6++ y6+ y7+
3×10^3
73.6 73.7 73.8 73.9 74.0
Sample 9: IQDSLSSTASALGK
y7+ b3+ y9+ y11+ y8+ y10+ y12+ y5+
72.4 72.6 72.8 73.0
Sample 9: ITFGGPSDSTGSNQNGER++
y10+ y15+ y8+ y13+ y9+ y11+ y14+ y13++
2×10^3
59.4 59.6 59.8 60.0 60.2
Sample 9: ITFGGPSDSTGSNQNGER+++
y8+ y7+ y6+ y5+ y4+ y9+ y10+ y3+
6×10^3
59.4 59.6 59.8 60.0 60.2
Sample 9: GQGVPINTNSSPDDQIGYYR
y6+ y9+ y7+ y11+ y4+ y10+ y12+ y16++
2×10^3
73.0 73.1 73.2 73.3 73.4
Sample 9: DGIIWVATEGALNTPK
b4+ b5+ y11+ b6+ y12+ y9+ y10+ y7+
9×10^3
82.1 82.3 82.5 82.7
Sample 9: NPANNAAIVLQLPQGTTLPK
y9+ b9+ y8++ b8+ b7+ y8+ b9++ y10+
7×10^3
82.8 83.0 83.2 83.4
AQUA peptide
SARS-CoV-2 peptide
Fragment ions
Retention time (min)

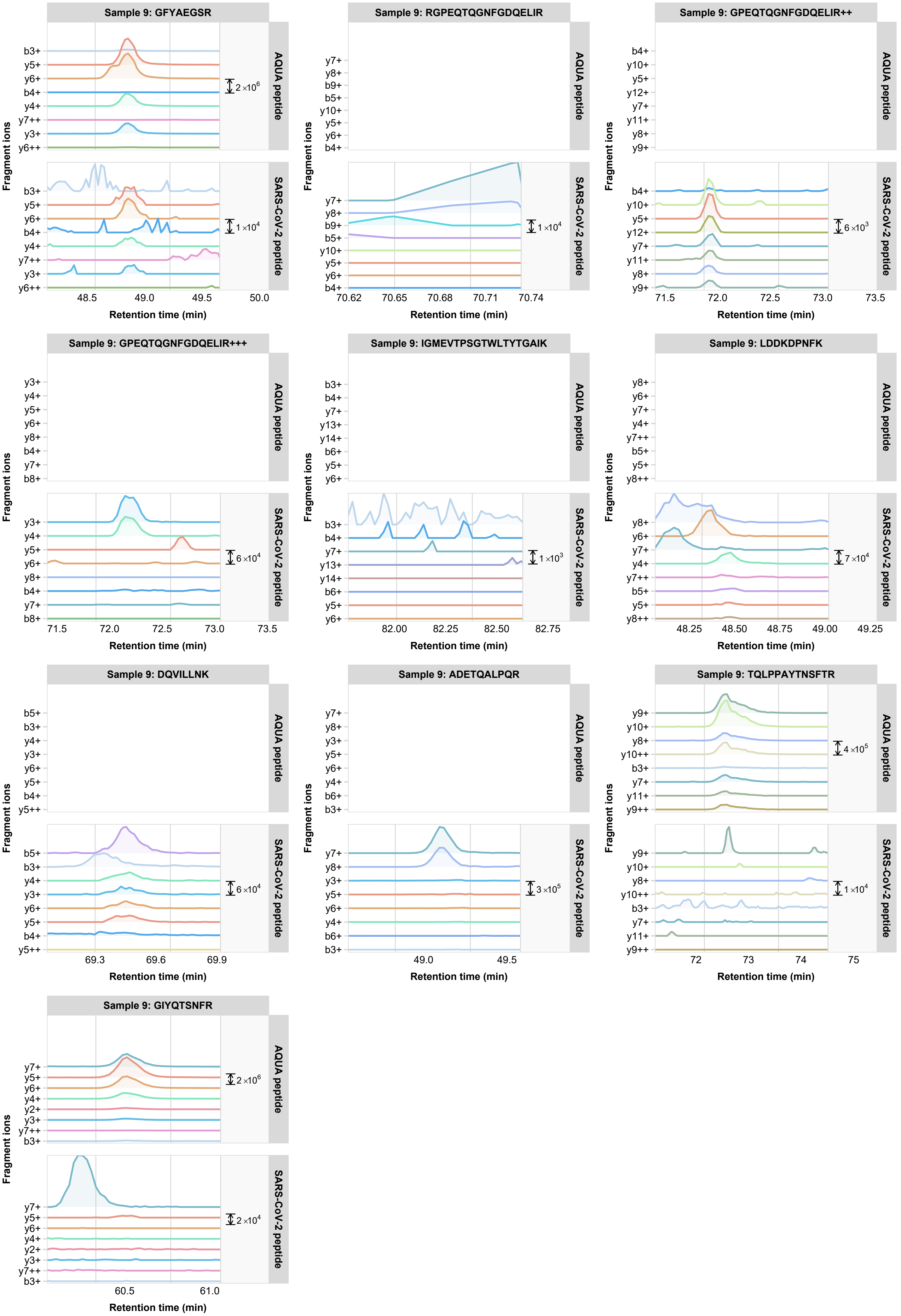

Sample 9: GFYAEGSR
AQUA peptide
SARS-CoV-2 peptide
b3+ y5+ y6+ b4+ y4+ y7++ y3+ y6++
2 ×10^6
1 ×10^4
48.5 49.0 49.5 50.0
Sample 9: RGPEQTQGNFGDQELIR
y7+ y8+ b9+ b5+ y10+ y5+ y6+ b4+
1 ×10^4
70.62 70.65 70.68 70.71 70.74
Sample 9: GPEQTQGNFGDQELIR++
b4+ y10+ y5+ y12+ y7+ y11+ y8+ y9+
6 ×10^3
71.5 72.0 72.5 73.0 73.5
Sample 9: GPEQTQGNFGDQELIR+++
y3+ y4+ y5+ y6+ y8+ b4+ y7+ b8+
6 ×10^4
71.5 72.0 72.5 73.0 73.5
Sample 9: IGMEVTPSGTWLTYTGAIK
b3+ b4+ y7+ y13+ y14+ b6+ y5+ y6+
1 ×10^3
82.00 82.25 82.50 82.75
Sample 9: LDDKDPNFK
y8+ y6+ y7+ y4+ y7++ b5+ y5+ y8++
7 ×10^4
48.25 48.50 48.75 49.00 49.25
Sample 9: DQVILLNK
b5+ b3+ y4+ y3+ y6+ y5+ b4+ y5++
6 ×10^4
69.3 69.6 69.9
Sample 9: ADETQALPQR
y7+ y8+ y3+ y5+ y6+ y4+ b6+ b3+
3 ×10^5
49.0 49.5
Sample 9: TQLPPAYTNSFTR
y9+ y10+ y8+ y10++ b3+ y7+ y11+ y9++
4 ×10^5
1 ×10^4
72 73 74 75
Sample 9: GIYQTSNFR
y7+ y5+ y6+ y4+ y2+ y3+ y7++ b3+
2 ×10^6
2 ×10^4
60.5 61.0
Fragment ions
Retention time (min)

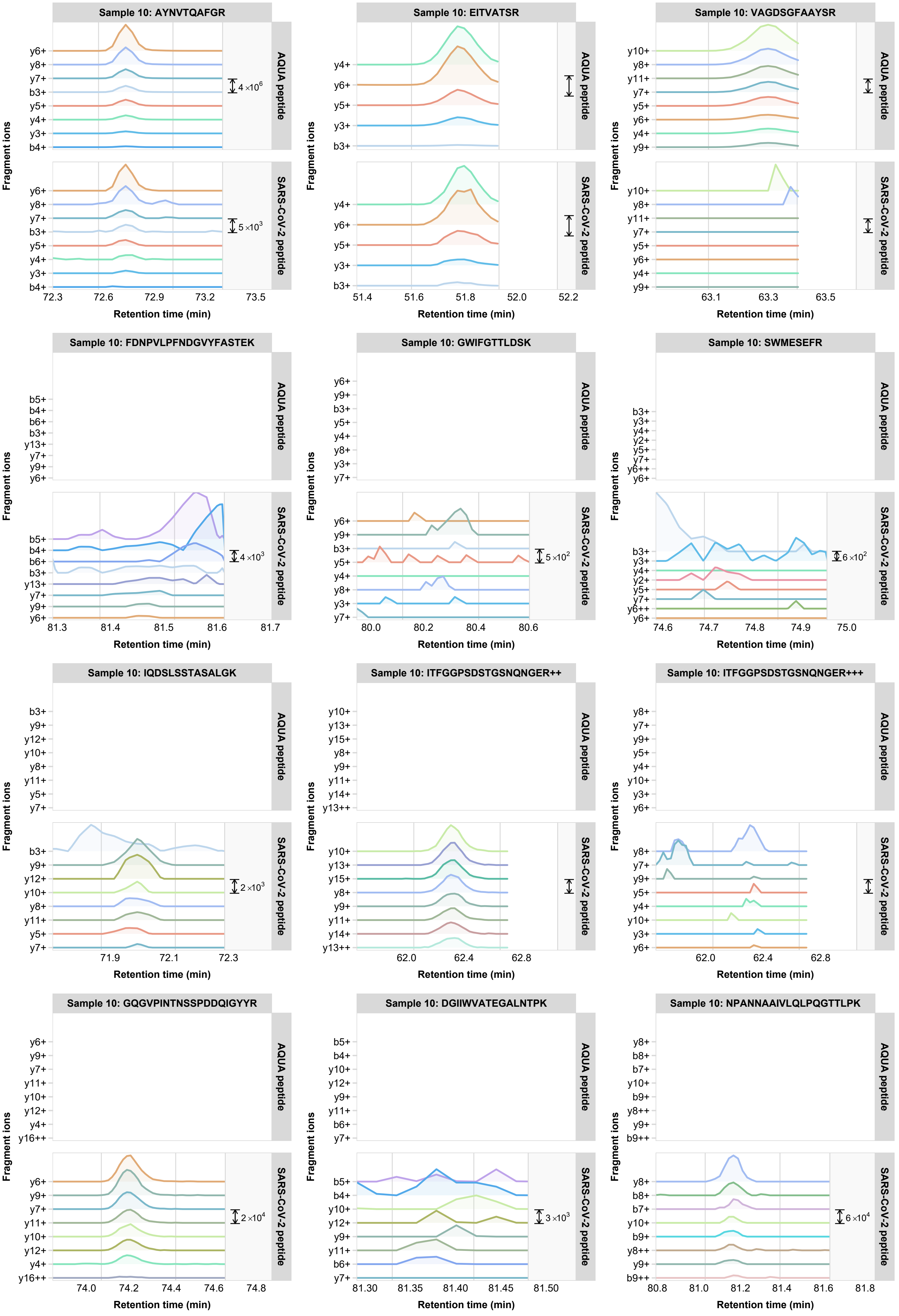
Sample 10: AYNVTQAFGR
AQUA peptide
SARS-CoV-2 peptide
y6+ y8+ y7+ b3+ y5+ y4+ y3+ b4+
4×10^6
5×10^3
72.3 72.6 72.9 73.2 73.5
Sample 10: EITVATSR
y4+ y6+ y5+ y3+ b3+
51.4 51.6 51.8 52.0 52.2
Sample 10: VAGDSGFAAYSR
y10+ y8+ y11+ y7+ y5+ y6+ y4+ y9+
63.1 63.3 63.5
Sample 10: FDNPVLPFNDGVYFASTEK
b5+ b4+ b6+ b3+ y13+ y7+ y9+ y6+
4×10^3
81.3 81.4 81.5 81.6 81.7
Sample 10: GWIFGTTLDSK
y6+ y9+ b3+ y5+ y4+ y8+ y3+ y7+
5×10^2
80.0 80.2 80.4 80.6
Sample 10: SWMESEFR
b3+ y3+ y4+ y2+ y5+ y7+ y6++ y6+
6×10^2
74.6 74.7 74.8 74.9 75.0
Sample 10: IQDSLSSTASALGK
b3+ y9+ y12+ y10+ y8+ y11+ y5+ y7+
2×10^3
71.9 72.1 72.3
Sample 10: ITFGGPSDSTGSNQNGER++
y10+ y13+ y15+ y8+ y9+ y11+ y14+ y13++
62.0 62.4 62.8
Sample 10: ITFGGPSDSTGSNQNGER+++
y8+ y7+ y9+ y5+ y4+ y10+ y3+ y6+
62.0 62.4 62.8
Sample 10: GQGVPINTNSSPDDQIGYYR
y6+ y9+ y7+ y11+ y10+ y12+ y4+ y16++
2×10^4
74.0 74.2 74.4 74.6 74.8
Sample 10: DGIIWVATEGALNTPK
b5+ b4+ y10+ y12+ y9+ y11+ b6+ y7+
3×10^3
81.30 81.35 81.40 81.45 81.50
Sample 10: NPANNAAIVLQLPQGTTLPK
y8+ b8+ b7+ y10+ b9+ y8++ y9+ b9++
6×10^4
80.8 81.0 81.2 81.4 81.6 81.8
Fragment ions
Retention time (min)

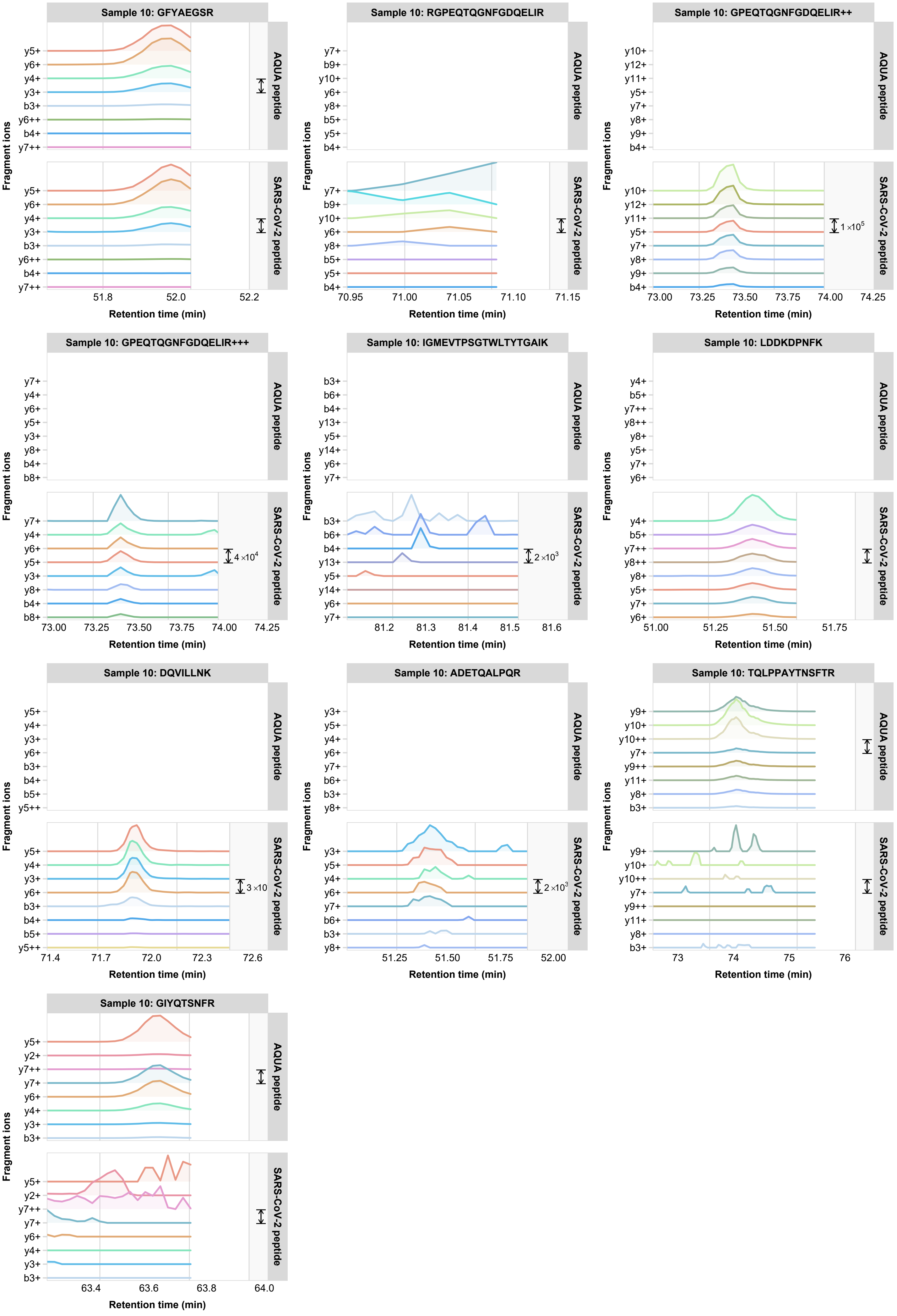
Sample 10: GFYAEGSR
y5+ y6+ y4+ y3+ b3+ y6++ b4+ y7++
51.8 52.0 52.2
Sample 10: RGPEQTQGNFGDQELIR
y7+ b9+ y10+ y6+ y8+ b5+ y5+ b4+
70.95 71.00 71.05 71.10 71.15
Sample 10: GPEQTQGNFGDQELIR++
y10+ y12+ y11+ y5+ y7+ y8+ y9+ b4+
1 ×10^5
73.00 73.25 73.50 73.75 74.00 74.25
Sample 10: GPEQTQGNFGDQELIR+++
y7+ y4+ y6+ y5+ y3+ y8+ b4+ b8+
4 ×10^4
73.00 73.25 73.50 73.75 74.00 74.25
Sample 10: IGMEVTPSGTWLTYTGAIK
b3+ b6+ b4+ y13+ y5+ y14+ y6+ y7+
2 ×10^3
81.2 81.3 81.4 81.5 81.6
Sample 10: LDDKDPNFK
y4+ b5+ y7++ y8++ y8+ y5+ y7+ y6+
51.00 51.25 51.50 51.75
Sample 10: DQVILLNK
y5+ y4+ y3+ y6+ b3+ b4+ b5+ y5++
3 ×10
71.4 71.7 72.0 72.3 72.6
Sample 10: ADETQALPQR
y3+ y5+ y4+ y6+ y7+ b6+ b3+ y8+
2 ×10^3
51.25 51.50 51.75 52.00
Sample 10: TQLPPAYTNSFTR
y9+ y10+ y10++ y7+ y9++ y11+ y8+ b3+
73 74 75 76
Sample 10: GIYQTSNFR
y5+ y2+ y7++ y7+ y6+ y4+ y3+ b3+
63.4 63.6 63.8 64.0
AQUA peptide
SARS-CoV-2 peptide
Fragment ions
Retention time (min)

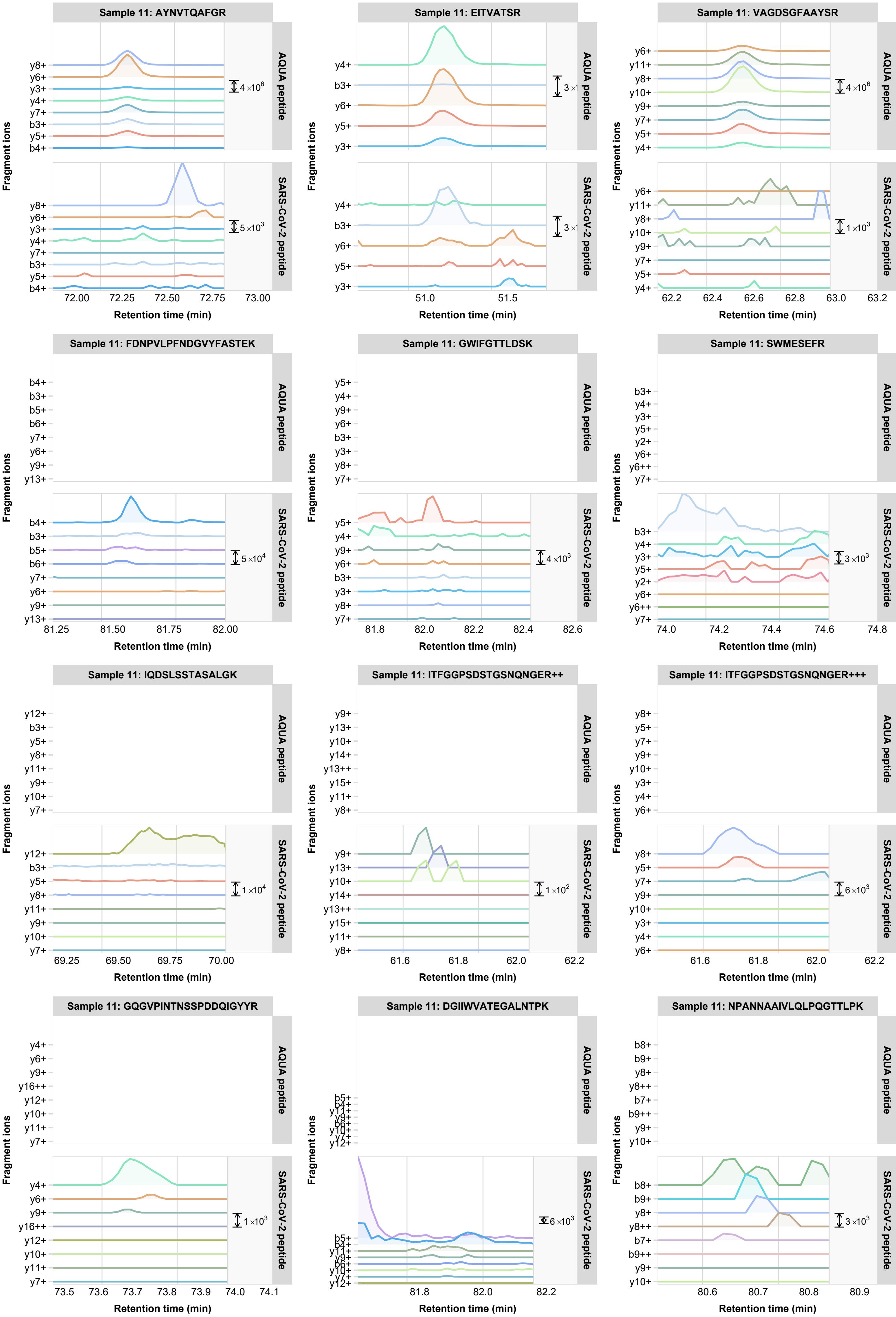

Sample 11: AYNVTQAFGR
Sample 11: EITVATSR
Sample 11: VAGDSGFAAYSR
Sample 11: FDNPVLPFNDGVYFASTEK
Sample 11: GWIFGTTLDSK
Sample 11: SWMESEFR
Sample 11: IQDSLSSTASALGK
Sample 11: ITFGGPSDSTGSNQNGER++
Sample 11: ITFGGPSDSTGSNQNGER+++
Sample 11: GQGVPINTNSSPDDQIGYYR
Sample 11: DGIIWVATEGALNTPK
Sample 11: NPANNAAIVLQLPQGTTLPK
AQUA peptide
SARS-CoV-2 peptide
Fragment ions
Retention time (min)
y8+ y6+ y3+ y4+ y7+ b3+ y5+ b4+
4×10^6
5×10^3
72.00 72.25 72.50 72.75 73.00
y4+ b3+ y6+ y5+ y3+
3×
51.0 51.5
y6+ y11+ y8+ y10+ y9+ y7+ y5+ y4+
4×10^6
1×10^3
62.2 62.4 62.6 62.8 63.0 63.2
b4+ b3+ b5+ b6+ y7+ y6+ y9+ y13+
5×10^4
81.25 81.50 81.75 82.00
y5+ y4+ y9+ y6+ b3+ y3+ y8+ y7+
4×10^3
81.8 82.0 82.2 82.4 82.6
b3+ y4+ y3+ y5+ y2+ y6+ y6++ y7+
3×10^3
74.0 74.2 74.4 74.6 74.8
y12+ b3+ y5+ y8+ y11+ y9+ y10+ y7+
1×10^4
69.25 69.50 69.75 70.00
y9+ y13+ y10+ y14+ y13++ y15+ y11+ y8+
1×10^2
61.6 61.8 62.0 62.2
y8+ y5+ y7+ y9+ y10+ y3+ y4+ y6+
6×10^3
61.6 61.8 62.0 62.2
y4+ y6+ y9+ y16++ y12+ y10+ y11+ y7+
1×10^3
73.5 73.6 73.7 73.8 73.9 74.0 74.1
b5+ b4+ y11+ y9+ b6+ y10+ y7+ y12+
6×10^3
81.8 82.0 82.2
b8+ b9+ y8+ y8++ b7+ b9++ y9+ y10+
3×10^3
80.6 80.7 80.8 80.9

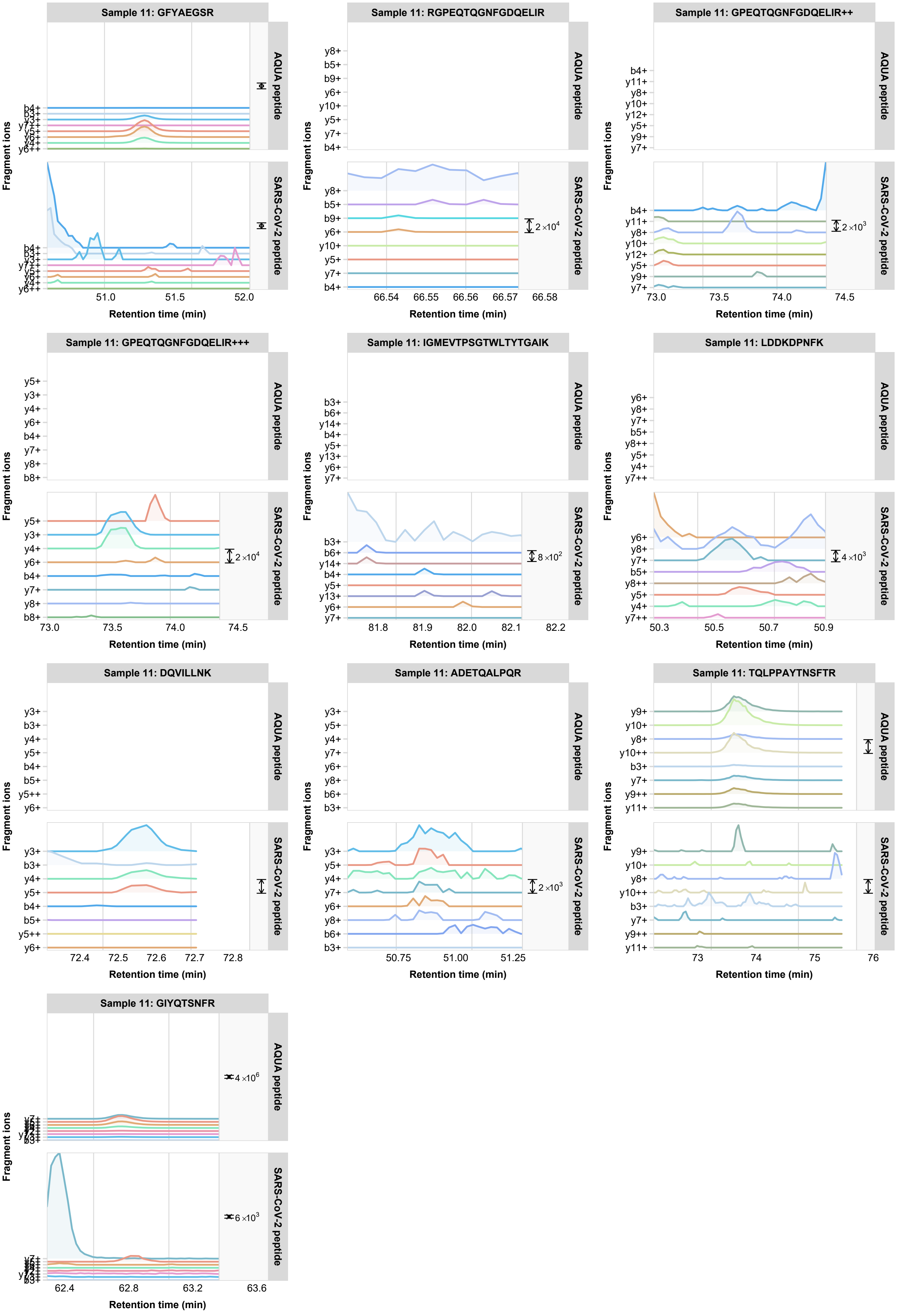

Sample 11: GFYAEGSR
AQUA peptide
SARS-CoV-2 peptide
b4+
b3+
y3+
y7++
y5+
y6+
y4+
y6++
51.0
51.5
52.0
Sample 11: RGPEQTQGNFGDQELIR
y8+
b5+
b9+
y6+
y10+
y5+
y7+
b4+
2×10^4
66.54
66.55
66.56
66.57
66.58
Sample 11: GPEQTQGNFGDQELIR++
b4+
y11+
y8+
y10+
y12+
y5+
y9+
y7+
2×10^3
73.0
73.5
74.0
74.5
Sample 11: GPEQTQGNFGDQELIR+++
y5+
y3+
y4+
y6+
b4+
y7+
y8+
b8+
2×10^4
73.0
73.5
74.0
74.5
Sample 11: IGMEVTPSGTWLTYTGAIK
b3+
b6+
y14+
b4+
y5+
y13+
y6+
y7+
8×10^2
81.8
81.9
82.0
82.1
82.2
Sample 11: LDDKDPNFK
y6+
y8+
y7+
b5+
y8++
y5+
y4+
y7++
4×10^3
50.3
50.5
50.7
50.9
Sample 11: DQVILLNK
y3+
b3+
y4+
y5+
b4+
b5+
y5++
y6+
72.4
72.5
72.6
72.7
72.8
Sample 11: ADETQALPQR
y3+
y5+
y4+
y7+
y6+
y8+
b6+
b3+
2×10^3
50.75
51.00
51.25
Sample 11: TQLPPAYTNSFTR
y9+
y10+
y8+
y10++
b3+
y7+
y9++
y11+
73
74
75
76
Sample 11: GIYQTSNFR
4×10^6
6×10^3
62.4
62.8
63.2
63.6
Fragment ions
Retention time (min)

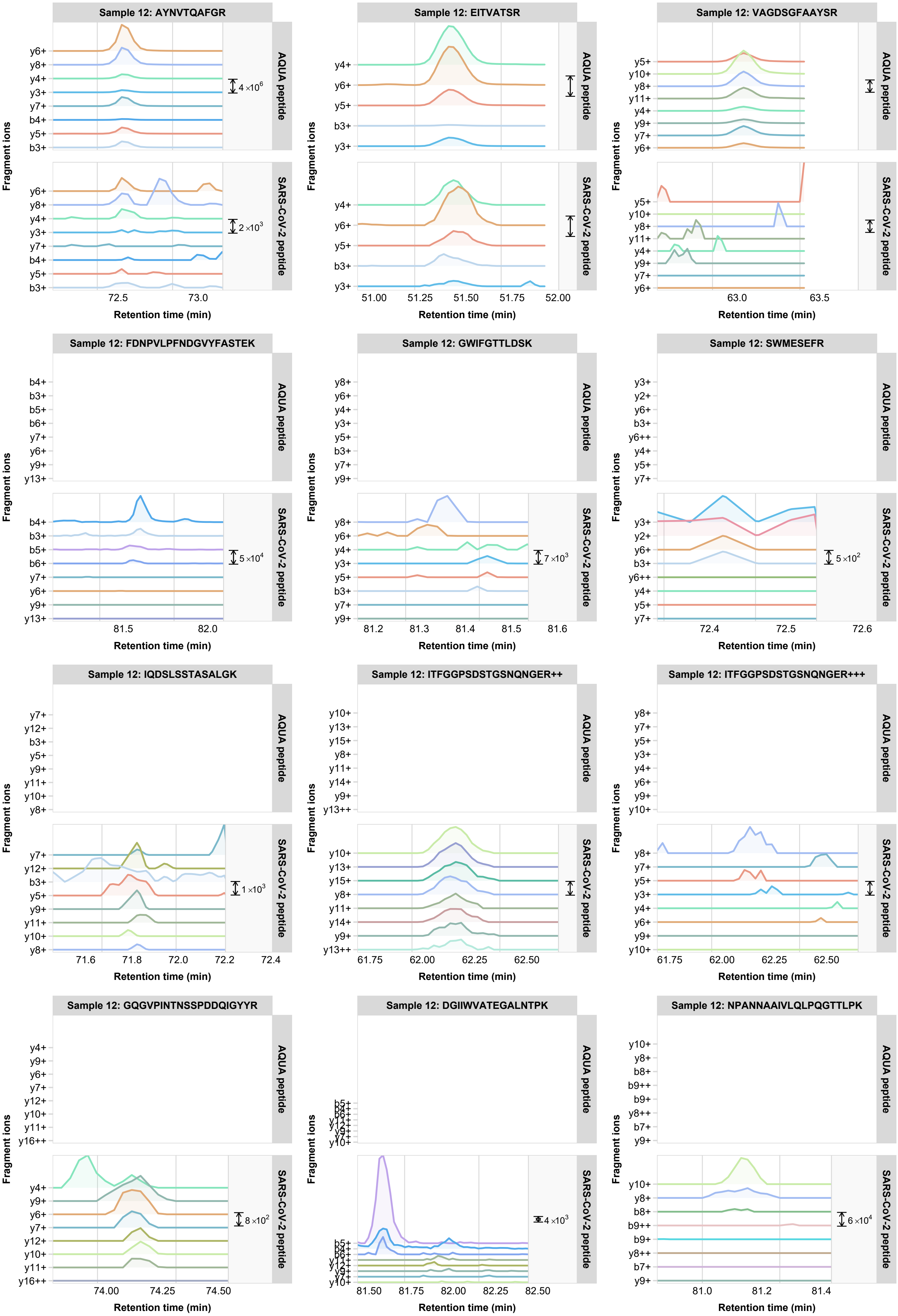

Sample 12: AYNVTQAFGR
y6+ y8+ y4+ y3+ y7+ b4+ y5+ b3+
4×10^6
2×10^3
72.5
73.0
Sample 12: EITVATSR
y4+ y6+ y5+ b3+ y3+
51.00 51.25 51.50 51.75 52.00
Sample 12: VAGDSGFAAYSR
y5+ y10+ y8+ y11+ y4+ y9+ y7+ y6+
63.0 63.5
Sample 12: FDNPVLPFNDGVYFASTEK
b4+ b3+ b5+ b6+ y7+ y6+ y9+ y13+
5×10^4
81.5 82.0
Sample 12: GWIFGTTLDSK
y8+ y6+ y4+ y3+ y5+ b3+ y7+ y9+
7×10^3
81.2 81.3 81.4 81.5 81.6
Sample 12: SWMESEFR
y3+ y2+ y6+ b3+ y6++ y4+ y5+ y7+
5×10^2
72.4 72.5 72.6
Sample 12: IQDSLSSTASALGK
y7+ y12+ b3+ y5+ y9+ y11+ y10+ y8+
1×10^3
71.6 71.8 72.0 72.2 72.4
Sample 12: ITFGGPSDSTGSNQNGER++
y10+ y13+ y15+ y8+ y11+ y14+ y9+ y13++
61.75 62.00 62.25 62.50
Sample 12: ITFGGPSDSTGSNQNGER+++
y8+ y7+ y5+ y3+ y4+ y6+ y9+ y10+
61.75 62.00 62.25 62.50
Sample 12: GQGVPINTNSSPDDQIGYYR
y4+ y9+ y6+ y7+ y12+ y10+ y11+ y16++
8×10^2
74.00 74.25 74.50
Sample 12: DGIIWVATEGALNTPK
b5+ b4+ b6+ y11+ y12+ y9+ y7+ y10+
4×10^3
81.50 81.75 82.00 82.25 82.50
Sample 12: NPANNAAIVLQLPQGTTLPK
y10+ y8+ b8+ b9++ b9+ y8++ b7+ y9+
6×10^4
81.0 81.2 81.4
AQUA peptide
SARS-CoV-2 peptide
Fragment ions
Retention time (min)

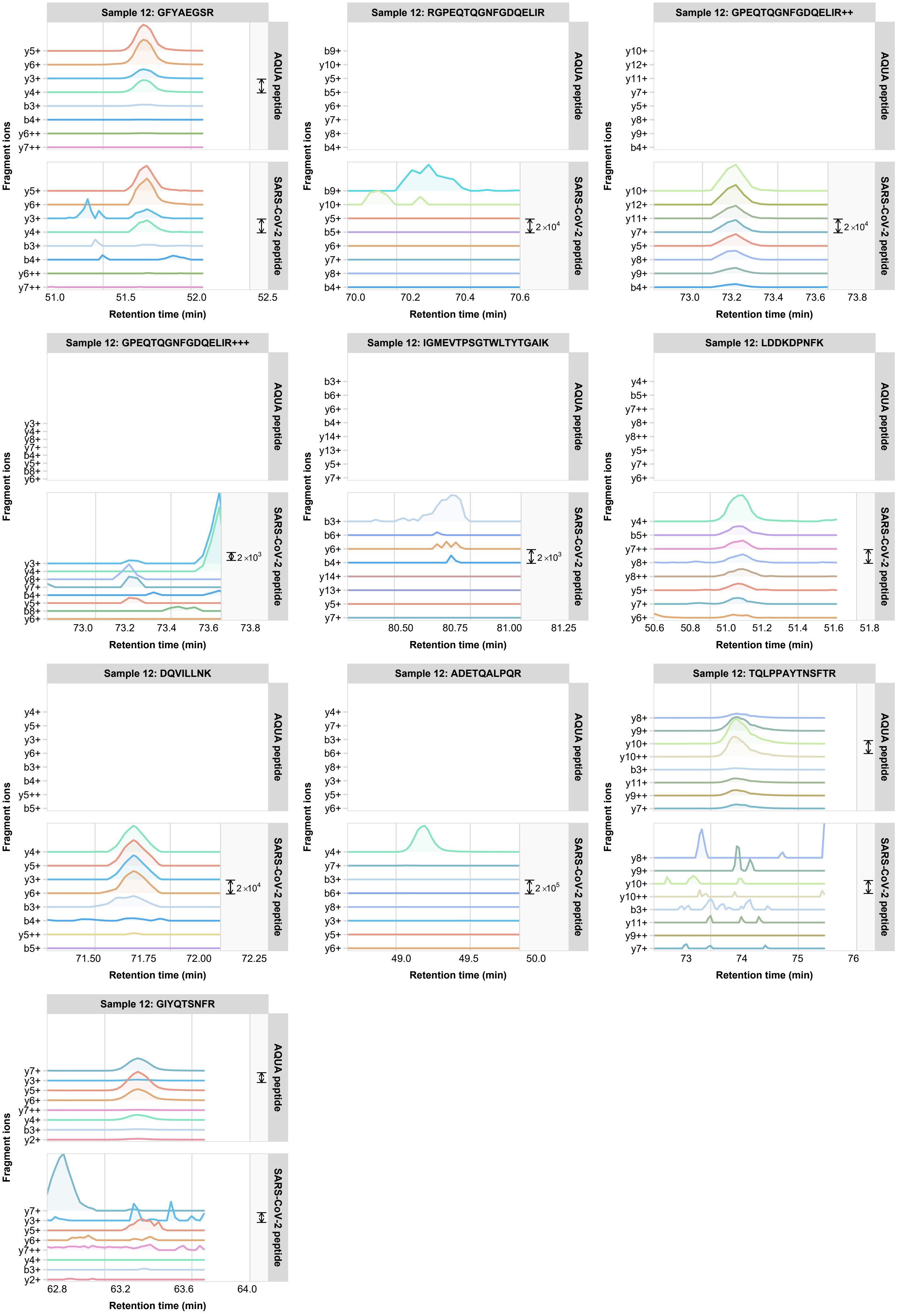
Sample 12: GFYAEGSR
y5+
y6+
y3+
y4+
b3+
b4+
y6++
y7++
AQUA peptide
SARS-CoV-2 peptide
Fragment ions
51.0
51.5
52.0
52.5
Retention time (min)
Sample 12: RGPEQTQGNFGDQELIR
b9+
y10+
y5+
b5+
y6+
y7+
y8+
b4+
2 ×10⁴
70.0
70.2
70.4
70.6
Sample 12: GPEQTQGNFGDQELIR++
y10+
y12+
y11+
y7+
y5+
y8+
y9+
b4+
2 ×10⁴
73.0
73.2
73.4
73.6
73.8
Sample 12: GPEQTQGNFGDQELIR+++
y3+
y4+
y8+
y7+
b4+
y5+
b8+
y6+
2 ×10³
73.0
73.2
73.4
73.6
73.8
Sample 12: IGMEVTPSGTWLTYTGAIK
b3+
b6+
y6+
b4+
y14+
y13+
y5+
y7+
2 ×10³
80.50
80.75
81.00
81.25
Sample 12: LDDKDPNFK
y4+
b5+
y7++
y8+
y8++
y5+
y7+
y6+
50.6
50.8
51.0
51.2
51.4
51.6
51.8
Sample 12: DQVILLNK
y4+
y5+
y3+
y6+
b3+
b4+
y5++
b5+
2 ×10⁴
71.50
71.75
72.00
72.25
Sample 12: ADETQALPQR
y4+
y7+
b3+
b6+
y8+
y3+
y5+
y6+
2 ×10⁵
49.0
49.5
50.0
Sample 12: TQLPPAYTNSFTR
y8+
y9+
y10+
y10++
b3+
y11+
y9++
y7+
73
74
75
76
Sample 12: GIYQTSNFR
y7+
y3+
y5+
y6+
y7++
y4+
b3+
y2+
62.8
63.2
63.6
64.0

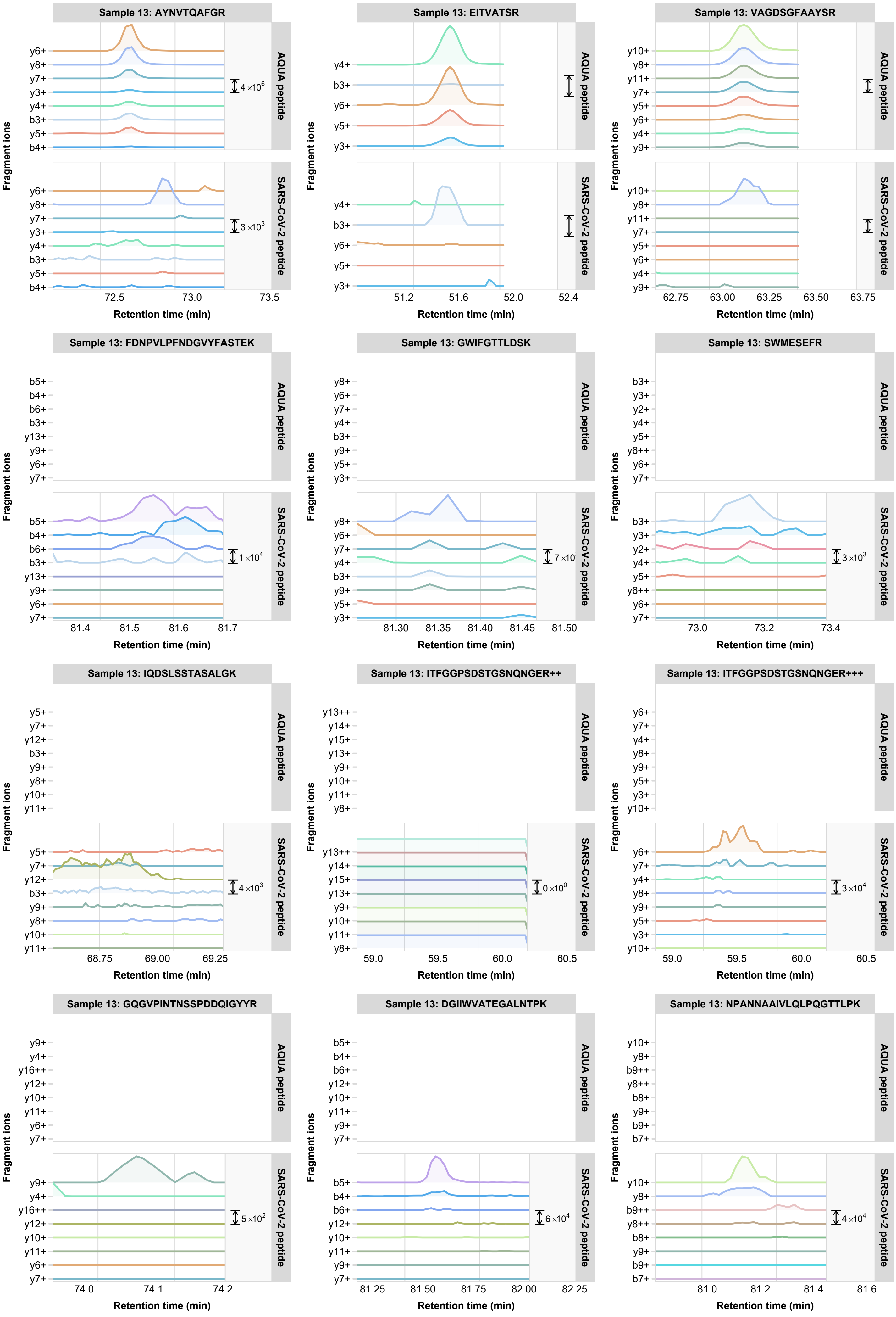
Sample 13: AYNVTQAFGR
AQUA peptide
SARS-CoV-2 peptide
y6+ y8+ y7+ y3+ y4+ b3+ y5+ b4+
4 ×10^6
3 ×10^3
72.5 73.0 73.5
Retention time (min)
Fragment ions
Sample 13: EITVATSR
y4+ b3+ y6+ y5+ y3+
51.2 51.6 52.0 52.4
Sample 13: VAGDSGFAAYSR
y10+ y8+ y11+ y7+ y5+ y6+ y4+ y9+
62.75 63.00 63.25 63.50 63.75
Sample 13: FDNPVLPFNDGVYFASTEK
b5+ b4+ b6+ b3+ y13+ y9+ y6+ y7+
1 ×10^4
81.4 81.5 81.6 81.7
Sample 13: GWIFGTTLDSK
y8+ y6+ y7+ y4+ b3+ y9+ y5+ y3+
7 ×10
81.30 81.35 81.40 81.45 81.50
Sample 13: SWMESEFR
b3+ y3+ y2+ y4+ y5+ y6++ y6+ y7+
3 ×10^3
73.0 73.2 73.4
Sample 13: IQDSLSSTASALGK
y5+ y7+ y12+ b3+ y9+ y8+ y10+ y11+
4 ×10^3
68.75 69.00 69.25
Sample 13: ITFGGPSDSTGSNQNGER++
y13++ y14+ y15+ y13+ y9+ y10+ y11+ y8+
0 ×10^0
59.0 59.5 60.0 60.5
Sample 13: ITFGGPSDSTGSNQNGER+++
y6+ y7+ y4+ y8+ y9+ y5+ y3+ y10+
3 ×10^4
59.0 59.5 60.0 60.5
Sample 13: GQGVPINTNSSPDDQIGYYR
y9+ y4+ y16++ y12+ y10+ y11+ y6+ y7+
5 ×10^2
74.0 74.1 74.2
Sample 13: DGIIWVATEGALNTPK
b5+ b4+ b6+ y12+ y10+ y11+ y9+ y7+
6 ×10^4
81.25 81.50 81.75 82.00 82.25
Sample 13: NPANNAAIVLQLPQGTTLPK
y10+ y8+ b9++ y8++ b8+ y9+ b9+ b7+
4 ×10^4
81.0 81.2 81.4 81.6

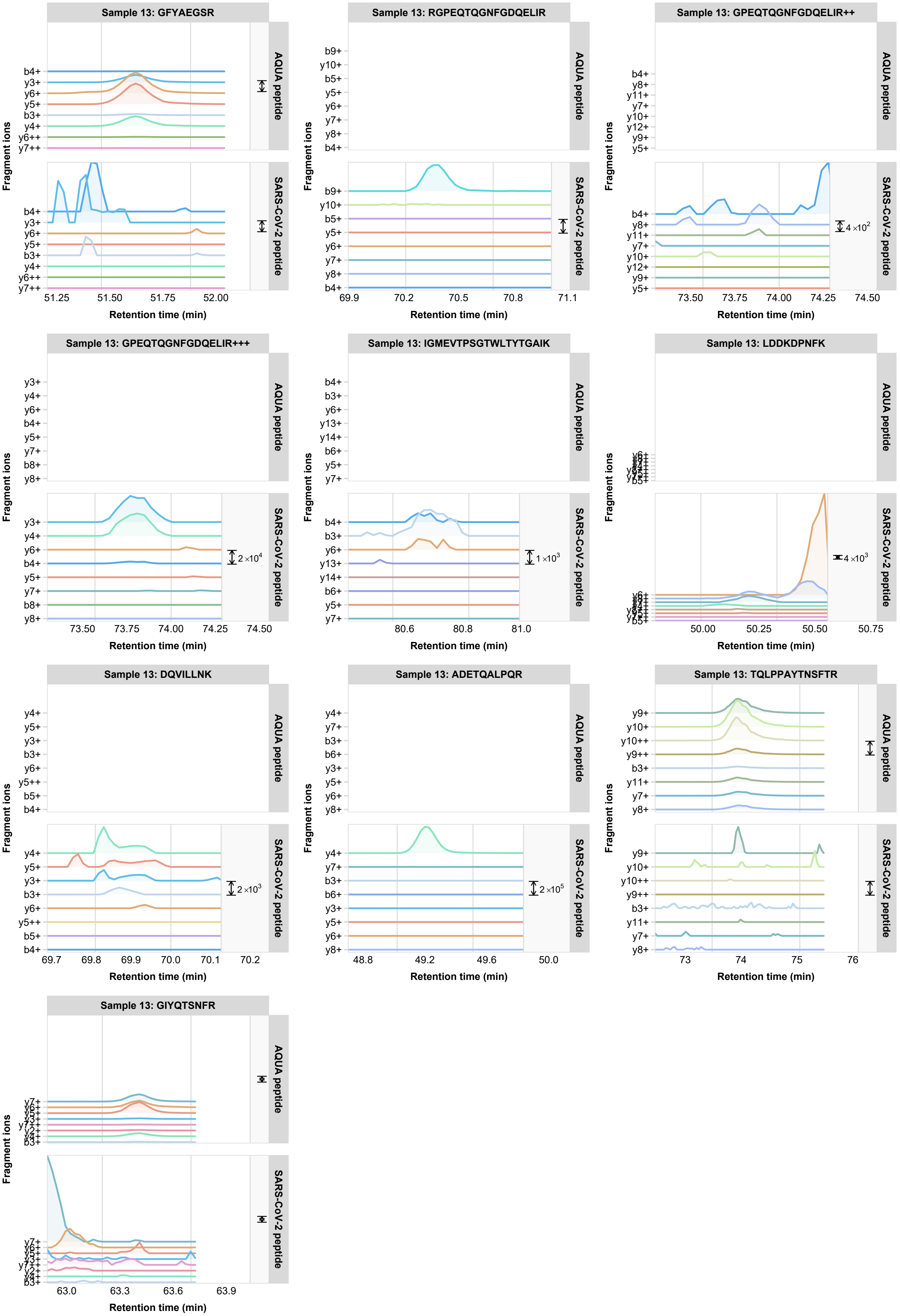
Sample 13: GFYAEGSR
AQUA peptide
SARS-CoV-2 peptide
b4+
y3+
y6+
y5+
b3+
y4+
y6++
y7++
Fragment ions
51.25
51.50
51.75
52.00
Retention time (min)
Sample 13: RGPEQTQGNFGDQELIR
b9+
y10+
b5+
y5+
y6+
y7+
y8+
b4+
69.9
70.2
70.5
70.8
71.1
Sample 13: GPEQTQGNFGDQELIR++
b4+
y8+
y11+
y7+
y10+
y12+
y9+
y5+
4 ×10²
73.50
73.75
74.00
74.25
74.50
Sample 13: GPEQTQGNFGDQELIR+++
y3+
y4+
y6+
b4+
y5+
y7+
b8+
y8+
2 ×10⁴
Sample 13: IGMEVTPSGTWLTYTGAIK
b4+
b3+
y6+
y13+
y14+
b6+
y5+
y7+
1 ×10³
80.6
80.8
81.0
Sample 13: LDDKDPNFK
4 ×10³
50.00
50.25
50.50
50.75
Sample 13: DQVILLNK
y4+
y5+
y3+
b3+
y6+
y5++
b5+
b4+
2 ×10³
69.7
69.8
69.9
70.0
70.1
70.2
Sample 13: ADETQALPQR
y4+
y7+
b3+
b6+
y3+
y5+
y6+
y8+
2 ×10⁵
48.8
49.2
49.6
50.0
Sample 13: TQLPPAYTNSFTR
y9+
y10+
y10++
y9++
b3+
y11+
y7+
y8+
73
74
75
76
Sample 13: GIYQTSNFR
63.0
63.3
63.6
63.9

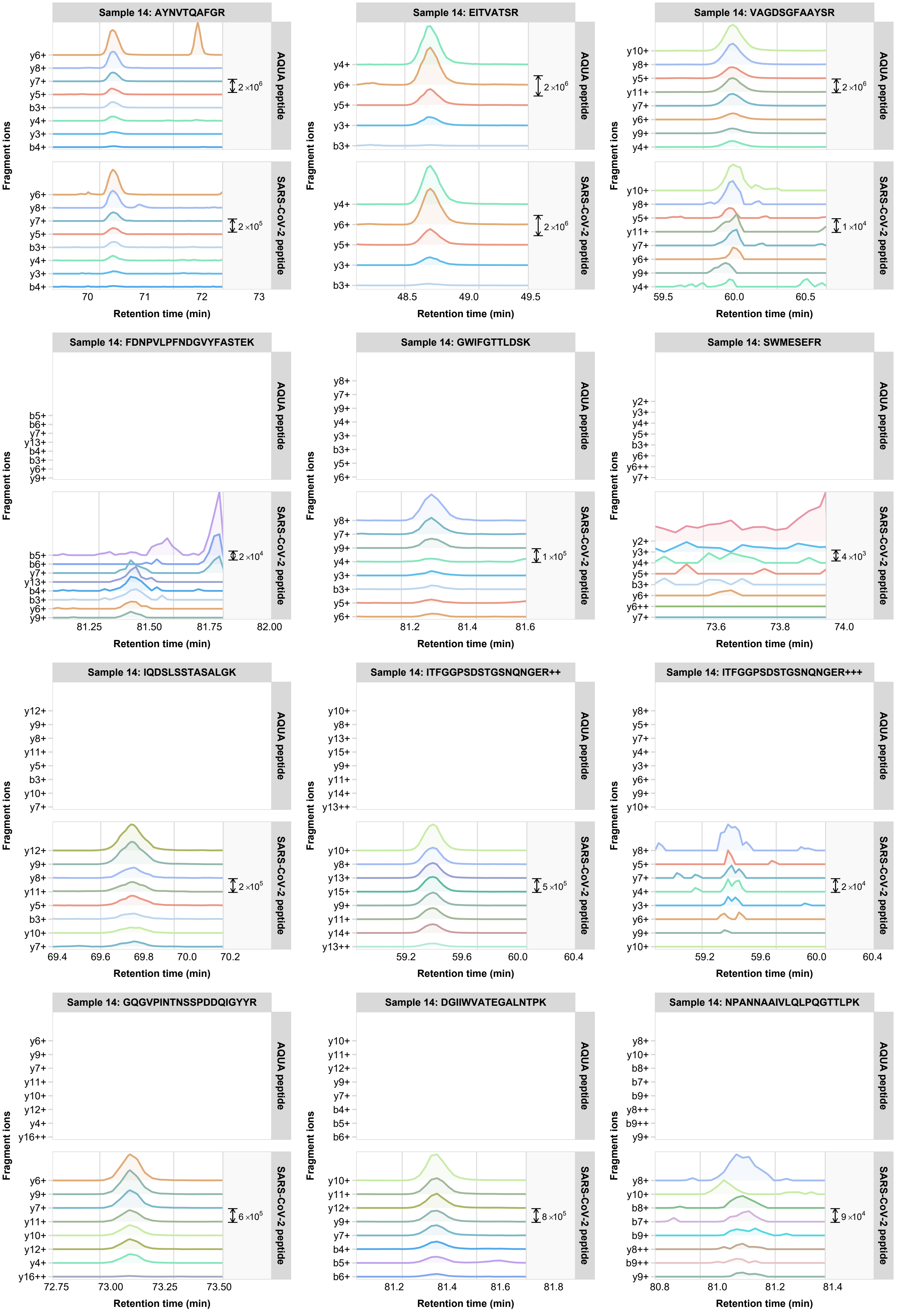

Sample 14: AYNVTQAFGR
AQUA peptide
SARS-CoV-2 peptide
y6+ y8+ y7+ y5+ b3+ y4+ y3+ b4+
2×10^6
2×10^5
70 71 72 73
Sample 14: EITVATSR
y4+ y6+ y5+ y3+ b3+
2×10^6
2×10^6
48.5 49.0 49.5
Sample 14: VAGDSGFAAYSR
y10+ y8+ y5+ y11+ y7+ y6+ y9+ y4+
2×10^6
1×10^4
59.5 60.0 60.5
Sample 14: FDNPVLPFNDGVYFASTEK
b5+ b6+ y7+ y13+ b4+ b3+ y6+ y9+
2×10^4
81.25 81.50 81.75 82.00
Sample 14: GWIFGTTLDSK
y8+ y7+ y9+ y4+ y3+ b3+ y5+ y6+
1×10^5
81.2 81.4 81.6
Sample 14: SWMESEFR
y2+ y3+ y4+ y5+ b3+ y6+ y6++ y7+
4×10^3
73.6 73.8 74.0
Sample 14: IQDSLSSTASALGK
y12+ y9+ y8+ y11+ y5+ b3+ y10+ y7+
2×10^5
69.4 69.6 69.8 70.0 70.2
Sample 14: ITFGGPSDSTGSNQNGER++
y10+ y8+ y13+ y15+ y9+ y11+ y14+ y13++
5×10^5
59.2 59.6 60.0 60.4
Sample 14: ITFGGPSDSTGSNQNGER+++
y8+ y5+ y7+ y4+ y3+ y6+ y9+ y10+
2×10^4
59.2 59.6 60.0 60.4
Sample 14: GQGVPINTNSSPDDQIGYYR
y6+ y9+ y7+ y11+ y10+ y12+ y4+ y16++
6×10^5
72.75 73.00 73.25 73.50
Sample 14: DGIIWVATEGALNTPK
y10+ y11+ y12+ y9+ y7+ b4+ b5+ b6+
8×10^5
81.2 81.4 81.6 81.8
Sample 14: NPANNAAIVLQLPQGTTLPK
y8+ y10+ b8+ b7+ b9+ y8++ b9++ y9+
9×10^4
80.8 81.0 81.2 81.4
Fragment ions
Retention time (min)

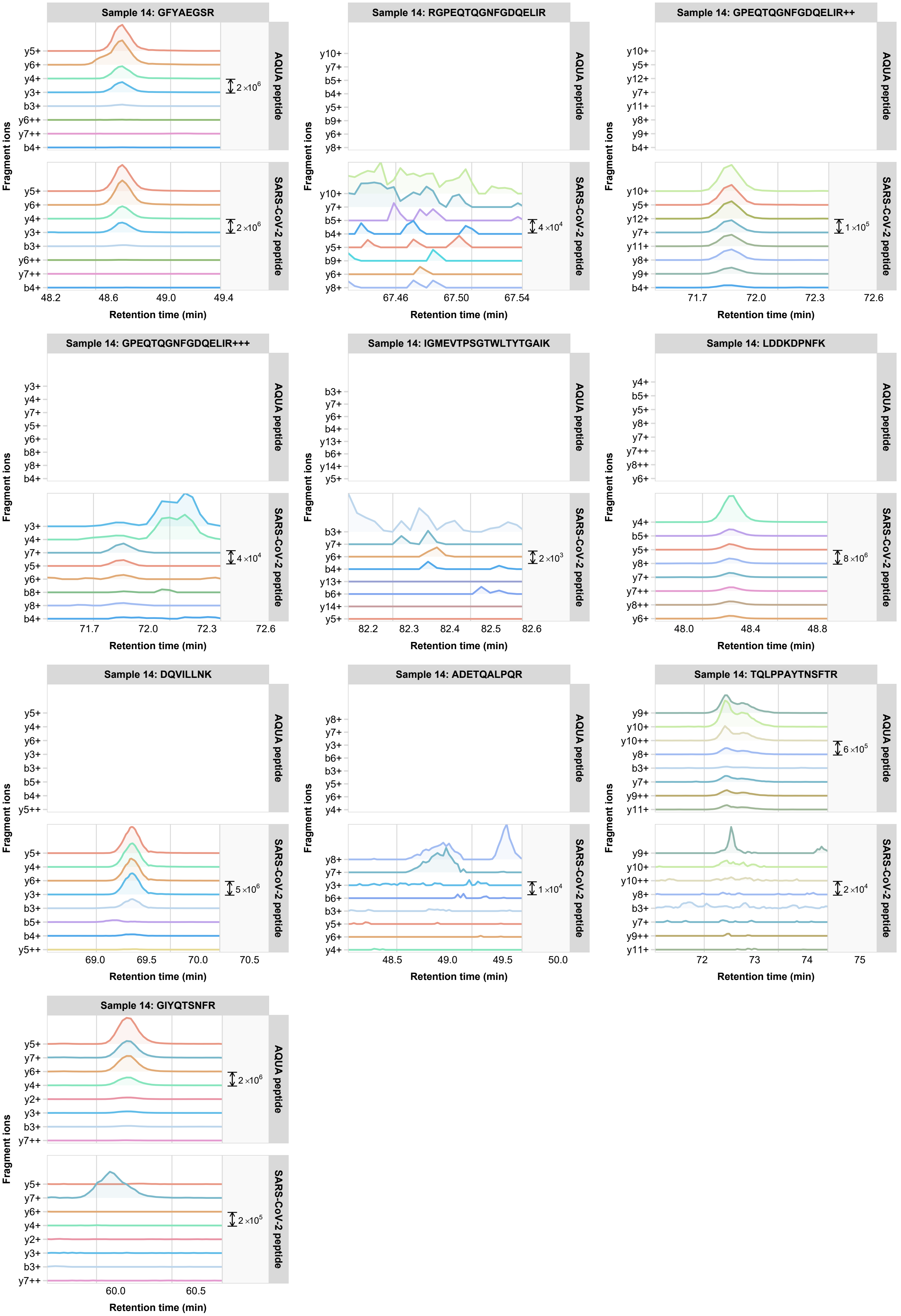
Sample 14: GFYAEGSR
Sample 14: RGPEQTQGNFGDQELIR
Sample 14: GPEQTQGNFGDQELIR++
Sample 14: GPEQTQGNFGDQELIR+++
Sample 14: IGMEVTPSGTWLTYTGAIK
Sample 14: LDDKDPNFK
Sample 14: DQVILLNK
Sample 14: ADETQALPQR
Sample 14: TQLPPAYTNSFTR
Sample 14: GIYQTSNFR
AQUA peptide
SARS-CoV-2 peptide
Fragment ions
Retention time (min)
2 ×10^6
4 ×10^4
1 ×10^5
2 ×10^3
8 ×10^6
5 ×10^6
1 ×10^4
6 ×10^5
2 ×10^4
2 ×10^5

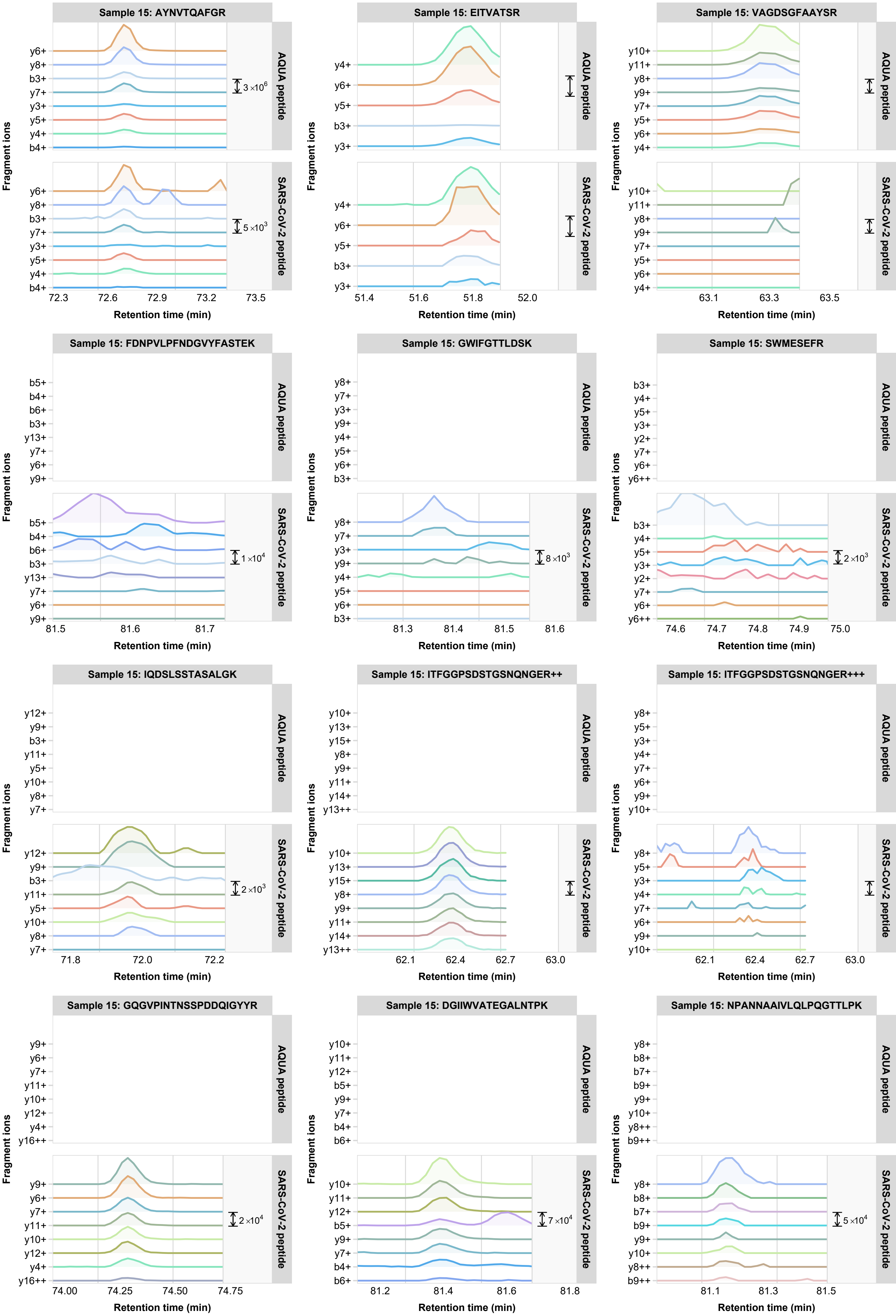
Sample 15: AYNVTQAFGR
AQUA peptide
SARS-CoV-2 peptide
y6+ y8+ b3+ y7+ y3+ y5+ y4+ b4+
3 ×10^6
5 ×10^3
72.3 72.6 72.9 73.2 73.5
Sample 15: EITVATSR
y4+ y6+ y5+ b3+ y3+
51.4 51.6 51.8 52.0
Sample 15: VAGDSGFAAYSR
y10+ y11+ y8+ y9+ y7+ y5+ y6+ y4+
63.1 63.3 63.5
Sample 15: FDNPVLPFNDGVYFASTEK
b5+ b4+ b6+ b3+ y13+ y7+ y6+ y9+
1 ×10^4
81.5 81.6 81.7
Sample 15: GWIFGTTLDSK
y8+ y7+ y3+ y9+ y4+ y5+ y6+ b3+
8 ×10^3
81.3 81.4 81.5 81.6
Sample 15: SWMESEFR
b3+ y4+ y5+ y3+ y2+ y7+ y6+ y6++
2 ×10^3
74.6 74.7 74.8 74.9 75.0
Sample 15: IQDSLSSTASALGK
y12+ y9+ b3+ y11+ y5+ y10+ y8+ y7+
2 ×10^3
71.8 72.0 72.2
Sample 15: ITFGGPSDSTGSNQNGER++
y10+ y13+ y15+ y8+ y9+ y11+ y14+ y13++
62.1 62.4 62.7 63.0
Sample 15: ITFGGPSDSTGSNQNGER+++
y8+ y5+ y3+ y4+ y7+ y6+ y9+ y10+
62.1 62.4 62.7 63.0
Sample 15: GQGVPINTNSSPDDQIGYYR
y9+ y6+ y7+ y11+ y10+ y12+ y4+ y16++
2 ×10^4
74.00 74.25 74.50 74.75
Sample 15: DGIIWVATEGALNTPK
y10+ y11+ y12+ b5+ y9+ y7+ b4+ b6+
7 ×10^4
81.2 81.4 81.6 81.8
Sample 15: NPANNAAIVLQLPQGTTLPK
y8+ b8+ b7+ b9+ y9+ y10+ y8++ b9++
5 ×10^4
81.1 81.3 81.5
Fragment ions
Retention time (min)

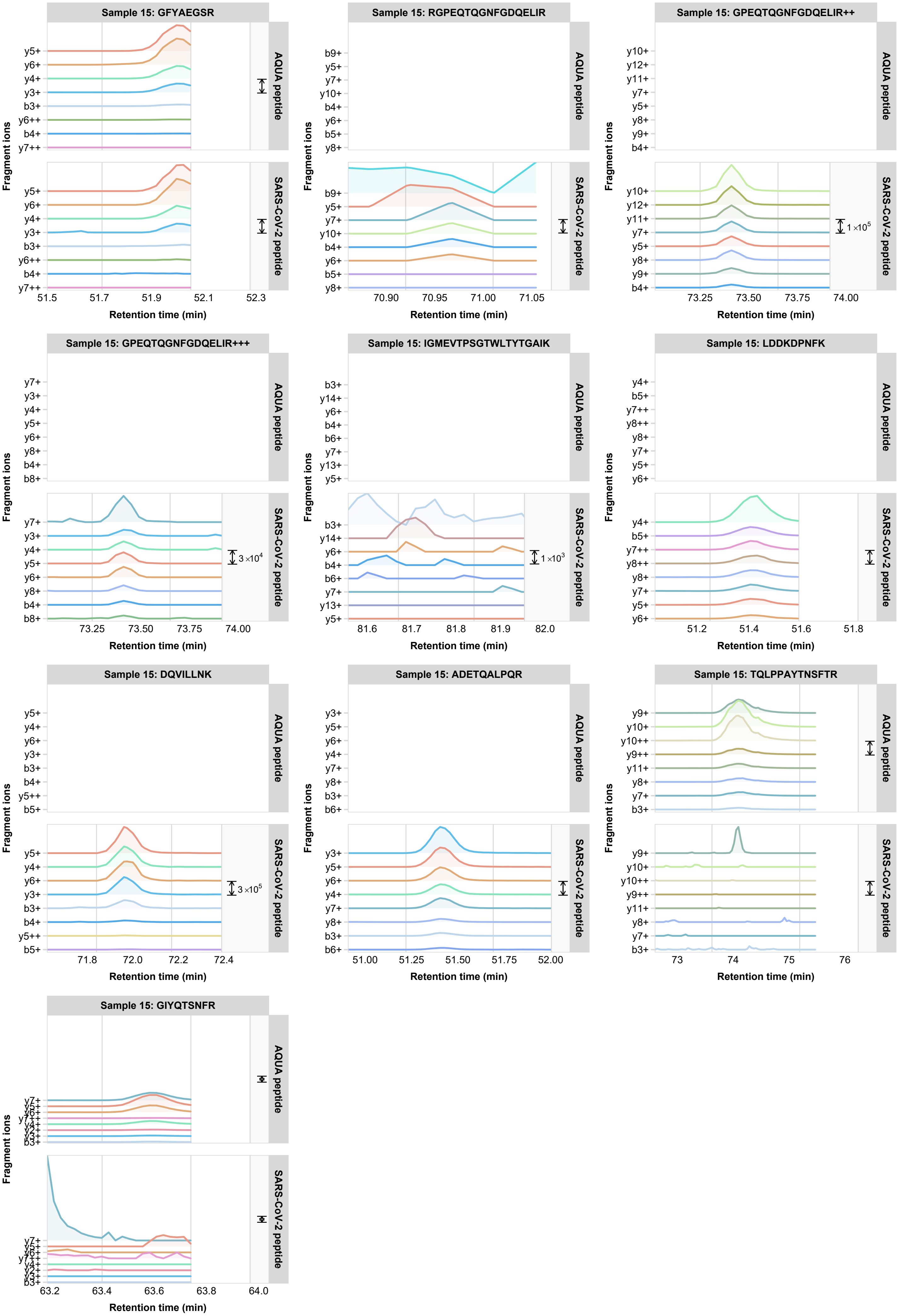
Sample 15: GFYAEGSR
Sample 15: RGPEQTQGNFGDQELIR
Sample 15: GPEQTQGNFGDQELIR++
Sample 15: GPEQTQGNFGDQELIR+++
Sample 15: IGMEVTPSGTWLTYTGAIK
Sample 15: LDDKDPNFK
Sample 15: DQVILLNK
Sample 15: ADETQALPQR
Sample 15: TQLPPAYTNSFTR
Sample 15: GIYQTSNFR
AQUA peptide
SARS-CoV-2 peptide
Fragment ions
Retention time (min)
1 ×10^5
3 ×10^4
1 ×10^3
3 ×10^5
